# Energy allocation explains how protozoan phenotypic traits change in response to temperature and resource supply

**DOI:** 10.1101/2023.07.25.550219

**Authors:** Andrea Perna, Enrico S. Rivoli, Julia Reiss, Daniel M. Perkins

## Abstract

To survive and reproduce, living organisms need to maintain an efficient balance between energy intake and energy expenditure. Changes in environmental conditions can disrupt previously efficient energy allocation strategies, and organisms are required to change their behaviour, physiology, or morphology to cope with the new environment. However, how multiple phenotypic traits interact with one another and with environmental conditions to shape energy allocation remains poorly understood. To better understand this type of phenotype-environment interactions, we develop a predictive framework, grounded in energetic and biophysical principles that allows us to make predictions on how metabolic rate and movement speed should change in response to environmental temperature and resource supply, differentiating between short-term, acute exposure to novel conditions and longer-term exposure that allows acclimation or adaptation. We tested these predictions by exposing axenic populations of the ciliate *Tetrahymena pyriformis* to different combinations of temperature and resource availability. We measured population growth, cell size, respiration, and movement. Acute increases in temperature led to higher movement speeds and respiration rates, consistent with expectations from physical scaling relationships such as the Boltzmann-Arrhenius equation and the viscous drag acting on movement. However, by around 3.5 days after the introduction of *Tetrahymena* into a novel environment, all measured traits shifted toward values closer to those of the original environment. These changes likely reflect phenotypic acclimation responses that restored a more efficient energy allocation under the new conditions. Changes in cell size played a key role in this process, by simultaneously affecting multiple phenotypic traits, including metabolic rate and the energetic costs of movement. In small microbial consumers like *Tetrahymena*, body size can change rapidly, relative to ecological and seasonal timescales. Changes in body size can therefore be effectively leveraged - alongside physiological and biochemical regulations – to cope with environmental changes.

**Open research:** The raw data and all the analysis code used for this work are publicly available on github (https://github.com/pernafrost/Tetrahymena), and they are also deposited in Dryad (https://doi.org/10.5061/dryad.2v6wwpzvv). By downloading and running the code, it is possible to reproduce all the figures presented in the manuscript and in the Appendix. Please refer to the linked repositories for additional information, and feel free to contact the authors in case you need additional guidance.

## Introduction

To survive and reproduce, living organisms need to obtain energy from their environment, and to allocate it efficiently to competing processes of growth, movement, and reproduction (Alexander, 1996). Different strategies of energy allocation manifest, in practice, as different phenotypes: having a larger or smaller body size, or a different growth rate and movement patterns, for example. When the environment changes, the change can affect the organisms directly (e.g. by increasing the metabolic rate when temperature rises) and indirectly (e.g. if resources become scarce). As a consequence, energy allocation strategies that used to be efficient in the original environment may become inefficient, and organisms are required to adapt to the new environment by changing their behaviour, or their phenotype, or they will incur a fitness cost.

An extensive body of scientific literature has identified general patterns of phenotypic variation in relation to environmental variables such as temperature and the availability of resources. These empirically observed patterns are described in terms of ‘rules’ of phenotypic change such as the *temperature-size rule* – the observation that ectotherms grow to a larger adult size when they develop in a colder environment (Atkinson and Sibly, 1997; Angilletta Jr et al., 2004; Walters and Hassall, 2006), or the relation between metabolic rate, environmental temperature and body size observed across different taxa (Gillooly et al., 2001; Brown et al., 2004). Phenotypic traits such as the movement and the behaviour of organisms are also modulated in consistent ways by environmental variables, with temperature, again, playing an important role (e.g. Gibert et al., 2016; Abram et al., 2017).

Understanding all these general patterns of phenotypic change is important because they not only affect the survival of individual organisms, but also ecological variables: population growth rate (DeLong and Hanson, 2009), carrying capacity (Bernhardt et al., 2018), and predation risk (Gibert et al., 2017), with cascading effects beyond a single species (Dell et al., 2014) to also affect community assembly (Saito et al., 2021) and macroscale patterns in the spatial distribution of species (e.g. Colombo et al., 2025).

The proximate causes of these ‘rules’ of phenotypic variation – namely, the physiological and developmental regulations involved – are becoming increasingly well understood (e.g. Verberk et al., 2021; Hermaniuk et al., 2021; Arroyo et al., 2022). This growing understanding allows us to disentangle the contributions of genetic and plastic factors that mediate phenotypic changes in response to different environments (Hairston Jr et al., 2005; Carroll et al., 2007; Plum et al., 2022; Vinton et al., 2022).

However, it is also important to understand the ultimate causes, and particularly the adaptive value, of commonly observed phenotypic changes in relation to the environment. This type of understanding can help make general predictions about the direction of phenotypic change: if a phenotypic shift in a particular direction brings an adaptive benefit – we could argue – this is the kind of shift that is more likely to be produced and to persist in a population on the long-term, whether mediated by plasticity or by evolution. Phenomena that appear to be maladaptive, like non-adaptive reaction norms could then be interpreted in terms of limited previous selective pressure when organisms are exposed to environmental conditions outside the range of those historically experienced (Ghalambor et al., 2007). Transient phenotypic changes that are different over short and long timescales (Dupont et al., 2024; DeLong et al., 2017) can also be interpreted in relation to adaptive value benchmarks, for instance in terms of mechanistic constraints that could prevent the implementation of plastic adaptive responses over short timescales, such as temporal lags for genetic expression and morphological changes.

Traditionally, insight into the question, which morphological traits and behavioural strategies are ‘optimal’ for a given set of environmental parameters, has been provided by optimisation theory. Optimisation theory has been used to make predictions about a number of different phenotypic traits, ranging from the size and shape of an organism (Cooper and Denny, 1997; West et al., 2001), to the arrangement of its constituent units, from molecules to organs (Banavar et al., 2002; Itzkovitz and Alon, 2007; Pérez-Escudero and de Polavieja, 2007; Pérez-Escudero et al., 2009), to movement (Gazzola et al., 2014; Labonte, 2023), foraging strategies (MacArthur and Pianka, 1966; Charnov, 1976; Pyke et al., 1977) and predator-prey interactions (Visser, 2011) – see (Parker and Smith, 1990; Alexander, 1996) for a review and additional references. Optimisation theory does not tell how a phenotype is produced from development or evolution (Gould et al., 1979), but it tells us which characteristics a phenotypic trait should have to maximize some proxy of fitness – food intake, survival rate, offspring production, etc. – thus providing a theoretical benchmark that can sometimes illuminate the interpretation of empirically observed phenomena.

In the present work, we adopt a related framework, which involves considering the energetic costs and benefits of different phenotypic and ‘behavioural’ traits explicitly, to study how a model microorganism, the ciliate *Tetrahymena pyriformis* (henceforth *Tetrahymena*) responds to changes in environmental conditions characterised by differences in temperature and concentration of food. We make predictions on the direction of change of phenotypic traits (i.e. whether they are expected to increase or decrease) for four main traits: rate of population increase, cell size (expressed as cell volume), respiration (as a proxy for metabolic rate), and movement speed. While the first three of these phenotypic traits have been extensively considered in the scientific literature on thermal adaptation, movement speed is a ‘behavioural’ trait that hasn’t yet systematically and quantitatively been put in relation with the other three.

We test these predictions empirically, by measuring how the phenotypic traits of *Tetrahymena* change under different culture conditions and different timescales. We interpret our results based on the idea that the physical parameters of the environment affect the organisms directly (for example, through the effect of temperature on metabolic rate) and in so doing they alter the energetic costs and benefits of different phenotypic traits and behaviours. For example, water viscosity affects the cost of movement, and the availability of feeding resources determines the energetic return of foraging. We formalise the link between movement, drag, and energetic costs relying on well-established low-Reynolds-number hydrodynamics formulas. We then attempt to understand the phenotypic changes of cell size, movement speed, population growth rate, and respiration that happen over longer timescales in terms of a simple ‘energy allocation equation’, whose balance between energy acquisition and energy expenditure needs to be restored in the new environment through the combined effect of multiple phenotypic changes.

Hence, the resulting framework provides a means to show how single-celled eukaryotes allocate energy to feeding, basal metabolism and reproduction and to estimate important parameters for these calculations (such as the values of activation energy for respiration and movement).

## Theoretical framework

The energy that individual organisms (here, individual *Tetrahymena* cells) can invest in reproduction and hence population growth results from a balance between energy intake from feeding *F*, and energy expenditure for maintenance (basal metabolism) *B*, and movement (propulsion) *P*:

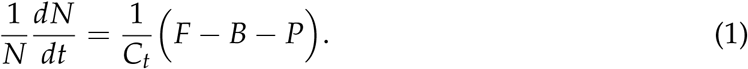

In the equation, the left-hand side represents the *per capita* growth rate of the population (*N* is the population size – the number of cells – and 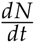 is the growth rate of the entire population), *F*, *B*, and *P* have units of power (e.g. nanoWatts) and *C_t_* can be thought of as the amount of energy required to form a new cell.

The dependence of the individual variables *F*, *B*, *P*, and *C_t_* on the characteristics of the organism or of the environment can be made explicit by progressively adding multiple levels of detail, as we do in the Appendix, sec. 3. Here, we mainly introduce eq. 1 to make predictions about qualitative patterns that we can observe experimentally, and it will be sufficient to remind the general forms of terms *F*, *B*, and *P*, so that readers can follow the logic without the burden of too many details, which remain available in the Appendix, sec. 3.

The term *B*, representing the basal metabolic rate, is expected to increase with increasing temperature *T* and also to increase with cell volume *V* (relative to a reference volume *V*_0_) following the equation of metabolic theory

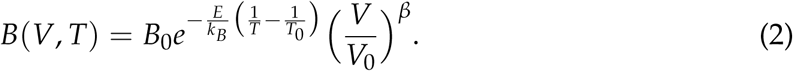

Here, *B*_0_ is the value of metabolic rate measured at the reference temperature *T*_0_. Essentially, metabolic rate increases with temperature following an Arrhenius equation characterised by the value of activation energy *E*, whereas *k_B_* is the Boltzmann constant, *T* is the environmental temperature (in Kelvin), and *T*_0_ is the reference temperature. The scaling of metabolism with volume *V* (here expressed as relative to a reference volume *V*_0_) follows an allometric relation with exponent *β* (Gillooly et al., 2001; Brown et al., 2004), where *β* ≈ 1 for unicellular eukaryotes (DeLong et al., 2010).

The term *P* represents swimming-related power dissipation. Swimming entails energetic costs, associated with overcoming the drag of water. For small organisms swimming at low Reynolds numbers, the power *P* required for swimming at speed *U* scales as the square of *U*

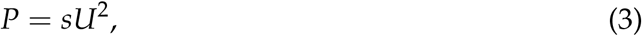

where *s* is a proportionality constant, increasing linearly with cell radius and also affected by cell shape and by water viscosity (which in turn can change with temperature). The detailed equations are presented in the Appendix, sec. 3, but readers can skip the full equations if they are willing to accept that *P* increases linearly with cell radius, and quadratically with speed *U*.

Finally, the term *F* represents feeding. We assume a Holling type II functional response in which the encounter rate with food is proportional not only to food density, but also to the volume of medium effectively processed per unit time. In a simple ‘direct interception’ framework, this volume scales with the movement speed of *Tetrahymena* (see e.g. Kiørboe (2011); Visser (2011) for analogous models). This leads to the following equation for the feeding rate, which explicitly depends on the swimming speed *U*:

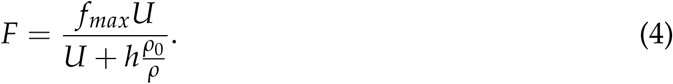

As before, *U* is the movement speed of *Tetrahymena*, while *h* is a ‘half saturation speed’ or the speed at which the feeding rate reaches half the maximum value, *f_max_*is the maximum feeding rate (which in our system might be limited by digestion time more than by prey handling). Finally, *ρ* is the concentration of resources in the environment, here expressed in relation to the concentration *ρ*_0_ of a standard medium (in our experiments, *ρ*_0_ is the concentration of the 100 % growth medium).

We choose a feeding response mediated by speed because organisms the size of *Tetrahymena* or larger can only feed efficiently by producing fluid currents relative to their body (Karp-Boss et al., 1996), and *Tetrahymena* achieves this primarily through self-motion, rather than by generating currents in the fluid through the action of feeding cilia around its oral apparatus (Liu, Costello and Kanso, 2025). At low Reynolds numbers, forces produced on the fluid are necessarily associated with equal and opposite forces on the microorganism, and scale proportionally to its movement (except for those microorganisms that attach to the substrate).

Ciliates can further modulate their feeding by generating ciliary currents that transport particles to the cytostome, rather than by direct interception through locomotion alone, so while swimming speed can be viewed as a proxy for the overall rate of fluid processing, some of the parameters of equation 4 could also reflect a shifting allocation of ciliary activity from somatic cilia to feeding cilia. While at low prey density, efficient feeding would likely require that movement and feeding remain broadly coupled (an increase of movement speed, leading to increased prey encounters, would be inefficient if the encountered prey is not consumed); at high prey density (*ρ* ≫ 1), however, when feeding becomes less dependent on movement speed *U*, it is possible that the activity of feeding cilia may remain high, contributing to sustain the maximum feeding rate *f_max_*.

Our model formulation should be interpreted as a minimal or ‘null’ model linking feeding to the rate at which cells interact with their surrounding medium, and we expect future work to refine this model to further link its parameters to the differential activity of oral and locomotory cilia.

Below, we analyse the model by asking questions such as: how should the swimming speed of *Tetrahymena* change with temperature if the energy that *Tetrahymena* allocated to movement was a fixed proportion of its total metabolic energy expenditure? And how should the swimming speed change as a function of temperature and resources if *Tetrahymena*, instead, optimized the proportion of energy allocated to movement in each environment? To structure the range of possible theoretical questions, we focus in particular on formulating three testable predictions and one theoretically-based consideration.

### Prediction 1: Exposure to high temperature and sufficient resources should lead to higher metabolic rate and faster population growth, unless compensated by other phenotypic changes

The fact that metabolic rate increases with temperature is a direct implication of the Arrhenius-type dependence of metabolic rate on environmental temperature (eq. 6).

As for population growth, we can reason as follows. When resources are abundant, feeding *F* will approach the maximum rate (*F* ≃ *f_max_*, eq. 4), and will mainly be limited by prey handling and digestion time (changes of movement speed *U*, and of the associated prey encounter rate, have a comparatively smaller impact on the actual feeding in these conditions of abundant resources). This means that under abundant resources, the high metabolic rate induced by the higher temperature is also likely to speed up the digestion process: both the *F* and the *B* term in equation 1 increase in similar proportion, and so also does their difference, resulting in faster *per capita* growth rate of the population.

The same reasoning does not apply when resources are scarce. Under these conditions, metabolic rate *B* is still expected to increase with temperature (consistent with the effect of temperature on chemical reactions described by the Arrhenius equation), but at low density of resources feeding *F* is limited by prey encounter, not by digestion time! This case will be considered in more detail below (in the ‘prediction 3’ subsection).

### Prediction 2: Cells exposed to a higher temperature over short timescales will increase their movement speed, but this effect will be described by a lower activation energy than for respiration

If cells do not modulate the proportion of metabolic energy that they allocate to movement, we would expect that, when their metabolic rate increases, for instance because of increased environmental temperature, their movement speed should also increase: a faster metabolism can power a faster movement. However, speed should increase less than metabolic rate with increasing temperature, i.e. speed should have a lower activation energy. In the case of a microscopic swimmer, such as *Tetrahymena*, swimming at low Reynolds number, the power *P* dissipated for moving in water at speed *U* increases proportionally to the square of speed (*P* ∝ *U*^2^, eq. 3), meaning that a 4-fold increase in the metabolic power invested into movement would lead only to a two-fold increase of swimming speed. This non-linear relation between power consumption and movement speed leads to a predicted activation energy for speed approximately 50 % the activation energy for respiration (this relation between metabolic power and movement speed can be calculated more precisely by also accounting for the effect of temperature on water viscosity – eq. 14 – and is illustrated in the Appendix, sec. 3.1).

Our second prediction is that *Tetrahymena* should exhibit such an increase of movement speed with increasing temperature, at least over short timescales, before genetic or metabolic regulations intervene to alter the proportion of metabolic energy allocated to movement relative to the other biological functions.

### Prediction 3: Short-term changes of movement speed and metabolic rate are unlikely to be sustainable in the long-term

The changes of metabolic rate and movement speed with environmental temperature predicted above reflect simple physico-chemical changes in the rate of metabolic reactions, and in the physical forces acting on the microorganism from the environment (such as the drag) but they are unlikely to reflect efficient energy allocation strategies for the fitness of the individual organism.

For example, an increase of movement speed at high temperature, and the associated increase of metabolic power *P* allocated to movement (as in eq. 1) is only beneficial if it is also matched by a corresponding increase in energy intake from feeding *F*. However, this is unlikely to be the case because the energetic gains from feeding are expected to increase more slowly than the energetic costs of movement. In simple encounter-based models, the volume searched for food – and thus the attack rate – scales at most linearly with speed (at best, *F* ∝ *U* in eq. 4), while the energetic cost of movement increases faster than speed (*P* ∝ *U*^2^).

This mismatch between feeding energy acquisition rate and power consumption of movement is robust to the details of model formulation: any mechanism for which feeding increases sub-quadratically with speed will lead to the same qualitative outcome. Predators typically only encounter new prey items in proportion to the speed at which they move, i.e. linearly with speed – much less than quadratically. Food encounter models that account for the diffusion of food molecules (or for prey movement, in case of living prey), predict that an increase of predator speed leads to an even less than proportional (i.e. sublinear) gain in food intake (Karp-Boss et al., 1996). Digestion time can further limit the uptake of resources independently of the rate at which they are encountered.

These ‘optimal foraging’ arguments can be developed quantitatively, as they amount to finding the movement speed *U* that maximizes the *per capita* growth rate for a given value of temperature and concentration of resources. The derivation is straightforward and involves maximizing eq. 1 with respect to speed, after substituting the speed-explicit expressions for *P* and *F* (eq. 3 and 4, respectively). Readers can find this derivation in the Appendix, sec. 3.1.

The optimal foraging speed satisfies

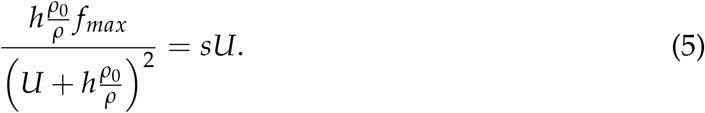

For example, the optimal speed calculated in this way for a cell of length *l* = 50 *µ*m, width *w* = 25 *µ*m, maximum feeding rate *f_max_* = 6 nW, ‘half saturation speed’ *h* = 250 *µ*m/s, and efficiency of conversion of metabolic energy to movement *η_c_* = 0.02 % follows the patterns shown in figure 1. In this model, the optimal speed remains relatively constant with temperature (left panel in fig. 1), with a slight increase of speed for increasing temperature entirely explained by the reduction of water viscosity with increasing temperature (swimming fast is less energetically costly at high temperature). The same model predicts a non-monotonic effect of growth medium concentration on optimal movement speed (right panel in fig. 1). Intuitively, the curve can be interpreted as follows: when nutrient resources in the growth medium are scarce, optimal speed is very low because the energy consumption associated with a fast movement is not compensated by feeding. Movement speed reaches a maximum for intermediate concentrations of the growth medium, to then decrease again. In fact, when food is abundant, a large amount of food available within a small volume, and feeding is limited by handling and digestion, rather than by food encounter rate: a higher speed does not bring additional benefits. (For another similar treatment of optimal swimming speed see also Visser (2007)).

**Figure 1:**
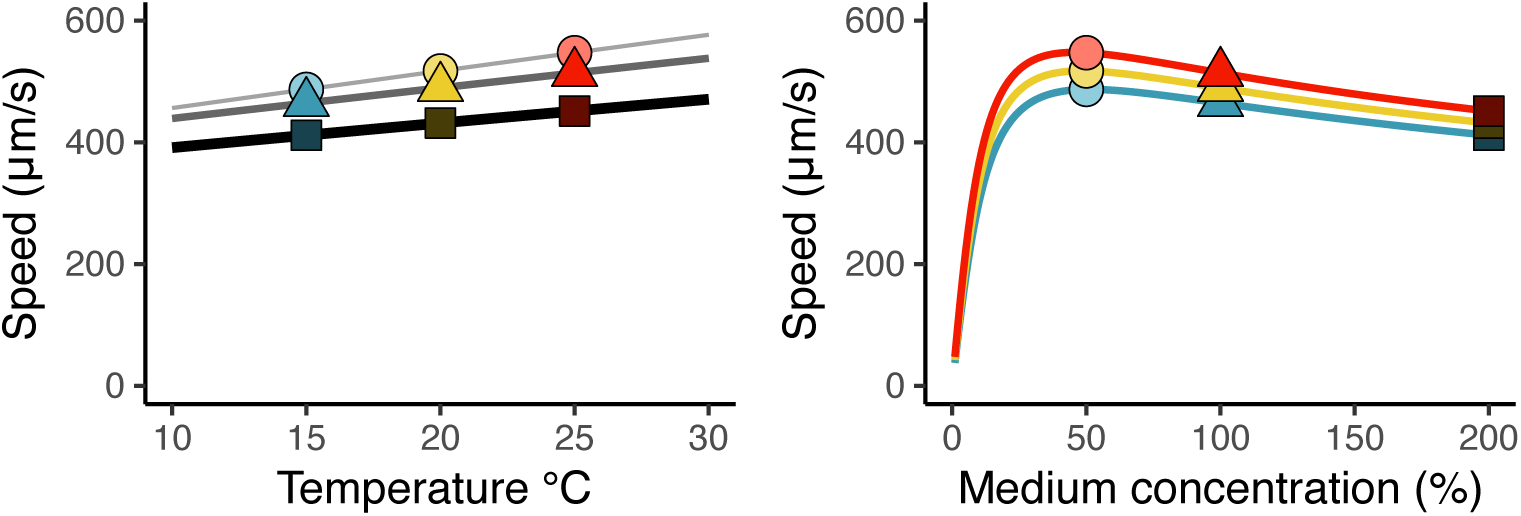
Estimated optimal speed as a function of medium concentration and temperature; model details are provided in the text. Colours: blue = 15 °C; yellow = 20 °C; red = 25 °C; light, narrow line and circles = 50 % growth medium concentration; grey line and triangles = 100 % growth medium; dark, thick line and squares= 200 % growth medium.

This modelling approach cannot be considered predictive in a quantitative sense because of the large number of simplifying assumptions and parameters, such as considering a uniform distribution of non-moving prey, ignoring the additional costs of movement from increased predation and, for the example of fig. 1, focusing only on parameters related to the environment (temperature, viscosity, resources) while fixing those related to the organism itself (cell size and shape, maximum feeding rate, efficiency of swimming, etc.)

Nevertheless, the existence of optimal values of swimming speed that are generally different from those passively determined by the scaling of metabolic rate will be sufficient for us to formulate our third prediction, that the changes in movement speed and metabolic rate that are expected to happen during an acute, short-term exposure to a new environment are unlikely to be sustainable in the long-term. Some phenotypic or ‘behavioural’ changes will be required to compensate for the introduced suboptimal allocation of energy. For example, at high temperature organisms might try to limit the increase of speed below the simple thermodynamically-driven increase that we expect to observe immediately after the exposure to a new temperature condition, because the faster speed does not ‘pay off’ in terms of energetic returns; they could also try to limit the increase of metabolic rate itself, also to cope with the less than proportional increase of energetic returns from feeding. At low temperature, similar compensatory changes could happen, this time in the opposite direction. Similarly, the concentration of resources in the environment could also lead to a modulation of movement speed, particularly when resources are scarce and feeding is limited by prey encounter.

### Consideration: Changes in cell volume and morphology can modulate metabolic rate and movement speed

Phenotypic changes that affect the total metabolic rate of an organism, and the allocation of metabolic energy to maintenance, movement, growth, and reproduction, could involve changes in its physiology, e.g. biochemical regulations related to enzyme structure, changes in the concentration of regulatory intracellular molecules, density of mitochondria etc. (see e.g. Angilletta, 2009 for a review). However, also morphological traits such as cell volume and elongation can affect metabolic rate and movement speed. We see this in equation 2, where metabolic rate scales with cell volume and in equation 3, where power dissipation is dependent on parameters *s* which is itself proportional to cell radius. For this reason, we can expect that modulations of metabolic rate and movement speed can be mediated by changes in cell volume and shape, in addition to changes in physiology.

Guided by these different considerations and theoretical predictions, we now move to empirical measurements on how *Tetrahymena pyriformis* responds to changes of environmental conditions.

### Conceptual summary

As a guide for the reader to navigate through the several pages of methods and results, table 1 provides a conceptual summary of theoretical predictions and of factors that could play a role in determining phenotypic variation. A similar table, presented at the end of the results section, will review these same points with a focus on the experimental evidence provided by our experiments (table 2).

**Table 1:** Summary of theoretical predictions and factors to be controlled.

| Prediction | Timescale | Explanation |
| --- | --- | --- |
| (1) High temperature will lead to higher metabolic rate and faster population growth, if resources are not limiting. | Both short and long timescales. | The rate of metabolic reactions increases with temperature. |
| (2) Movement speed will increase with temperature, but the increase will be described by a lower activation energy than metabolic rate. | Short timescales only. | At low Reynolds numbers, movement speed scales as the square root of the metabolic power invested for movement. |
| (3) Under prolonged exposure to stable temperature and resources, movement speed should revert to less extreme values than under short-term exposure. | Medium and long timescales. | Increases or decreases of movement speed induced by temperature may be suboptimal; optimal speed is determined instead by resource acquisition rate and swimming power costs. |
| Factors to be controlled | Reason |  |
| Does acclimation affect metabolic rate through physiological up- and down-regulation? Or through body size changes? | *Per capita* metabolic rate can be modulated by regulating metabolic pathways, but it is also affected by body size: larger organisms consume proportionally more energy. What is the contribution of these non-exclusive mechanisms? |  |
| Does acclimation affect movement speed through physiological regulations, e.g. by increasing ciliary activity? | Movement speed can also be affected by morphological changes in body size and shape; for example, larger organisms are likely to move faster. |  |
| Effects of temperature on oxygen availability. | Temperature-dependent oxygen availability – rather than feeding resources – could limit maximum cell size and population growth rates. |  |
| Different locomotion modes – swimming and crawling – could affect the relation between energy expenditure and movement speed. | Theoretical arguments supporting predictions (2) and (3) above are based on organisms acting their forces on the water and do not necessarily apply to crawling organisms. |  |
| Instantaneous changes associated with temperature ‘shocks’. | Over very short timescales, the physiology and movement of microorganisms could be altered by the *change of environmental conditions*, rather than by environmental conditions themselves. |  |

**Table 2:** Summary of evidence related to the points formulated in. table 1.

| Prediction / question | Empirical observations |
| --- | --- |
| (1) High temperature will lead to higher metabolic rate and faster population growth, if resources are not limiting. | Only partially observed: temperature affected population growth rates in the experiments (fig. 3(b) and Appendix sec. 2.8). Metabolic rate increased with temperature over short timescales (evidenced by the positive activation energy in fig. 4(a) and (c) and Appendix sec. 2.2), but much less over long timescales, (Appendix fig. 13), possibly due to acclimation (baseline rates in fig. 4(b) are different). |
| (2) Movement speed will increase with temperature, but the increase will be described by a lower activation energy than metabolic rate. | Broadly observed in our experiments, which found consistently lower activation energy for movement speed (fig. 4(d) and (f), and Appendix sec. 2.3) than for respiration (fig. 4(a) and (c) and Appendix sec. 2.2)), broadly confirming the prediction. |
| (3) Under prolonged exposure to stable temperature and resources, movement speed should revert to less extreme values than under short-term exposure. | Observed in our experiments: movement speed measured over a medium-term exposure showed lower values of activation energy in our experiments (fig. 4(g) and (i), and Appendix sec. 2.4), compared to measurements over a short-term exposure (fig. 4(d) and (f), and Appendix sec. 2.3)). |
| Does acclimation affect metabolic rate through physiological up- and down-regulation? Or through body size changes? | Cell volume alone is a good predictor of metabolic rate (fig. 5, left), but physiological regulations are also likely to contribute. |
| Does acclimation affect movement speed through physiological regulations, e.g. by increasing ciliary activity? | Based on the estimated scaling exponent between cell size and movement speed (Appendix, sec. 2.11), cell size accounted largely, but not completely for observed movement speed (fig. 5, right). |
| Oxygen availability could limit maximum cell size and population growth rates. | Oxygen was unlikely to be a directly limiting factors in our experimental conditions (Appendix, sec. 2.9 and 3.2). |
| Different locomotion modes – swimming and crawling – could affect the relation between energy expenditure and movement speed. | This is possible, but our analyses only focused on swimming cells (Appendix, sec. 1.3), which is the main locomotion mode of *Tetrahymena*. |
| Observed patterns are associated with handling or temperature ‘shocks’ rather than with environmental conditions themselves. | In preliminary experiments, movement speed remained stable also over longer recording times (Appendix sec. 1.3). |

## Materials and Methods

### Cell cultures

Axenic cultures of a clonal strain of *Tetrahymena pyriformis*, strain 1630/1W, were obtained from CCAP (http://www.ccap.ac.uk) and cultured at 20 °C in a Proteose peptone - yeast extract medium (20 g Proteose peptone + 2.5 g yeast extract in 1 L deionised water). Throughout, we will refer to this ‘standard’ medium composition as the 100 % concentration, to distinguish it from a 50 % medium (with half the concentration of Proteose peptone and yeast extract) and a 200 % medium, with double concentration of yeast and peptone.

### Design of the experiment

Cells were initially maintained in the exponential growth phase at 20 °C and 100 % medium concentration for approximately 50 days (under constant temperature and in the absence of light) to acclimate to our laboratory conditions in preparation for the experiment. At the beginning of the experiment, cultures were split across 9 different treatments (3 temperature conditions × 3 medium concentrations), and were exposed to these stable growth conditions for at least three weeks (corresponding to a minimum of 20 generations), in four replicates for each condition (see fig. 2 for a schematic diagram of the experiment and Appendix section 1 for a detailed description of the culture protocol). We chose a treatment duration of at least 3 weeks because previous studies had shown that after such duration *Tetrahymena* develop detectable phenotypic changes (Fussmann et al., 2017). Throughout the entire study, exponential growth was maintained by adopting a serial transfer regime, which involved the transfer of approximately 1 mL of culture into approximately 24 mL of fresh growth medium every 1-4 days, as required to keep population densities within the exponential growth range (details in the Appendix, section 1). At the end of the main acclimation period to different temperature and resources, cultures were further split across a wide range of temperature conditions (keeping the same medium concentration at which they were acclimated) to measure thermal response curves for different phenotypic traits (respiration rate, cell volume, movement speed, and population growth rate). Specifically, we measured acute thermal responses for movement speed and respiration. For movement speed, measurements were taken 1.5–3 minutes after transfer to the test temperature. For respiration, measurements were taken between approximately 30 minutes to three hours after transfer. We refer to responses measured over timescales of minutes as acute, corresponding to a ‘short-term, transient exposure’. We also measured medium-term responses, over timescales of a few days (2 to 6 generations after the change of environment, corresponding to 3.5 days on average, with differences depending on incubation temperature) for population growth, movement speed again, and cell volume changes.

**Figure 2:**
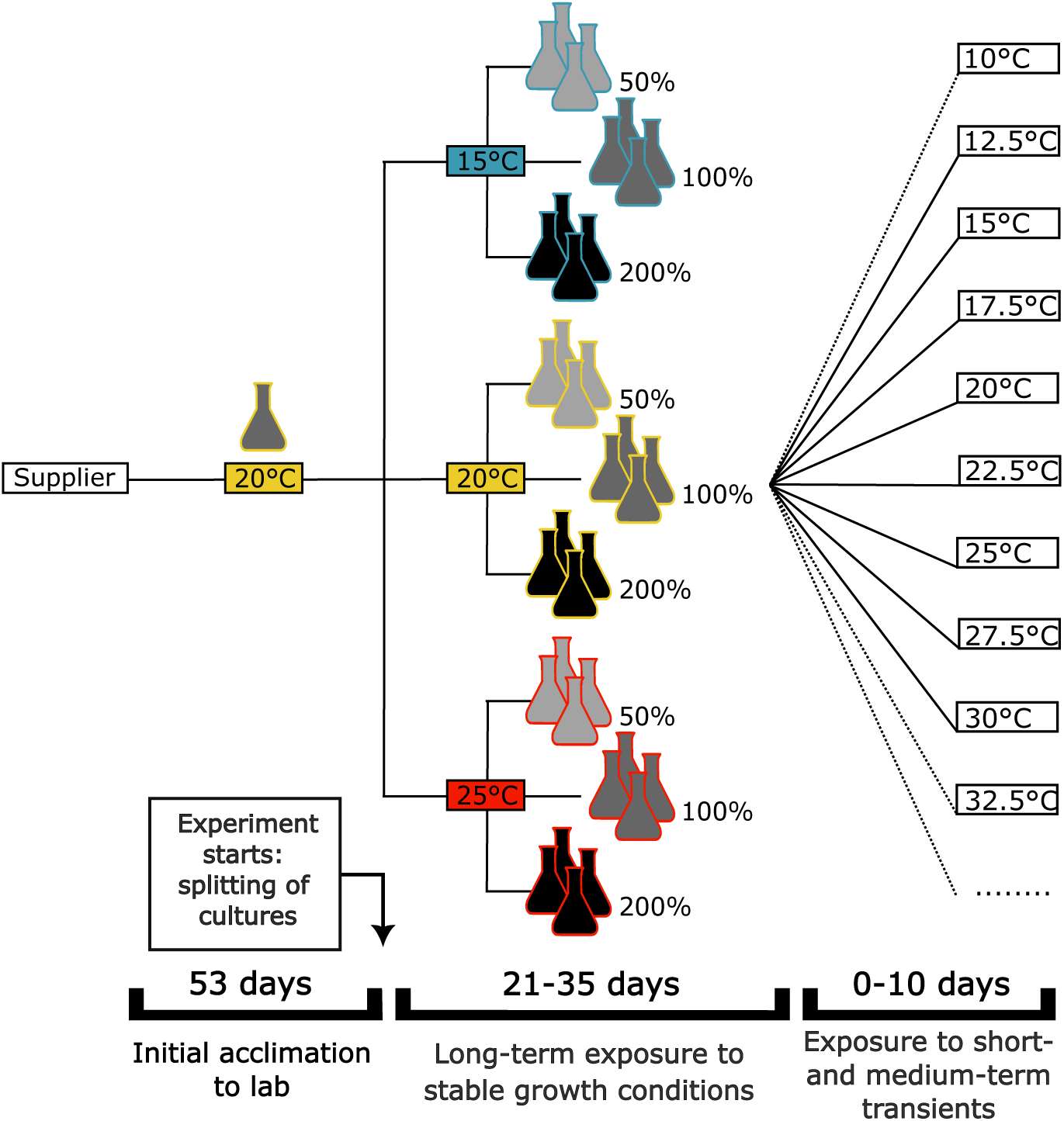
Schematic diagram of the experiment. After an initial acclimation to exponential growth in laboratory conditions at 20°C and 100 % culture medium (over approx. 50 days), cultures were split across combinations of medium concentration (50 %, 100 % and 200 % of the standard culture medium concentration) and temperature, in four replicates and the experiment started. After a minimum of three weeks growing in these stable growth conditions, cultures were further split across a wide range of temperatures to measure their thermal response curves for different phenotypic traits (respiration, movement, cell size, population growth), over different durations of exposure, ranging from a few minutes to a few cellular generations, to differentiate between short-term and medium-term effects.

### Population growth

Population densities were estimated at the time of each subculture into fresh medium (i.e. every 1 to 4 days) by counting cells under the microscope. Assuming exponential growth throughout the experiment, the *per capita* growth rate of the population (or ‘growth rate’, for brevity), expressed in number of generations per day, was calculated as log_2_(*^N^*^(*t*)^/*N*(0))/*d*, where *N*(0) is the density at the beginning of a subculture, *N*(*t*) is the final density of the subculture, and *d* is the length of the subculture period in number of days. Note that we use the logarithm base 2, rather than the natural logarithm, because in the case of organisms that reproduce by binary divisions like *Tetrahymena* this has an intuitive interpretation as the number of generations per day (Reiss and Schmid-Araya, 2010).

### Respiration

Oxygen consumption was measured using a 24-channel PreSens Sensor Dish Reader and 2 mL oxygen sensor vials from PreSens (https://www.presens.de/) inside thermal cabinets at the desired temperature (details of the experimental procedure are provided in the Appendix sec. 1.3). Population-level respiration rates were then converted to rates of energy consumption per individual cell by accounting for population density in the vial, and assuming the equivalence 1 mol O_2_ = 478576 J.

### Imaging

*Tetrahymena* cultures were imaged under the microscope in cell-counting slides (BVS100 FastRead), which provide a constant depth of 100 *µ*m (details of the procedure in the Appendix section S1.3). A microscope camera (Lumenera Infinity 3-3UR https://www.lumenera.com/infinity3-3ur.html) was used to record videos at 30 fps. Temperature while imaging under the microscope was controlled using a temperature-controlled stage (Linkam PE120 Peltier System).

### Video tracking and movement analysis

We used a custom-made software written in *Python* (available at https://github.com/pernafrost/Tetrahymena) to extract trajectories and measurements of cell volume and cell morphology directly from the videos.

The instantaneous movement speed of each tracked particle was measured from the trajectories as the displacement per unit time of the centre of mass of the particle, measured over a timescale of 1/10 of a second (3 frames). We took the median of all the instantaneous speed values along the trajectory as the individual speed of the particle. We excluded from the analysis particles that did not move and cells with an unnaturally small size that might have been incorrectly detected in the image. Unless otherwise specified, in the analyses and in the figures presented here we focus on the top 20 % fastest moving cells within each tested experimental condition, to avoid bias from temporarily immobilised cells. (Further details of the video tracking and movement analysis are provided in the Appendix, section 1.3, including an explanation for the choice of focusing the analysis on the fastest cells).

### Estimation of cell volume

The video-tracking software fits an ellipse around each segmented *Tetrahymena* cell. Cell volume *V* was estimated based on the major and minor axes of the best-fitting ellipse (respectively *l* and *w*) using the formula 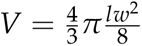.

### Feeding rate

One additional variable that we consider in interpreting our results in terms of the energy budget of *Tetrahymena* is its feeding rate. We do not measure the feeding rate of *Tetrahymena* directly in our experiments, although we have indirect evidence about feeding from other experimental variables such as population growth rate. We model feeding rate based on a Holling type II functional response function in which movement speed determines the ‘attack rate’ (as described in the ‘Theoretical framework’ section, eq. 4).

### Statistical analyses

We used linear mixed-effects (LME) modeling (Bolker et al., 2009; Zuur et al., 2009) to test for the effects of temperature and growth medium concentration (included as fixed categorical factors) on population growth rate and individual cell volume using the *lme4* package in *R* v.3.0.2 (R Core Team, 2022). Culture identity was added as a random effect to account for the multi-level structure of our data (i.e. cultures nested within treatments) and its unbalanced nature (e.g. there is variation in the number of measurements made for different culture lines). Significance of fixed factors (temperature and medium concentration) and their interactions were determined using Satterthwaite’s method for approximating degrees of freedom for the t and F tests from the *lmerTest* package in *R*. Model assumptions were checked using the *performance* package in *R*.

### Fitting thermal response curves

Following the experimental long-term exposure of *Tetrahymena* to stable growth conditions with different temperature and food concentration, cells were tested across a range of different temperatures for measuring thermal response curves. This required first splitting each culture into multiple samples to expose the cells from each line to a wide range of temperature conditions over either a short timescale (minutes) or a medium-length timescale (days) as illustrated in the schematic diagram of fig. 2. Under these different temperature conditions, cells were measured for respiration, movement speed, population growth and cell volume. Thermal response curves were obtained by fitting the Sharpe-Schoolfield model (Sharpe and DeMichele, 1977; Schoolfield et al., 1981) implemented in the *rTPC R* package (Padfield et al., 2021).

The model assumes that the rate *R*(*T*) of the process at a given temperature increases following an Arrhenius-type relation with temperature, but also accounting for the plateauing and subsequent rapid decline of biological rates at very high temperature. In most of the analyses that we report throughout the manuscript, we focus on the Arrhenius-type part of the curve (where rates increase for increasing temperature), which can be summarised as

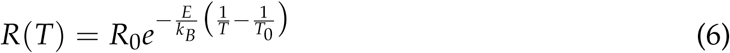

and is described by two parameters: a value of activation energy *E*, which describes how fast the rate increases with increasing temperature, and a reference rate *R*_0_, which is the rate of the process at the reference temperature *T*_0_. These are the two parameters that we report to summarise the experimental data. Throughout all our analyses, we set *T*_0_ = 293.15 K (20 °C). *k_B_*is the Boltzmann constant and *T* is the environmental temperature expressed in Kelvin.

The full Sharpe-Schoolfield equation in its *rTPC R* package version (Padfield et al., 2021) is:

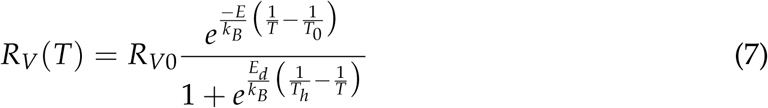

where *R_V_*(*T*) is the rate (already accounting for body size effects) at a given temperature *T*, *E_d_* is the deactivation energy, *T_h_*is the temperature at which 50 % of the units of a hypothetical rate-controlling enzyme become inactive, while *E* and *k_B_* are the activation energy and Boltzmann constant, respectively, as in the previous equation. *R_V_*_0_ is the rate at a reference temperature *T*_0_. Note that equation 7 is similar, but not identical, to the original equation proposed by Schoolfield (eq. 7 of Schoolfield et al., 1981), as the *rTPC* equation does not include a dependence on temperature in the pre-exponential factor, presumably as a way to facilitate the comparison with Arrhenius-type models.

## Experimental results

### Population growth rate and cell size in *Tetrahymena* populations exposed to different environmental conditions

After an initial transient during which cells acclimated to their stable growth conditions, population growth rate remained stable over time. This is illustrated in fig. 3a, where the fitted curves with zero slope shown in the figure were preferred (i.e. they had lower Bayesian Information Criterion) to an alternative linear model with non-zero slope over the temporal range for which the fit is shown in the figure (i.e. sometimes excluding the first measurement, or the first few measurements, after the change of environment). Note that while the growth rate remained constant within a given environmental condition, the baselines were different, i.e. some environments can support faster growth than others.

**Figure 3:**
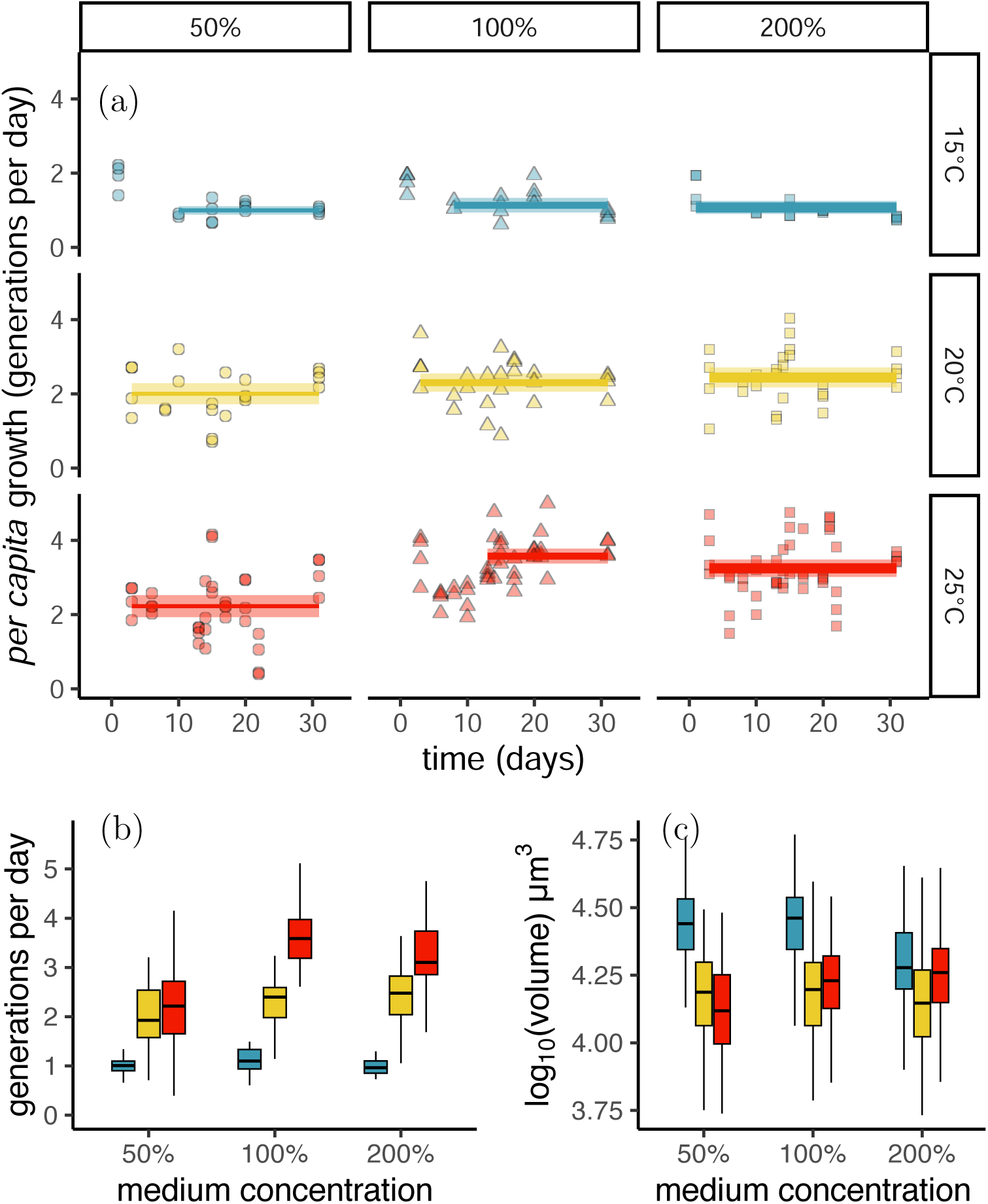
Effect of long-term exposure to different growth conditions on *Tetrahymena* populations. (a) *Per capita* growth rate of experimental cultures throughout the culture period. Each marker corresponds to the measurement of one culture line on a particular day. (b) Box-plot of the growth rates during the subculture period (temporal data are combined together). (c) Volume of individual cells in each treatment, measured after at least three weeks of exposure to stable culture conditions. In all the boxplots, horizontal lines are medians, boxes extend from the 25^th^ to the 75^th^ percentiles, whiskers extend to 1.5 times the interquartile range. Appendix section 2.1 presents additional measurements of cell size in terms of length and width.

Consistent with the expectations (prediction 1), temperature was a strong predictor of *per capita* growth rate (LME: F_[2,227]_ = 142.71, *P <* 0.0001; fig. 3b), with growth rate generally increasing with temperature. There was also evidence for an effect of growth medium concentration on *per capita* growth rates (LME: F_[2,227]_ = 15.88, *P <* 0.0001; fig. 3b), but the relative effect was small compared to temperature (fig. 3b). Finally, there was also evidence of an interaction between temperature and concentration (LME: F_[4,227]_ = 7.82, *P <* 0.0001; fig. 3b): at low concentration of nutrients, *per capita* growth rate increased by 124 % from 15 °C to 25 °C; while at intermediate and high concentrations of the growth medium, the *per capita* growth rate increased by over 200 % over the same temperature range. This pattern is consistent with the interpretation that the *per capita* growth rate is limited by the availability of food when food availability is scarce, but it becomes asymptotically unaffected by the concentration of the growth medium when nutrient resources are in excess of a limiting value. In fact, an additional experiment in which we systematically varied the abundance of nutrient resources in the growth medium over a wider range of concentrations confirms this finding, showing that over a wide range of temperature conditions, the growth medium has a direct limiting effect on *Tetrahymena* growth only when its concentration is low (typically well below 100 %). We refer the interested reader to the Appendix, section 2.8, where the results from this additional experiment are presented.

There was also very strong evidence for an effect of acclimation temperature (LME: F_[2,679]_ = 440.5, *P <* 0.0001; fig. 3c) and growth medium concentration (LME: F_[2,1196]_ = 29.2, *P <* 0.0001; fig. 3c) on cell volume. Specifically, cells transferred to a low temperature increased their volume, but this effect was reduced at high concentration of resources in the growth medium, with evidence of an interaction between temperature and concentration (LME: F_[4,2089]_ = 54.6, *P <* 0.0001; fig. 3c). To put this effect into context, cells approximately double in volume throughout the cell cycle before they reproduce by fission. This corresponds to a span of log_10_(2) ≈ 0.3 units in the logarithmic y-axis scale of fig. 3(c), a difference of the same order as the difference between some of the boxes in the figure: recently divided cells at 15 °C were as large as cells cultured at 20 or 25 °C and which were close to their maximum size, near the end of their cell cycle. This indicates that differences in cell-cycle stage alone cannot explain the observed shifts in cell size, since newly divided cells at 15 °C were already larger than cells grown at 25 °C at any stage of their cell cycle.

### Temperature responses for respiration and movement

Focusing first on the effects of the transient conditions at which cells are tested, metabolic rate (respiration) was expected to increase with the temperature at which cells are tested (prediction 1), and we predicted that movement speed would also increase with increasing temperature, but with a less strong effect of temperature than for respiration, i.e. with a lower activation energy (prediction 2). In our data, the metabolic rate of *Tetrahymena* – measured from oxygen consumption – showed a clear dependence on the temperature at which cells were tested, with values of activation energy *E* measured from respiration broadly in the range of commonly reported values for metabolic energy consumption of many organisms (Gillooly et al., 2001; fig. 4(a), and Appendix table 3).

**Figure 4:**
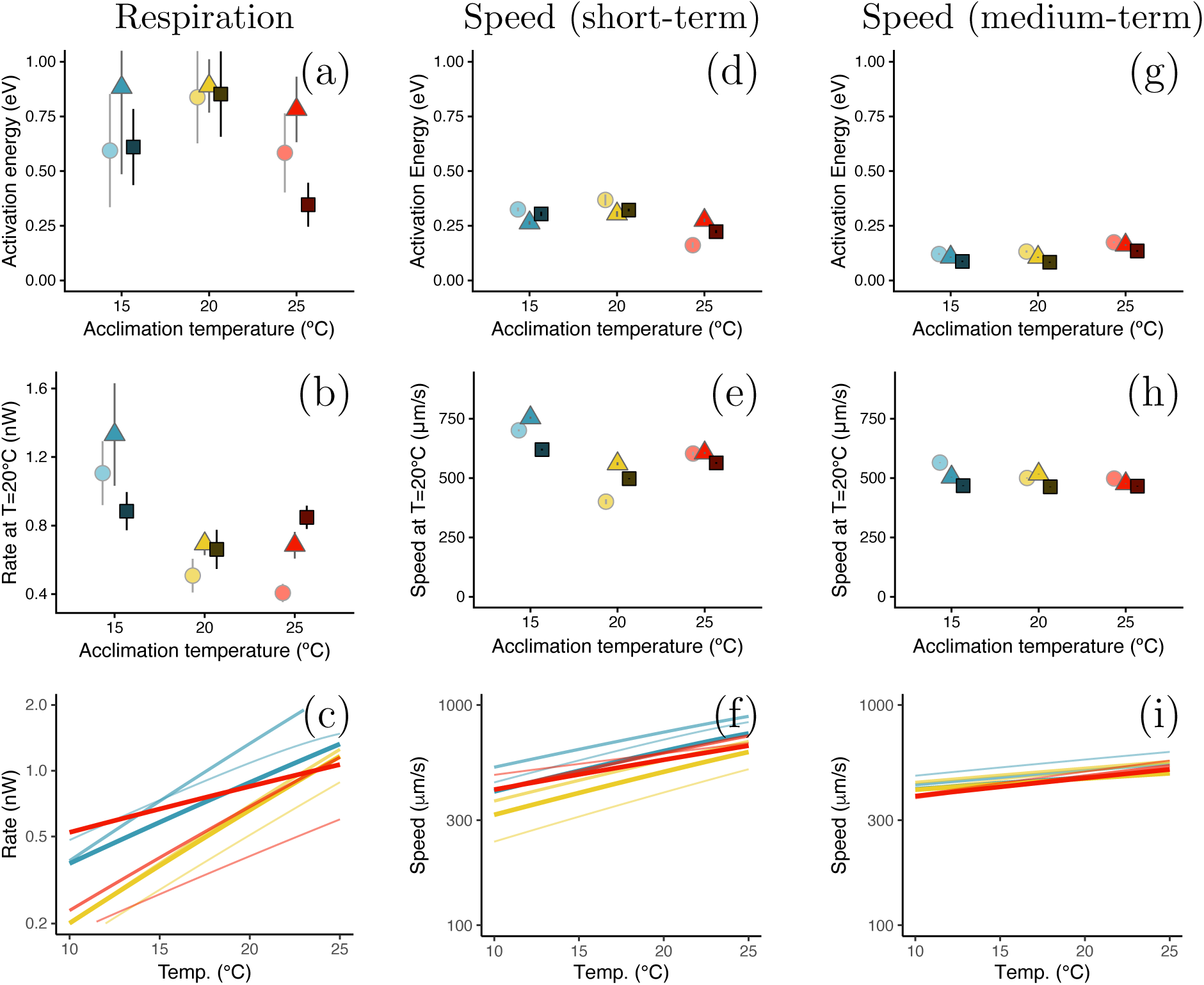
Thermal sensitivity of respiration (a-c) and movement speed (d-i). Top row: activation energy (the slope of the thermal response on an Arrhenius scale, quantifying the temperature sensitivity of each trait), middle row: rate at the reference temperature of 20 °C (representing baseline differences across different long-term acclimation conditions); bottom row: fitted thermal response curves for each of the 9 acclimation conditions (showing the overall relation between the measured parameter and temperature; data and full fits are reported in the Appendix sec. 2). Respiration rate is measured in nanowatts of energy consumption per individual cell. Movement speed is subdivided into an ‘acute’ response where speed was recorded over ‘short-term’ transient exposure to a new temperature, a few minutes after cells were moved to the new test temperature (d-f), and a medium-term response, in which cells were recorded at the first subculture (i.e. a few days) after they were moved to the new test temperature (g-i). Comparing these panels allows separating immediate physico-chemical effects of temperature from responses that emerge over time. Different colour hues correspond to acclimation temperature (blue = 15 °C, yellow = 20 °C, red = 25 °C); bright colours (circles / thin lines) indicate low food concentration (50 % medium), dark colours (squares / thick lines) indicate high food concentration (200 %). Error bars represent 1 standard deviation, estimated via bootstrap with residual resampling in the *rTPC* package.

**Table 3:** Measured parameters of thermal response curves for respiration (short-term response)). Values in brackets indicate the 95 % confidence intervals obtained from a bootstrap procedure with resampling of the residuals.

| Acclimation temperature | Medium concentration | Activation energy (eV) | Oxygen consumption at reference temperature (nW/cell) |
| --- | --- | --- | --- |
| 15 °C | 50 % | 0.594 [0.189 - 1.238] | 1.106 [0.824 - 1.578] |
|  | 100 % | 0.883 [0.409 - 2.075] | 1.331 [0.764 - 1.942] |
|  | 200 % | 0.610 [0.321 - 0.993] | 0.884 [0.657 - 1.106] |
| 20 °C | 50 % | 0.838 [0.543 - 1.448] | 0.508 [0.335 - 0.696] |
|  | 100 % | 0.890 [0.680 - 1.132] | 0.694 [0.576 - 0.820] |
|  | 200 % | 0.852 [0.575 - 1.337] | 0.661 [0.444 - 0.893] |
| 25 °C | 50 % | 0.584 [0.262 - 0.990] | 0.407 [0.296 - 0.502] |
|  | 100 % | 0.782 [0.483 - 1.109] | 0.685 [0.545 - 0.862] |
|  | 200 % | 0.346 [0.195 - 0.594] | 0.848 [0.718 - 0.981] |

Consistent with our second prediction, the values of activation energy for movement speed, measured over a short-term, acute exposure to different temperatures (fig. 4(b)) were lower than those measured for respiration. The values of activation of energy for speed, in the range 0.16-0.37 eV, were on average 42 % of the values of activation energy for respiration (which were in the range 0.35-0.89 eV; see fig. 4(a) and (b), and Appendix tables 3 and 4).

**Table 4:** Measured parameters of the thermal response curves for movement speed (short-term exposure). Values in brackets indicate the 95 % confidence intervals obtained from a bootstrap procedure with resampling of the residuals.

| Acclimation temperature | Medium concentration | Activation energy (eV) | Speed at reference temperature ($\mu\text{m/s}$) |
| --- | --- | --- | --- |
| 15 °C | 50 % | 0.325 [0.309 - 0.342] | 701 [693 - 709] |
|  | 100 % | 0.263 [0.248 - 0.278] | 754 [745 - 762] |
|  | 200 % | 0.304 [0.286 - 0.324] | 620 [614 - 627] |
| 20 °C | 50 % | 0.368 [0.320 - 0.421] | 401 [382 - 423] |
|  | 100 % | 0.304 [0.280 - 0.331] | 560 [548 - 572] |
|  | 200 % | 0.322 [0.310 - 0.334] | 498 [493 - 501] |
| 25 °C | 50 % | 0.161 [0.136 - 0.187] | 603 [589 - 618] |
|  | 100 % | 0.275 [0.260 - 0.291] | 606 [597 - 612] |
|  | 200 % | 0.223 [0.210 - 0.236] | 564 [557 - 570] |

The most notable effect of the acclimation process to different temperature and nutrient concentration conditions was that cold-reared cells (at 15 °C) proportionally consumed more oxygen (fig. 4(b)) when compared at the same reference temperature. Reversing the same argument, through acclimation, cells had partially compensated for the effects of temperature on metabolic rate, having similar metabolic rates when measured each at their respective acclimation temperature (illustrated in the Appendix, section 2.2 and fig. 13).

Cold-reared cells were also comparatively faster than warm-reared cells when compared at the same reference temperature (fig. 4(e)): the average intercept of thermal performance curve for speed of cells cultured at 15 °C was 692 *µ*m/s, while the speed of cells in the other conditions was 538 *µ*m/s, on average. This can also be stated by saying that, after acclimation, cells had partially compensated for the effects of temperature on movement, and they moved at more similar speed when measured at their respective rearing temperature.

These differences of metabolic rate and movement speed across cold-reared and warm-reared cells point to modulations of these variables over a long-term exposure to environments with different temperature and resources that could be consistent with our third prediction – that long-term exposure to a particular environment induces additional phenotypic adaptations and a modulation of these two physiological / behavioural variables. In order to test if the changes of movement speed induced by temperature over a short-exposure persisted beyond a short-term, acute transient exposure, we also measured movement speed of cells that were transferred to a different temperature condition for a medium-term exposure of one subculture (3.5 days on average; fig. 4 - right column). Under these conditions, we observed only a slight increase of speed with increasing temperature (demonstrated by the low values of activation energy, in the range 0.083-0.174 eV, fig. 4(g)). These values of activation energy, being smaller than the values of activation energy obtained for a short-term exposure (fig. 4(d)) support our third prediction and indicate that a timescale of 3.5 days is sufficient for the effects of temperature to manifest in terms of speed changes.

We also indicated in our third prediction that we expected an effect of the concentration of the growth medium on the movement speed of cells grown in each environment. Such a dependence of movement speed on resources is not apparent from the data presented in the right column of figure 4, for instance, in panel (h), cell populations kept at high growth medium concentration (dark squares) have a similar, or slightly lower speed than cell populations kept at low growth medium concentrations (see also the Appendix, table 5 where the actual values are reported). An additional experiment, performed on the same *Tetrahymena* strain, but over a wider range of growth medium concentrations is described in the Appendix, sec. 2.7. This experiment suggests that the weak effect of growth medium concentration on movement speed observed in the main experiment may be due to the specific values of growth medium concentrations tested in the main experiment. In fact, movement speed was affected by resources over a wider range of growth medium concentrations, extending below 50 % – the minimum value used in our main experiment (see Appendix, sec. 2.7 and fig. 19).

**Table 5:** Fitted parameters of the thermal response curves for movement speed measured after a few generations of acclimation to the new temperature. Values in brackets indicate the 95 % confidence intervals obtained from a bootstrap procedure with resampling of the residuals.

| Acclimation temperature | Medium concentration | Activation energy (eV) | Speed at reference temperature ($\mu\text{m/s}$) |
| --- | --- | --- | --- |
| 15 °C | 50 % | 0.121 [0.113 - 0.128] | 565 [562 - 568] |
|  | 100 % | 0.106 [0.103 - 0.111] | 504 [503 - 506] |
|  | 200 % | 0.087 [0.082 - 0.093] | 468 [466 - 470] |
| 20 °C | 50 % | 0.132 [0.124 - 0.142] | 500 [497 - 504] |
|  | 100 % | 0.106 [0.102 - 0.111] | 516 [515 - 518] |
|  | 200 % | 0.083 [0.079 - 0.088] | 463 [461 - 465] |
| 25 °C | 50 % | 0.174 [0.168 - 0.183] | 498 [495 - 501] |
|  | 100 % | 0.162 [0.157 - 0.167] | 476 [475 - 479] |
|  | 200 % | 0.135 [0.128 - 0.143] | 466 [464 - 469] |

### Effect of cell volume on the other variables

Cold-reared cells had a higher respiration rate and higher speed of movement (fig. 4(b) and (e), respectively). As shown above in figure 3(c), these cold-reared cells were also larger. Larger organisms typically move faster than smaller ones. They also have a higher *per capita* metabolic rate, simply because of their size. To what extent do differences in cell size alone explain the other observed differences in respiration rate and movement speed? To address this question visually, figure 5 plots the metabolic rate and movement speed as a function of cell volume for each long-term exposure condition.

**Figure 5:**
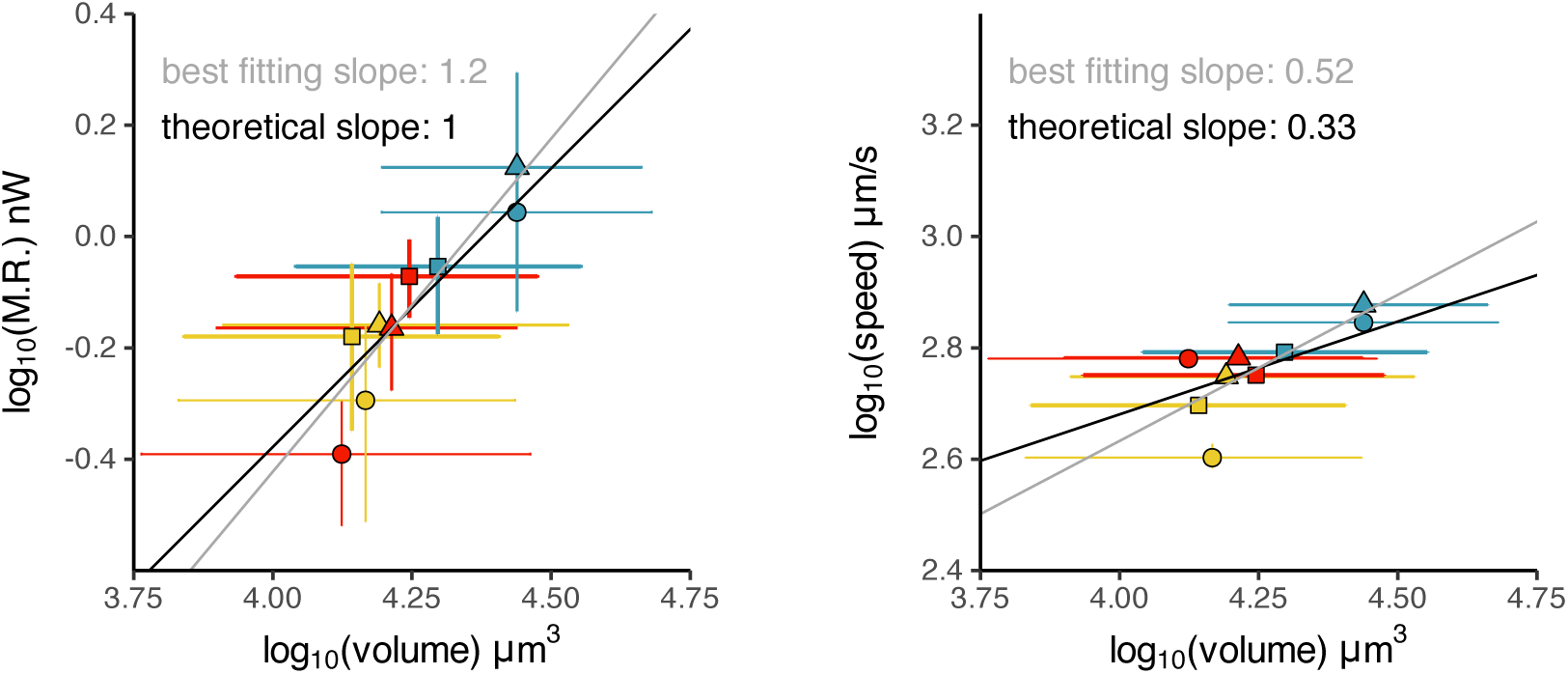
**Left:** Logarithm of metabolic rate measured at the reference temperature (20 °C) vs. logarithm of cell volume in the nine stable environmental conditions to which different *Tetrahymena* cell lines were acclimated. **Right:** Logarithm of movement speed measured at the reference temperature (20 °C) vs. logarithm of cell volume. Black lines: theoretical predictions (see main text). Grey lines: linear least-squares fit through the data. Markers represent the mean value across all replicate lines for each condition, and error bars span the 95 % confidence intervals averaged over all replicate lines.

The black lines superposed to each plot correspond to theoretical expectations under the hypothesis of respiration being directly proportional to cell volume (constant metabolic rate per unit cell biomass, irrespective of treatment) and movement speed being proportional to the linear cell dimension *U* ∝ *V*^1/3^ (a scaling that can be predicted based on the scaling of drag force with cell size, without other effects of the acclimation treatment, and which seems consistent with our empirical data; see the Appendix sec. 2.11 for both the theoretical scaling argument and the analysis of empirical data).

The theoretically predicted relations (black lines in the figure) provide a reasonably good fit through most data points, showing that cell volume is a good single explanatory variable also for respiration rate and movement speed. However, the best-fitting curves (grey lines in the figure) are steeper, indicating that cell volume alone likely does not account for all the variation in metabolic rate and movement speed across treatments. The fact that the fits are obtained by ordinary least squares, without accounting for measurement error – particularly along the x-axis – does not alter this conclusion: fitting techniques that account for regression dilution and for the measurement error in the variable on the x axis would lead to even steeper fitted slopes. The fact that we assumed metabolic rate to scale linearly with cell volume, rather than as volume to the power 3/4, also does not change this conclusion, as a 3/4 scaling would lead to the prediction of a shallower slope. We can hence conclude that cell volume accounts, to a large extent, but not completely, for the changes of metabolic rate and movement speed that we observe in our acclimated populations.

While changes of cell volume can partially compensate for the effects of temperature on metabolic rate and movement speed, cell volume is also involved in trade-offs with other phenotypic traits, and particularly with the *per capita* growth rate of the population and with oxygen limitations. As we made the choice to focus primarily on quantifying changes of metabolic rate and movement speed that made the object of our theoretical predictions, we do not present results related to these additional trade-offs here, but we refer the interested reader to the Appendix, sec. 2.5 and sec. 2.6 for data and analyses that support a negative relation between cell volume and *per capita* growth rate of the population, while we refer the reader to the Appendix, sec. 2.9, and 3.2 for data and theoretical calculations that suggest that while oxygen can become limiting for the metabolism and growth of unicellular aquatic organisms, oxygen was likely not limiting in our experimental conditions, which involved low densities of *Tetrahymena* and exponentially growing populations throughout the experiment.

### Summary of experimental results

In table 2 we re-examine the predictions formulated in the theoretical section (table 1) by considering if the empirical evidence carried by our experiments supports the predictions, or contrasts with them. For each element, We also provide links to the relevant figures or SI sections where the results are shown.

## Discussion

Based on well-established bio-physical principles and energetic considerations, we aimed to predict patterns of phenotypic changes that could be observed in unicellular consumers over short and long timescales. Our main focus was in relation to patterns of movement speed.

### Short-term effects of environmental conditions

We assumed that, over short-term, transient exposure to novel environmental conditions, the direct effects of temperature and resource availability dominate over physiological regulations and morphological phenotypic changes. This assumption allowed us to predict that the movement speed of swimming ciliates should have a lower activation energy than their metabolism, due to the quadratic scaling of power consumption with movement speed for swimmers in a viscous medium (prediction 2). The thermal response curves experimentally measured on *Tetrahymena* broadly confirmed this prediction, as the values of activation energy measured for movement speed under a short-term, transient exposure were lower than those measured for respiration (a proxy for metabolic power consumption): when environmental temperature increased, respiration increased disproportionately more than swimming speed. Previous studies have also found relatively low values of activation energy for short-term movement-speed responses in ciliates. For example, Glaser (1924) reported speed values for paramecia that correspond to an activation energy of approximately 0.37 eV, similar to our results.

Nevertheless, we note that the empirical values of activation energy measured for *Tetrahymena* indicate a greater difference than theoretically predicted, with activation energies for short-term movement speed at 42 % on average of the activation energies for respiration, against a theoretical prediction of approximately 50 %. Such a discrepancy can most likely be ascribed to noise in the data. It could also be due to additional effects that were not considered explicitly in our model, such as an increased energy dissipation at high temperature: for instance, the mitochondria might become slightly less efficient as temperature increases (see e.g. Martinez et al., 2016; Chung and Schulte, 2020; Sokolova, 2023); energy dissipation for swimming could also deviate from perfect quadratic scaling because of complex hydrodynamics within the ciliary layer – ciliary beating represents a substantial component of energy dissipation in ciliates (Katsu-Kimura et al., 2009; Gueron and Levit-Gurevich, 1999). Alternatively, already on the short timescale of movement speed measurements, cells might be rapidly modulating their energy allocation to movement through ‘fast’ regulatory mechanisms such as those based on the modulation of intracellular concentrations of calcium and cyclic nucleotides (see e.g. Brette, 2021).

### Environment-induced mismatches between energy intake and energy expenditure

Besides the short-term changes of movement and respiration, the quadratic scaling between movement speed and power consumption could introduce a mismatch between energy acquisition and energy expenditure, depending on resource availability and temperature. Specifically, with increasing temperature, prey encounter rate and prey capture (which depend on movement) may not keep up with the increase of metabolic energy requirements (prediction 3). This mismatch may require compensatory physiological and structural adaptations to take place when the exposure to different environmental conditions persists. The empirical data provide some support to this idea, for instance in the fact that movement speed becomes almost independent of environmental temperature after medium-term transient exposure to culture conditions with different temperature (as in fig. 4, right panels).

This mismatch between energy acquisition and energy expenditure mediated by both resource availability and temperature has a parallel in autotrophs. In autotrophs, such as unicellular algae, the energy acquisition process – in this case the photosynthesis –has a lower activation energy than the energy expenditure process (respiration), and this mismatch imposes phenotypic response strategies based on the up- or down-regulation of the two processes (see e.g. Padfield et al., 2016). It is interesting to note how – albeit for completely different reasons – similar temperature dependencies describe the energy intake and energy consumption of both primary producers and of ciliate consumers. A consequence of this similarity would be that phenotypic responses that compensate for the mismatch between energy supply and demand – such as changing body size (Deutsch et al., 2022) or changing the number of mitochondria inside the cell (Hochachka and Somero, 2002) – should present similar trends across both autotrophs and heterotrophs, in spite of the fact that the mechanistic origin of the mismatch between energy intake and expenditure could be completely different: the different activation energies of respiration and photosynthesis for autotrophs, and the temperature-dependent scaling of movement speed and prey encounter for heterotrophs.

### Physiological acclimation mediated through body size changes

Some of the major changes associated with growing in different environments for *Tetrahymena* were the change in *per capita* growth rate of the population, and changes of individual cell sizes (fig. 3). The reduction of body size with increasing temperature is a well-studied phenomenon, also described in unicellular protists (Atkinson et al., 2003). Various possible explanations have been put forward for this phenomenon. These range from temperature-induced limitations in oxygen supply relative to oxygen demand (Verberk et al., 2021; Forster et al., 2012) or relative to other metabolic costs (such as the increasing costs of maintaining membrane gradients in smaller cells, Atkinson et al., 2006; Czarnołęski et al., 2008; Hermaniuk et al., 2021), to developmental changes in the allocation of metabolic resources directed to maintenance and growth (Von Bertalanffy, 1957; Zuo et al., 2012), to cell size optimisation in relation to the available resources at a given temperature (DeLong, 2012, summarised in Weber de Melo et al., 2020). All these factors could limit cell growth and maximum cell size also in our experiments, although it may be difficult to establish a precise causal relation, especially if cell growth is regulated to maintain a ‘safety margin’ against the limiting factors. For example, oxygen concentrations were unlikely to play a direct limiting effect on *Tetrahymena* growth in our experiments, due to the low population densities maintained throughout the experiment (see the Appendix, sec. 1.2 for experimental population densities, Appendix sec. 3.2 for theoretical estimates of limiting oxygen concentrations, and Appendix sec. 2.9 for experimental evidence that cell size is independent of oxygen). This does not mean that oxygen limitations did not contribute to shaping the reaction norms to oxygen concentrations exhibited by *Tetrahymena*. Such reaction norms likely evolved in environments in which oxygen could indeed be limiting (Verberk et al., 2021), even just because, in natural conditions, populations do not indefinitely persist in their exponential growth phase, as we artificially made it happen in our experiments.

While we may not be able to establish a precise causal relation between cell size and the proximate or ultimate causes that limit cell growth, our experimental results contribute to describe how cell size interacts with the other phenotypic traits of metabolic rate and movement speed (fig. 5, and Appendix, section 2.11). Specifically, the fact that large cells have a higher metabolic rate and also move faster could be leveraged by *Tetrahymena* in the acclimation process to different environments. For a small microbial consumer with a short generation time like *Tetrahymena*, changes in cell size can take place rapidly relative to the typical timescales of seasonal environmental fluctuations, meaning that they can effectively complement physiological acclimation mechanisms.

However, cell size is also involved in a trade-off with generation time: large cells consume more energy, and so proportionally take more time, to grow to their maximum size. This could explain why *Tetrahymena* populations growing at 15 °C have a lower *per capita* growth than those kept at higher temperatures, in spite of the fact that the *per capita*metabolic rate is similar (cells growing at 15 °C exhibit higher metabolic than those from other growth conditions when compared at the same reference temperature, fig. 4(b). When accounting for the temperature at which they grow, however, their metabolic rates become similar to those observed under other conditions, as illustrated in more detail in the Appendix, sec. 2.2 and fig. 13).

To see why this trade-off exists, it is sufficient to consider the hypothetical situation in which, through rapid phenotypic acclimation mediated by changes in volume, cells were able to completely compensate for the effects of temperature, to the point that cells across the entire temperature range all had exactly the same *per capita* metabolic rate, they individually consumed food at the same rate, and they respired the same amount of oxygen per unit time, also spending the same amount of energy as they move around. Should we expect these cells to all have exactly the same population growth rate, irrespective of temperature? The answer is no, and the reason lies in the remaining term in equation 1: 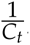, the reciprocal of the energetic content of one cell.

Additional data, shown in the Appendix, fig. 17 and 18 corroborate the idea of a direct relation between *per capita* growth rate of the population and the inverse of cell size: when *Tetrahymena* populations experience different temperature conditions in a medium-term, transient exposure, the reciprocal of cell size and *per capita* growth rates exhibit similar thermal response curves, with also some evidence of a peak followed by a decline at higher temperatures.

Taken together, all these different pieces of evidence support the view of cell volume as one important variable for the regulation of the energy budget of *Tetrahymena*.

In our experiments, exposure to different growth medium concentrations only had comparatively smaller effects on cell size, respiration, and movement speed. Possibly, this was a consequence of the fact that we tested only a limited range of culture medium concentrations (spanning a factor of 4 from 50 % to 200 % medium concentration), combined with otherwise non-limiting growth conditions (i.e. exponential growth phase throughout). Previous studies (Thomas et al., 2017; Bestion et al., 2018) have suggested that the concentration of nutrient resources in the environment becomes a limiting factor for population growth when the metabolic requirements of organisms are high (e.g. at high temperature) and resources are scarce. Our results show some agreement with these predictions, e.g. in figure 3(b), where cells cultured at 25 °C and 50 % medium concentration show a reduced growth rate, and the additional experiments presented in the Appendix, sec. 2.8 also point to an effect of growth medium concentration on *Tetrahymena* population growth that becomes important when food concentrations are particularly low.

## Conclusion

Our study highlights the multiple changes associated with the response of microorganisms to changes in their environment. Short-term, acute responses to changes in the environment are characterised by rapid and marked changes in metabolic activity and movement speed. These short-term changes mainly reflect simple physico-chemical processes, such as the effects of temperature on the rates of metabolic reactions (Brown et al., 2004; Arroyo et al., 2022) and the temperature dependence of water viscosity (Beveridge et al., 2010). Yet, movement and respiration are not affected in the same way by changes of temperature. This likely introduces a mismatch between energy intake and energy expenditure that imposes behavioural and physiological adjustments. In our experiments, the medium- and long-term exposure to a new environment resulted in the mitigation of acute responses, with both metabolic rate and movement speed converging towards similar values, independently of the environment in which cells were cultured. These adjustments were largely, but not entirely, mediated by changes in cell size.

An energy allocation framework can help make sense of empirically observed patterns of phenotypic change. In future, the framework could be further refined and generalised, to derive fully quantitative predictions on how individual organisms and ecological communities could adapt to future, and possibly unprecedented, changes in their environment.

## Acknowledgments

This work was supported by the Royal Society Research Grant RG170282 to Andrea Perna.

## Author contributions

AP: conceptualization, methodology, software, validation, formal analysis, investigation, resources, data curation, writing, visualization, supervision, project administration, funding acquisition; ESR: methodology, investigation, data curation; JR: conceptualization, resources, writing - review and editing; DMP: conceptualization, methodology, formal analysis, resources, data curation, writing - review and editing, supervision, project administration

## Data and Code Availability

The data and analysis code used for this work are all publicly available at https://github.com/pernafrost/Tetrahymena. A snapshot of data and analysis code before manuscript submission is hosted in Dryad https://doi.org/10.5061/dryad.2v6wwpzvv.

## Appendix

### 1 Additional details on methods

#### 1.1 Cell cultures

Axenic cultures of *Tetrahymena pyriformis* strain 1630/1W were obtained from CCAP (http://www.ccap.ac.uk). Before starting the experiment, we acclimated the cultures to our laboratory growth conditions for 53 days at 20 °C (constant temperature, no light) in a Proteose peptone - yeast extract medium (20 g Proteose peptone + 2.5 g yeast extract in 1 L deionised water – this is our ‘100 % medium concentration’). During this time, population density and medium quality were maintained stable by adopting a serial transfer regime, with repeated dilutions into fresh growth medium every 1-4 days (the exact volumes transferred on each subculture and the frequency of the transfers was variable as we aimed at maintaining population densities in each culture flask within the range of exponential cell growth). Throughout the experiment, the cell culture flasks were kept horizontal on a shaker plate to facilitate the exchange of gases between the culture medium and air. At 53 days we started 12 subculture lines, all kept at a temperature of 15 °C, with medium concentrations of either 50 %, 100 %, or 200 % (in four replicates for each combination of temperature and growth medium). At 55 days we also started 24 additional subculture lines, kept at 20 °C and 25 °C, for all medium concentrations (in four replicates each). Together, this makes 9 experimental conditions: 3 temperature conditions × 3 conditions of medium concentration, each in 4 replicates, for a total of 36 parallel cultures. Cultures were kept in these new conditions and maintained in the exponential growth phase by frequently subculturing into fresh medium for a minimum of 21 cellular generations (estimated from population counts), before being measured for movement speed, respiration and cell size over short-term exposure to various temperature conditions (typically, between 10 °C and 30 °C with intervals of 2.5 °C, but also extending to higher temperatures in the case of movement speed). After 33 days of experiment in which cells were kept growing in exponential growth at one of the twelve combinations of temperature and medium concentrations (35 days for the lines kept at 15 °C) (corresponding to at least 38 generations of stable environmental conditions), all cells were taken from their initial culture and split across multiple temperature conditions (between 12.5 and 30 °C), where they were further kept in culture (without further subcultures) for measurements of population growth, changes of cell size, and movement speed over medium-term exposure (i.e. over a timescale of 2-6 generations).

#### 1.2 Cultures were in the exponential growth phase

In our experiment, cell populations were maintained in their exponential growth phase throughout, by subculturing frequently into fresh culture medium (every 1 to 4 days, depending on the estimated cell density of the culture on the subculture days).

In preliminary experiments, we had observed that if we let *Tetrahymena* cultures grow without subculturing for extended periods (e.g. one week), we typically reached maximum densities of the order of 1,000,000 cells per mL. These values are similar to commonly reported values in the scientific literature for *Tetrahymena* species. For example, Hofmann and Cleffmann (1981) report maximum densities between 400,000 cells/mL and 1.7 × 10^6^ cells/mL in *Tetrahymena thermophila*, depending on the culture medium used (in their article, they also provide some evidence that *T. thermophila* population growth was not limited by oxygen availability, even at the highest densities). Reported maximum densities for *T. pyriformis* are also of the order of 1 million cells/mL; for instance, Mogami et al. (2004) identify the late exponential - early stationary phase at 1.2 − 1.6 × 10^6^ cells/mL. DeLong and Hanson DeLong and Hanson (2009) report much lower maximum culture densities for *T. pyriformis*, at 8379 cells/mL, but we believe that there might be a typo or an error in their reporting of measurement units, because not only their maximum density is about two magnitude orders lower than the typical values reported in other studies, but also their reported respiration rates per individual cell are approximately two orders of magnitude higher than what we measure in our experiments.

In our culture conditions, the estimated median cell density was 4440 cells/mL (with a large variation due to exponential population growth), and for this reason we believe that growth took place in the exponential phase. However, in order to test, directly from our data, whether cell growth rate was limited by density-dependent effects, in figure 6 on the left we report the estimated distribution of densities at hourly intervals throughout the entire acclimation period for each of the nine culture conditions (values between direct counts were interpolated with an exponential). On the right panel in the figure, we report the measured growth rate as a function of culture density. If, within the range of culture densities that we used, population growth was not limited by density, then the *per capita* growth rate should be constant independently of density 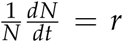, with *r* a constant (*N* is the population density). If, instead, within the same range of population densities the *per capita* growth rate was limited by population density, we should see the *per capita* growth rate decrease at high densities: 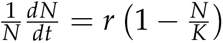, where *K*, also a constant, expresses the ‘carrying capacity’ of the growth environment (growth medium and temperature).

**Figure 6:**
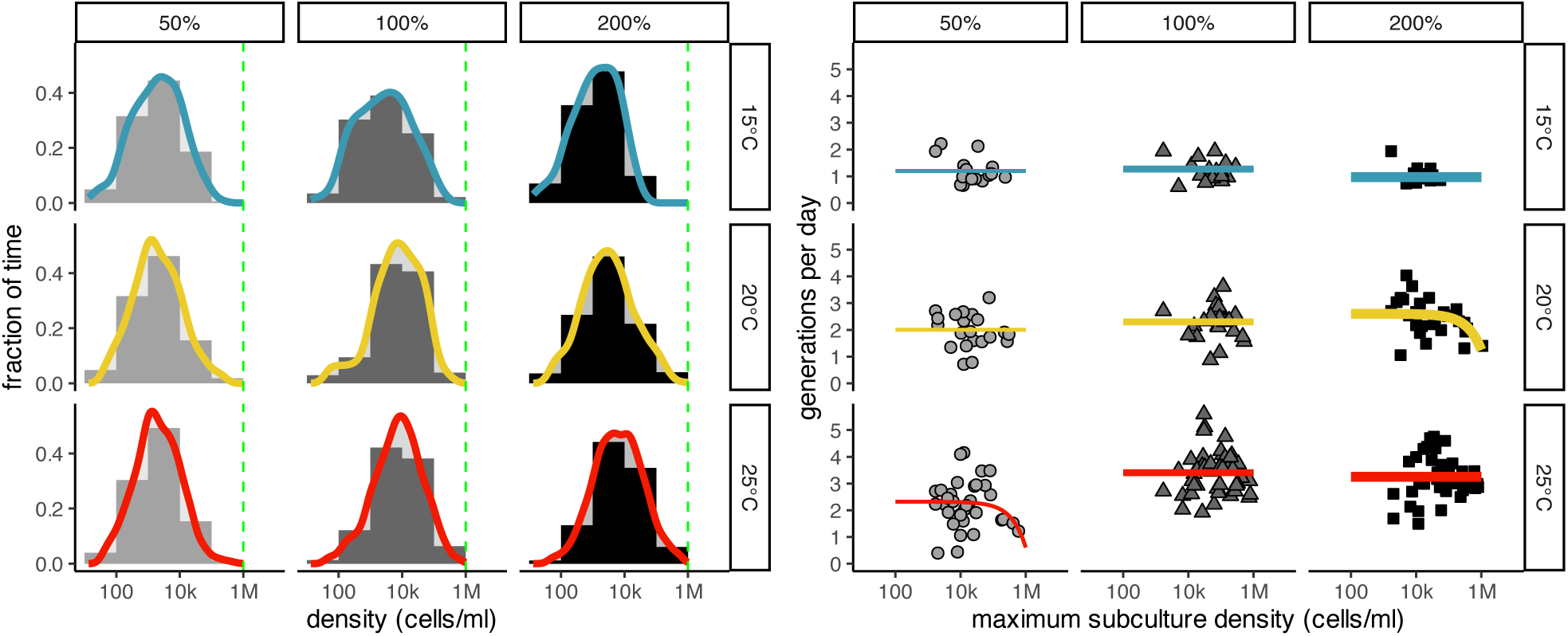
*Tetrahymena* growth happened in the exponential growth phase. **Left:** Estimated time spent by *Tetrahymena* cell cultures at each population density throughout the experiment (long-term exposure to stable growth conditions phase). The dashed green vertical line is added to visually mark an approximate value of maximum density for *Tetrahymena*, at around 1 million cells/mL. **Right:** When the cultures approach the carrying capacity (high population densities), we should expect to observe a decrease in the *per capita* growth rate. The figure plots the measured growth rate of each subculture as a function of the maximum density *N* of cells observed for the same subculture. Fitted lines are the best-fitting model (i.e. the model with the minimum Akaike information criterion AIC) chosen among (1) a model in which the growth rate (expressed in number of generations per day) was constant and independently of culture density throughout the entire range of the data, and (2) a model in which the growth rate was ∝ (1 *N*/*K*) (i.e. decreasing at high density).

Selecting between these two alternative models (based on Akaike information criterion – AIC), the model with constant number of generations per day was preferred in 7 culture conditions out of 9. In the condition with growth temperature T=25 °C, and medium concentration 50 %, the model with density-limited growth was slightly preferred, but the AIC difference was negligible (ΔAIC = 0.132). Finally, in the growing condition with temperature T=20 °C and 200 % medium, the density-limited model was preferred with a non-negligible ΔAIC = 3.702. However, also in this case the fitted value of carrying capacity was large (*K* = 1.9 × 10^6^ cells/mL), and the number of subcultures affected by growth-limiting densities was small (i.e. the density independent model would become the preferred model again if we excluded only two of the 28 subcultures). For this reason we consider the assumption that our *Tetrahymena* cells were kept in their exponential growth phase throughout the experiment to be generally valid.

#### 1.3 Experimental procedures

##### Measurements of respiration

Oxygen consumption was measured by placing experimental populations of known density of *Tetrahymena* in 2 mL oxygen sensor vials, Sensor Dish Reader from PreSens (https://www.presens.de/). In order to control environmental temperature during the measurement, the vials were kept inside thermal cabinets (Lovibond TC 255 S, Tintometer GmbH, Germany) at the desired temperature.

Temperature affects the solubility of oxygen and can also affect the readings of the Sensor Dish Reader directly. Because of this, we had to take a series of precautions to minimize the changes of temperature during the measurements of respiration, in spite of the fact that, necessarily, there is a change of temperature associated with moving cells from their incubation temperature to the temperature at which they needed to be tested for respiration.

In advance of any respiration measurement, we filled small disposable flasks each with approximately 5 mL of sterile culture medium at the desired concentration (50 %, 100 %, 200 %) and stored them at the same temperature at which we would later measure respiration. This was done so that we had fresh culture medium already at the right temperature. We also placed the empty oxygen sensor vials into their holding wells in the Sensor Dish Reader, and we placed the whole kit in the incubator to acclimate at the desired temperature.

Just before the respiration measurement started, we took cells from their incubation temperature, we counted their density in cell-counting slides under the microscope and we transferred the appropriate amount of cell suspension into the previously prepared small disposable flasks with culture medium at the testing temperature, so as to obtain a density of approximately 8000 cells/mL. We then mixed gently by swirling the flasks and we used this diluted suspension to fill the oxygen sensor vial. The vials prepared in this way were then placed back onto the Sensor Dish reader and in the thermal cabinet, leaving their lid open in order to allow temperature and oxygen to equilibrate. After 15 to 30 minutes, we sealed the vials with their lids, and we started the recording.

The density of 8000 cells/mL was chosen as it gave a good compromise between recording time and oxygen depletion across the range of temperature conditions that we tested. At lower densities, cells would only consume oxygen very slowly, leading to long recording times, potentially with some population growth happening during the recording itself, and leaving open the possibility that cells might start showing acclimation responses to the environmental conditions used for recording. Higher cell densities could lead to significant oxygen depletion during the recording, and consequently to short recording times, with the possibility of incomplete temperature stabilization during the recording.

The Sensor Dish Reader has an internal temperature measurement, and we only used the portion of recording for which temperature had stabilized to approximately 0.5°C around the set temperature and there was almost no change in recorded oxygen concentration within the vials used as a control. Control vials were prepared in a similar way as the experimental ones, but they did not contain any *Tetrahymena* cells.

The Sensor Dish Reader recorded the measured oxygen concentration ([O_2_] in units of *µ*mol/L) at 15 s intervals throughout the experiment. Respiration rates were then calculated from the slope of a regression line fitted through the oxygen concentration data during the time interval for which temperature was stable and oxygen consumption was linear. The exact starting time and duration of this interval was variable depending on the recording temperature (oxygen consumption was reduced at low temperature), but as an indication it started approximately 20 minutes after the vials were sealed and ended approximately 1 to 4 hours later (with longer recordings at low temperature), when we stopped the recording, or in a few cases, when oxygen started to get depleted from the culture medium.

Population-level respiration rate was then converted to rate of energy consumption per individual cell by accounting for population density in the vial, and assuming an equivalence 1 mol O_2_ = 478576 J.

After each usage, the oxygen sensor vials were cleaned with deionised water, rinsed with ethanol, and left to dry in the oven at 50 °C for approximately 48 hours.

##### Measurements of speed

*Tetrahymena* movement speed was measured from videos taken under the microscope at the desired temperature. For each measurement, we placed a cell-counting slide (BVS100 FastRead) on the microscope stage to get to temperature; we then gently pipetted a drop of culture (20 *µ*L) directly on the slide. After pipetting, we waited 90 seconds before starting the recording, to let temperature in the culture to fully equilibrate and to exclude transient effects on *Tetrahymena* behaviour associated with the pipetting and with the abrupt change of temperature (rather than with the value of temperature itself). Preliminary experiments in which we monitored *Tetrahymena* movement for up to 20 minutes after pipetting indicated that the movement patterns stabilized a few seconds after pipetting, and remained stable until the end of the 20 minutes period (see below, Appendix section 1.3).

We used a custom-made software (available at https://github.com/pernafrost/Tetrahymena) written in *Python* and based on the *opencv*, *trackpy*, and *scikit-image* libraries to extract trajectories and measurements of cell volume and cell morphology directly from the videos. The software returns measurements for each ‘tracked particle’, where a particle generally corresponds to one individual cell. However, there isn’t a perfect one-to-one correspondence between *Tetrahymena* cells and particles: a cell moving in and out of the field of view of the microscope camera would be labelled as a different particle, and occasionally the identity of two cells swimming close to each other and overlapping in the images could be swapped. Given the extremely large number of individual measurements both across and within experimental conditions, we ignored this small intrinsic potential for pseudo-replication in our analyses.

Speed was measured under two different conditions: a ‘short-term exposure’ and a ‘medium-term exposure’ to temperature. In practice, the **speed data under short-term exposure** were collected by pipetting cells from one replicate line corresponding to one of the nine acclimation conditions (3 temperature conditions crossed with 3 medium concentration values) directly on the microscope slide at a different temperature, ranging from 10 to 40 °C. In order to reduce experimental time with these measurements, only two out of the four replicates were randomly used for these measurements. **Speed data under medium-term exposure** were collected as part of the same experiment that we used to measure thermal response curves for population growth: each experimental replica from all conditions (9 conditions × 4 replicas each) was split into 8 different flasks, which were then transferred to temperatures ranging from 12.5 to 30 °C (12.5, 15, 17.5, 20, 22.5, 25, 27.5, 30). Speed was then measured at the same temperature at which each flask had been stored, after an incubation period corresponding approximately to two to six *Tetrahymena* generations.

##### Exclusion of slow-moving cells

For all the analyses of movement, we focused on the 20 % fastest cells recorded in each measurement under the microscope. This is because *Tetrahymena* typically move by swimming in the water column, and was primarily observed swimming in the water column throughout the entire duration of the experiment when we checked the culture flasks under the microscope. However, for the purpose of video-tracking, the cells were confined in the small volume (∼ 1 *µ*L) and narrow space of the microscope slide (100 *µ*m thickness) and could occasionally be crawling on the surface of the slide, or be temporarily stuck trying to move in the vertical direction, or have their movement otherwise affected by the confinement. The swimming speed of *Tetrahymena* is known to decrease substantially when the microorganism crawls on a surface Ohmura et al. (2018). This effect was also present in our experiments, and led to an approximately bimodal speed distributions in each measured sample (illustrated in figure 7), with fast cells exhibiting nearly straight helical trajectories, suggestive of unimpeded movement, while slow cells produced more meandering paths. Ideally, one could fit the two modes of the distribution, but in practice we used a fixed threshold on the percentiles of speed within each condition, as this offered an easy to implement alternative, which is robust to situations in which the size of the observed population is small (because percentiles can be estimated more efficiently than modes). Moving the threshold to include or to exclude a different fraction of slow-moving cells can shift the estimated value of average speed of a population, but these effects are similar across all experimental conditions and do not lead to major qualitative changes in the patterns that we describe. For example, fig. 8 shows the fitted value of speed at the reference temperature (corresponding to the data in fig. 4(e)) when the threshold for including or excluding cells is moved across the full range, from 100 % (all cells are included) to 5 % (only the top 5 % fastest cells are retained in the analysis). When few fastest cells are retained in the analysis, the measured speed value is obviously higher than if a larger fraction is retained, but the order of the curves remains broadly the same, with cold-reared cells (in cyan) exhibiting consistently faster speed, irrespective of the value of threshold used.

**Figure 7:**
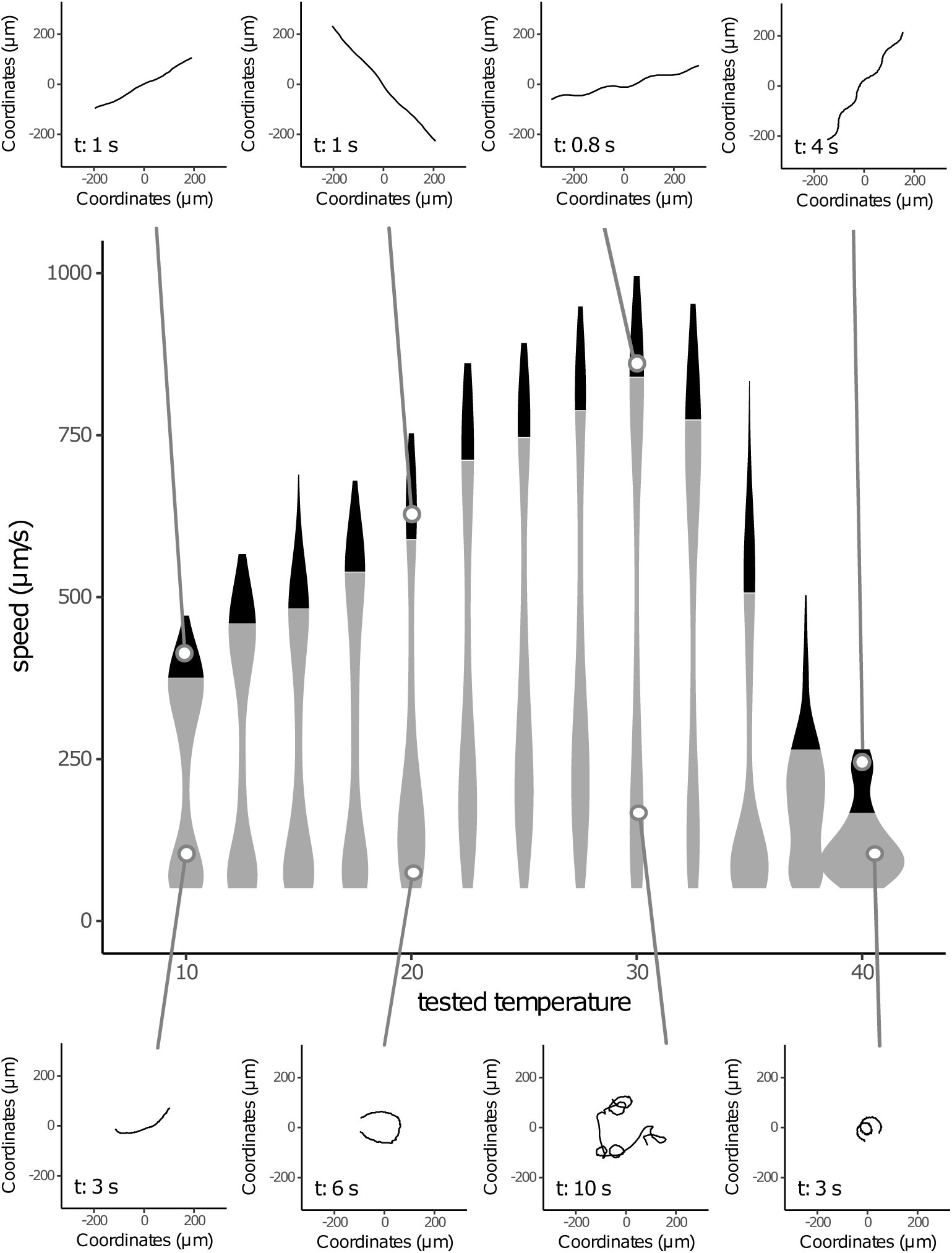
Violin plots illustrating the distribution of *Tetrahymena* movement speed for one single culture line and replicate (cells kept at 50 % growth medium concentration and 15 °C, replicate *a*). The black area in each violin marks the 20 % fastest cells tracked in each sample. The panels at the top and at the bottom of the figure show typical examples of trajectories of individual cells whose median speed is approximately indicated by the lines connecting onto the central figure panel. Note that all the top and bottom panels have the same spatial scale but different temporal length of trajectories shown, with the duration in seconds indicated on the panel itself.

**Figure 8:**
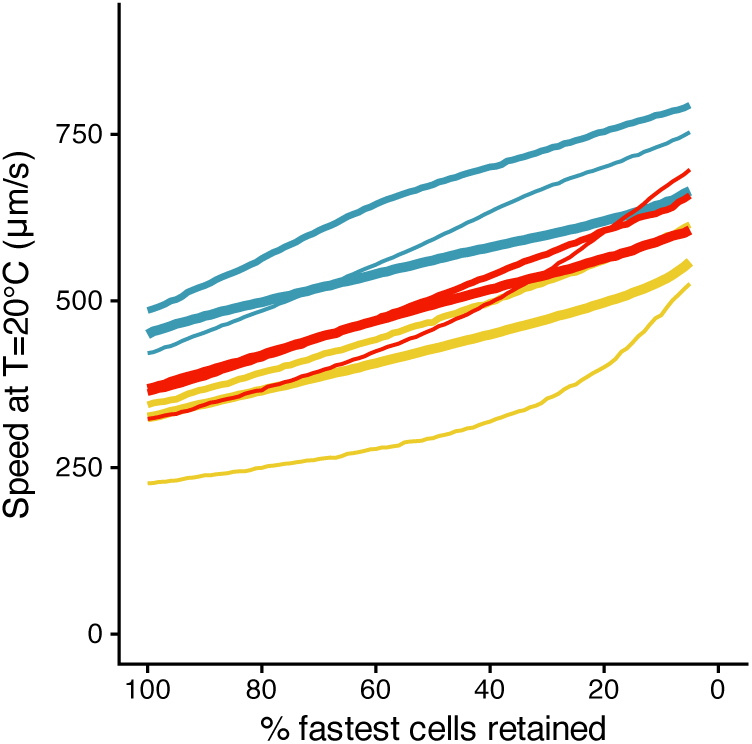
Fitted values of short-term movement speed, measured at the reference temperature, for each of the nine culture conditions, as a function of the fraction of fastest moving cells retained in the analysis. Speed increases monotonically from the left (when the entire cell population is included) to the right (when only a small fraction of fastest moving cells is included), but the vertical arrangement of the curves remains broadly stable, with cold adapted populations consistently exhibiting faster speed than in the other conditions. Line colours indicate different temperature conditions (blue = 15 °C, yellow = 20 °C, red = 25 °C; thin lines correspond to low food concentration: 50 % medium; thick lines correspond to high food concentration: 200 %).

Figure 9 shows the effect of including different fractions of fast moving cells on the values of activation energy measured over a short-term exposure and over a medium-term exposure to different temperature conditions. While the exact measured values varied slightly depending on the fraction of cell populations included, the activation energy measured over an acute, short-term exposure to a particular temperature condition was always higher than the activation energy for medium-term exposure.

**Figure 9:**
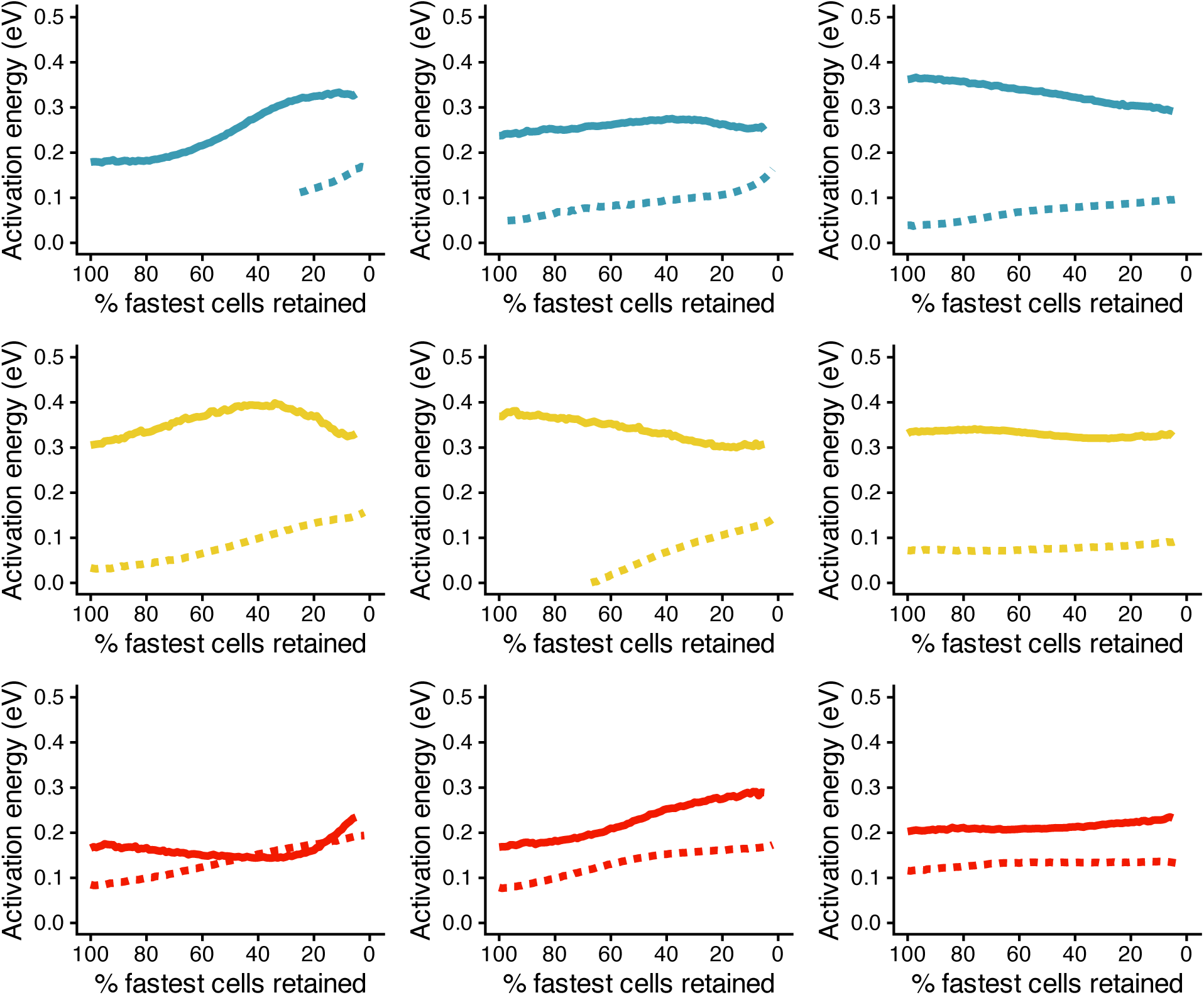
Fitted values of activation energy for movement speed under a short-term exposure (continuous lines) or medium-term exposure (dotted lines), for each of the nine culture conditions, as a function of the fraction of fastest moving cells retained in the analysis. Line colours and the order of plot rows reflect different temperature conditions (top row / blue = 15 °C, middle row / yellow = 20 °C, bottom row / red = 25 °C); plot columns correspond to different growth medium concentrations (left = 50 %, centre = 100 %, right = 200 %). The dotted line does not span the entire range of the data in some plots because the fitting failed to converge for some conditions.

Given that the estimated values of activation energy do not change substantially when different fractions of fastest cells are retained, one might question the opportunity of focusing all the analyses only on fast swimming cells, excluding cells that are crawling or attached to the microscope slide. Previous studies of movement in ciliates often did not distinguish between swimming and crawling cells (e.g. Pennekamp et al. (2019)), or they selected either swimming or crawling cells indirectly, through the type of confinement used to observe cells under the microscope (e.g. Glaser (1924)), or by visually selecting cell trajectories with specific characteristics (e.g. Bullington (1930)). The few published studies that have analysed separately the swimming and crawling motion – often telling the two locomotion modes apart by visual inspection –(e.g. Ricci et al. (1998); Ohmura et al. (2018)) found marked differences in movement speed depending if the studied ciliates were swimming or crawling. For example, Ohmura et al. (2018) report that *Tetrahymena* speed decreased to 23 % of the original value when cells interacted with the glass of the microscope slide. Even when the slide was coated with an anti-adhesive agent, the movement speed of crawling cells remained below 50 % of the speed of cells swimming in the water column.

At the population level, these differences in movement speed between swimming and crawling cells are likely to produce bimodal distributions of speed values, similar to those that we also observed in our experiments (fig.7). The bi-modality of the speed distribution means that common measures of central tendency, such as the mean or the median, do not adequately summarise the data. This problem is compounded by the fact that the proportion of crawling cells in each observed population is affected by the specific characteristics of the experimental setup in which cells were recorded, such as the depth of the observation microscope chamber, the plane of focus of the microscope camera used for imaging the cells, and the surface properties of the microscope slide. For example, Ohmura et al. (2018) report that individual *Tetrahymena pyriformis* cells crawled on a glass microscope slide for 5.2 s on average before returning to swim in the water column, but this crawling duration increased by more than fivefold, up to 26.8 s, when the glass slide was coated with 2-methacryloyloxyethyl phosphorylcholine (MPC; an antiadhesion agent). Given that details such as the depth of the observation chamber vary from one study to another, population-level averages are also difficult to compare across experiments from different research groups.

While the alternance of swimming and crawling may represent a problem when measuring movement speed under the miscroscope, holotrich ciliates like *Tetrahymena* are commonly observed swimming in the water column when they are not confined to narrow observation volumes. In fact, in our experiments, we typically observed *Tetrahymena* cells swimming in the bulk of the culture medium whenever we checked directly the culture flasks under the microscope (a setting that is not suitable for recording because cells swim in and out of the plane of focus). Given that all other population-level measurements that we collected in our study, such as population growth rates and respiration, are based on populations of these swimming, not confined cells, we prefer to report the speed values that better represent the movement of swimming cells.

For these reasons, the movement speed of swimming cells provides a measurement that is better comparable with other present and future studies, and is also better comparable with other measurements of population growth rates and respiration.

##### The measured movement speed is not driven by transient effects

The timescale over which we measured movement speed in our ‘short-term’ speed measurement responses was quite short: only 1.5 - 3 minutes after cells were pipetted onto the microscope slide, and one might wonder if the measured speed is not affected by transient changes of movement, induced for instance by the pipetting itself onto the microscope slide. In a preliminary series of experiments on a different population of the same strain of *Tetrahymena pyriformis* 1630/1W, we recorded cells under the microscope for long periods of 20 minutes, in order to monitor potential changes of speed over time (approximately 30 experimental cultures with different combinations of acclimation temperature and test temperature were recorded in this way).

These preliminary experiments served as a control for the theoretical possibility that, over short timescales, cells could change speed while the temperature of the suspension on the microscope slide was still equilibrating, or because of the stress of being transferred to the new environment; over long timescales, oxygen might get depleted from the very thin layer of suspension – 100 *µ*m – between the microscope slide and its cover, also affecting the movement of *Tetrahymena*. However, our preliminary experiments led us to exclude the presence of transient changes of movement speed associated with the transfer of *Tetrahymena* cells onto the microscope slide. As an illustrative example, figure 10 presents data for cells cultured at 20 °C and 100 % culture medium, and tested across a range of temperature conditions (data for cells cultured at 15 and 25 °C, and tested across the whole range of temperature conditions – not shown here – are also similar).

**Figure 10:**
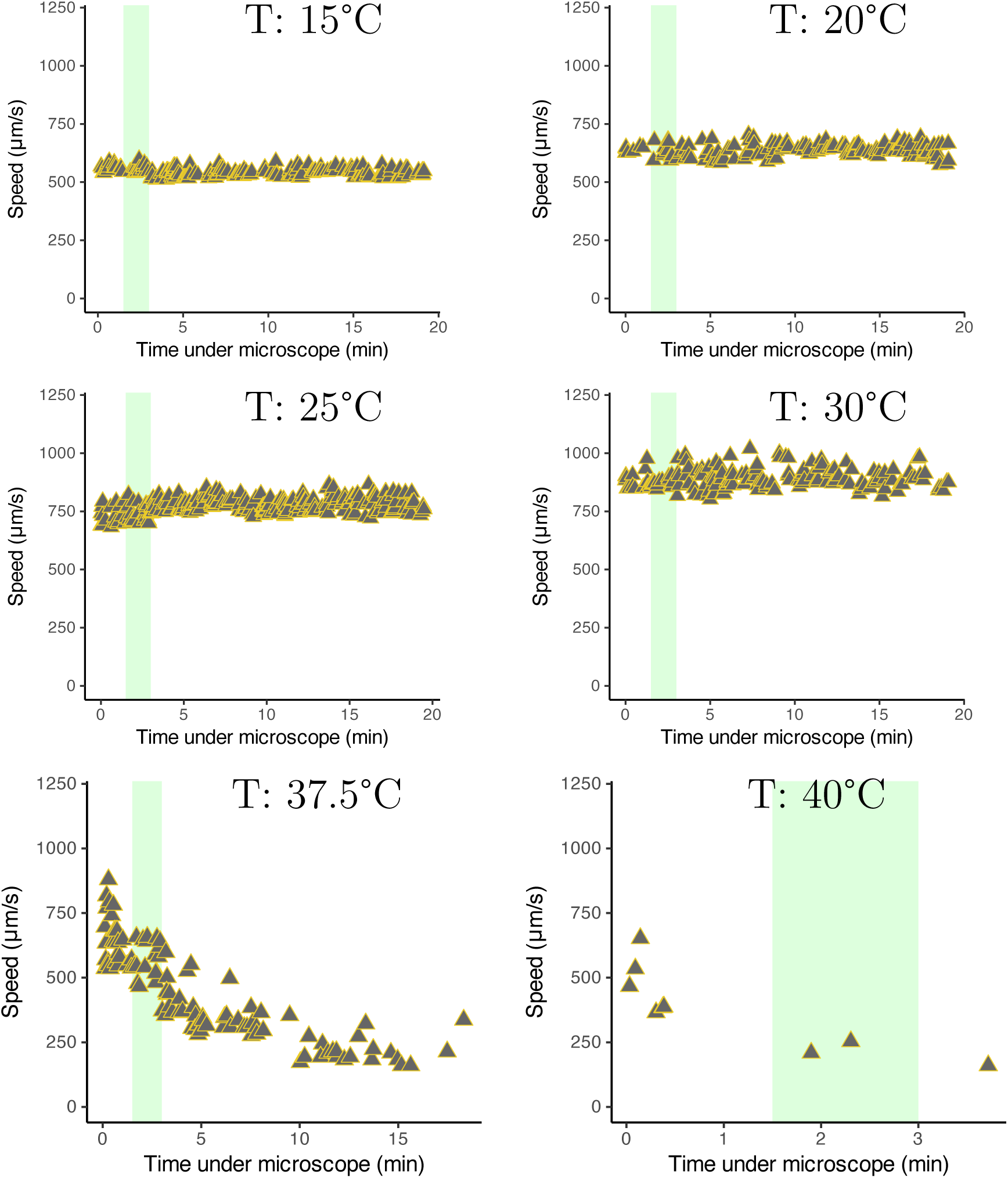
Movement speed over time. Examples of the distribution of *Tetrahymena* speed as a function of the time lapsed since they were transferred onto the slide. In all the panels shown, cells acclimated to 20 °C and 100 % culture medium were transferred onto the slide at a new temperature at time 0 on the x axis, and their speed was measured in the same way as for all the other analyses in our manuscript (the speed values shown in the figure were calculated on the 20 % faster cells in each 1.5 minutes time interval and in each condition). The green shaded region corresponds to the time window that was used in the main experiment (1.5 – 3 minutes).

These data indicate that speed did not consistently increase or decrease throughout the 20 minutes recording period, except at very high testing temperature of 37.5 and 40 °C (bottom plots shown as an example), where cells stopped moving soon after they were transferred onto the microscope slide (in these conditions, fewer cells were detected by the tracking software as time progressed – the software does not detect objects that stop moving for a long time – and the speed of cells that were detected decreased). From these data we could conclude that the recording time-window used in the main experiment (highlighted in green in the figure) was representative of the acute movement speed of our *Tetrahymena pyriformis* cells also over slightly longer timescales, while focusing on the short recording duration, highlighted in green in the figure, allowed us to reduce considerably the amount of time required to run the experiments.

We should note here that Dupont and collaborators (2024) have recently reported of experiments on the related ciliate *Tetrahymena thermophila* in which the movement speed only equilibrated over a longer time-scale of over 30 minutes after the change of temperature. As their results are presented as an illustrative example in their paper, and the experimental methods are not described in full detail, we cannot interpret the differences between our respective studies in terms of different experimental methodologies or differences between the two *Tetrahymena* species used. We could speculate that, perhaps, temperature was still converging to the new value in their experiment (this would be consistent with the pattern that they observe). Irrespective of the reasons behind these differences, we remain confident that – at least in the conditions adopted in our own experiment – speed changes were almost instantaneous, consistent with the interpretation that the changes of short-term movement speed observed in our data are driven by the physics of the system (higher temperatures induce faster chemical reactions and – consequently – a faster movement) rather than by phenotypic changes aimed at buffering the fitness costs of the changed environment, as reportedly would be the case in Dupont et al. (2024). In fact, in our experiments, speed took more extreme values initially, and reverted to less extreme values after a medium-term exposure.

#### 1.4 Missing data points

Our study involves a large amount of data, with 9 treatments, each in 4 replicates, and where each replicate is then tested at 8 or more different temperature conditions for movement, respiration, population growth and cell size. The analysis was automated with custom made scripts in *R* and *Python*, and no data points were selectively altered or excluded. However, for the short-term speed response curves, only two of the four replicates were used (randomly chosen) in order to cut experimenter time and computing resources for the video tracking and analysis by reducing the number of recorded videos from 4 × 9 × 13 = 468 to approximately 2 × 9 × 13 = 234 videos for this specific measurement. Across the various measurements, a small number of combinations of conditions were not recorded because either (1) the density in the culture was too low or too high on the day when the recording was planned and we couldn’t record on another day, or (2) the video file was corrupted, or (3) particularly for cells grown and tested at the same temperature (e.g. grown at 15 °C and tested at 15 °C), we preferred to omit some speed measurement because of logistic constraints with maintaining the culture in axenic conditions (speed measurements took place under the microscope *not* in axenic conditions).

In the measurements of respiration, we removed a single data point for the respiration rate of the culture grown at 25 °C, 200 % growth medium concentration, replicate B, tested at 25 °C. Cells in this culture were recorded as having a respiration rate of approximately 4 nW, much higher than for the other lines and nearby temperature values. We believe that we probably made an error with the dilution in preparation for the oxygen recording (that is, cell density was much higher than the planned 8000 cells/mL), but we cannot confirm this *a posteriori*.

### 2 Additional details on results

This section presents additional experimental results that remain tangential to the main focus of the manuscript, but may still be of interest to some readers. The section also reports thermal response curves in full for each experimental condition (rather than summarising them in terms of their average value of activation energy and rate at the reference temperature.)

#### 2.1 Additional measurements of cell size and shape

Figure 11 reports measurements of *Tetrahymena* cell size and shape. The figure shows that when cells increased their volume at low temperature, the change was mainly driven by an increase in cell length (major axis length, fig. 11(a)), leading to a more elongated shape (fig. 11(c)), while cell width (minor axis length, fig. 11(b)) changed proportionally less.

**Figure 11:**
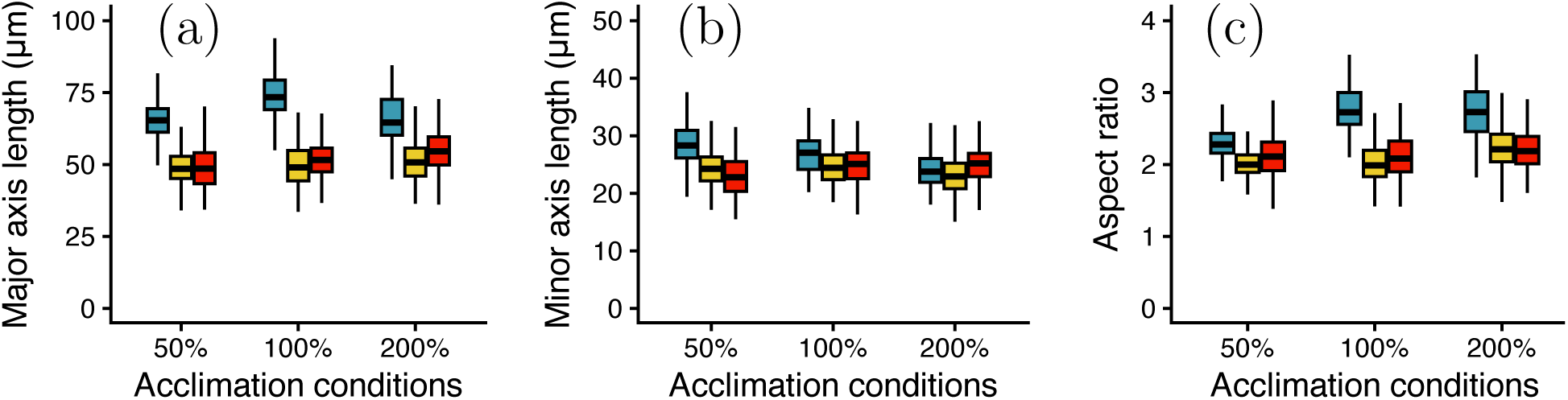
Cell size and shape in different culture conditions. (a) Major axis length (approximately equivalent to cell length). (b) Minor axis length (cell width). (c) Aspect ratio (ratio between length and width). All measurements were obtained by fitting an ellipse along the perimeter of the cell as imaged under the microscope and measuring the major and minor axis of the ellipse. The aspect ratio is calculated as the length of the major axis divided by the length of the minor axis.

We do not have an explanation for these changes of cell shape in terms of their effect on the drag coefficient and the efficiency of swimming (see below, Appendix section 3.6), or in terms of benefits for oxygen uptake (see below, Appendix section 3.2). One possibility is that changes in elongation reflect intracellular or cytoskeletal constraints. Alternatively, a more elongated shape could increase the number of cilia oriented orthogonally to the swimming direction, potentially enhancing propulsion power at the expense of efficiency. A third possibility is that these changes arise as a by-product of mechanisms of cell shape regulation that do not have a direct adaptive significance.

#### 2.2 Thermal response curves for respiration

Figure 12 reports the thermal response curves for respiration under a short-term exposure to the new temperature (approximate measurement range comprised between 20 minutes and 4 hours after the transfer to the new temperature). The fitted parameters of the thermal response curves are also reported in table 3.

**Figure 12:**
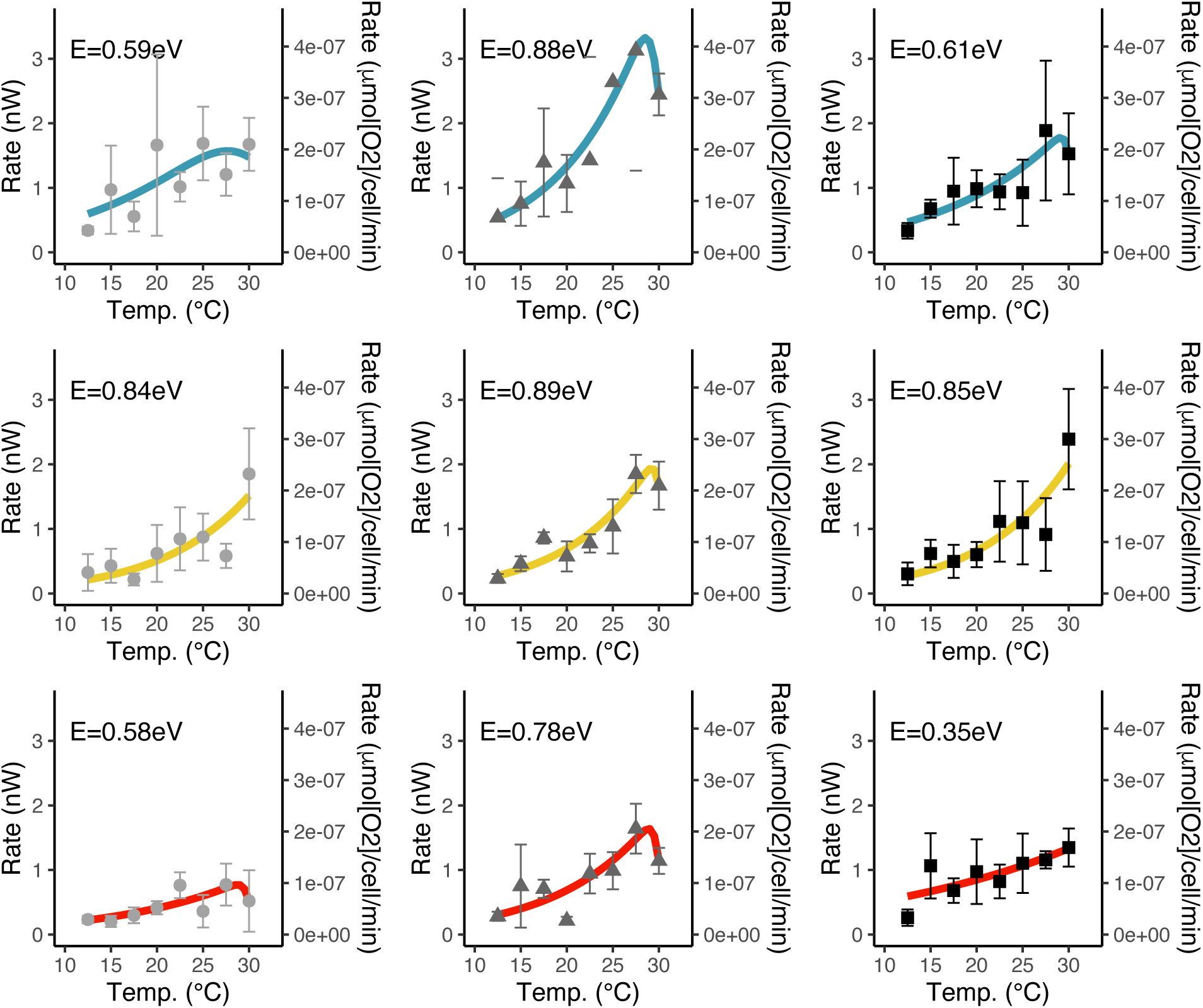
Thermal response curves for respiration. Each row corresponds to a different temperature condition (top = 15 °C, middle = 20 °C, bottom = 25 °C); each column corresponds to a different medium concentration (left = 50 %, centre = 100 %, right = 200 %). Data points represent the mean across four replicates and error bars are ±1 standard deviation. Continuous lines represent fitted Sharpe-Schoolfield curves (*rTPC* package version). The text on each graph reports the value of activation energy *E* for the Arrhenius part of the curve.

While the respiration data in figure 12 are somewhat noisy, when the thermal responses curves are plotted together on the same graph - as in figure 13 - we can appreciate the fact that cells grown under low temperature conditions tend to have higher *per capita* respiration rates, most likely because of their larger volume (see main text, fig. 3). As a result, after acclimation, metabolic rates (measured as oxygen consumption) converge across conditions when each population is measured at its own acclimation temperature.

**Figure 13:**
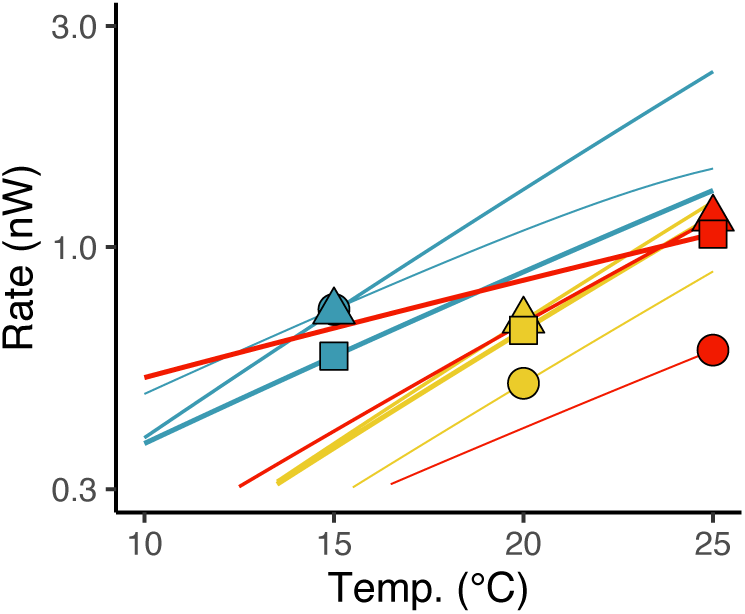
Thermal response curves for respiration rate across all the nine culture conditions. Different colour hues correspond to different temperature conditions (blue = 15 °C, yellow = 20 °C, red = 25 °C; bright colours and narrow lines correspond to low food concentration: 50 % concentration of culture medium; dark colours and thick lines correspond to high nutrients: 200 % culture medium concentration). The markers highlight the values of the thermal response curves at the acclimation temperature: circle = 50 % medium concentration, triangle = 100 % medium concentration, square = 200 % medium concentration.

Based on the average activation energy estimated from individual curves (table 3; mean ≈ 0.71 eV), we would expect per capita respiration to increase by a factor of 2.61 between 15 °C and 25 °C (i.e. *Q*_10_ = 2.61). However, in practice, respiration rates measured at the respective acclimation temperatures only increased by a factor of 1.38 on average (*Q*_10_ = 1.38).

#### 2.3 Thermal response curves for movement speed (short-term exposure)

Figure 14 reports the thermal response curves for movement speed under a short-term (1.5 – 3 minutes) exposure to the new temperature.

**Figure 14:**
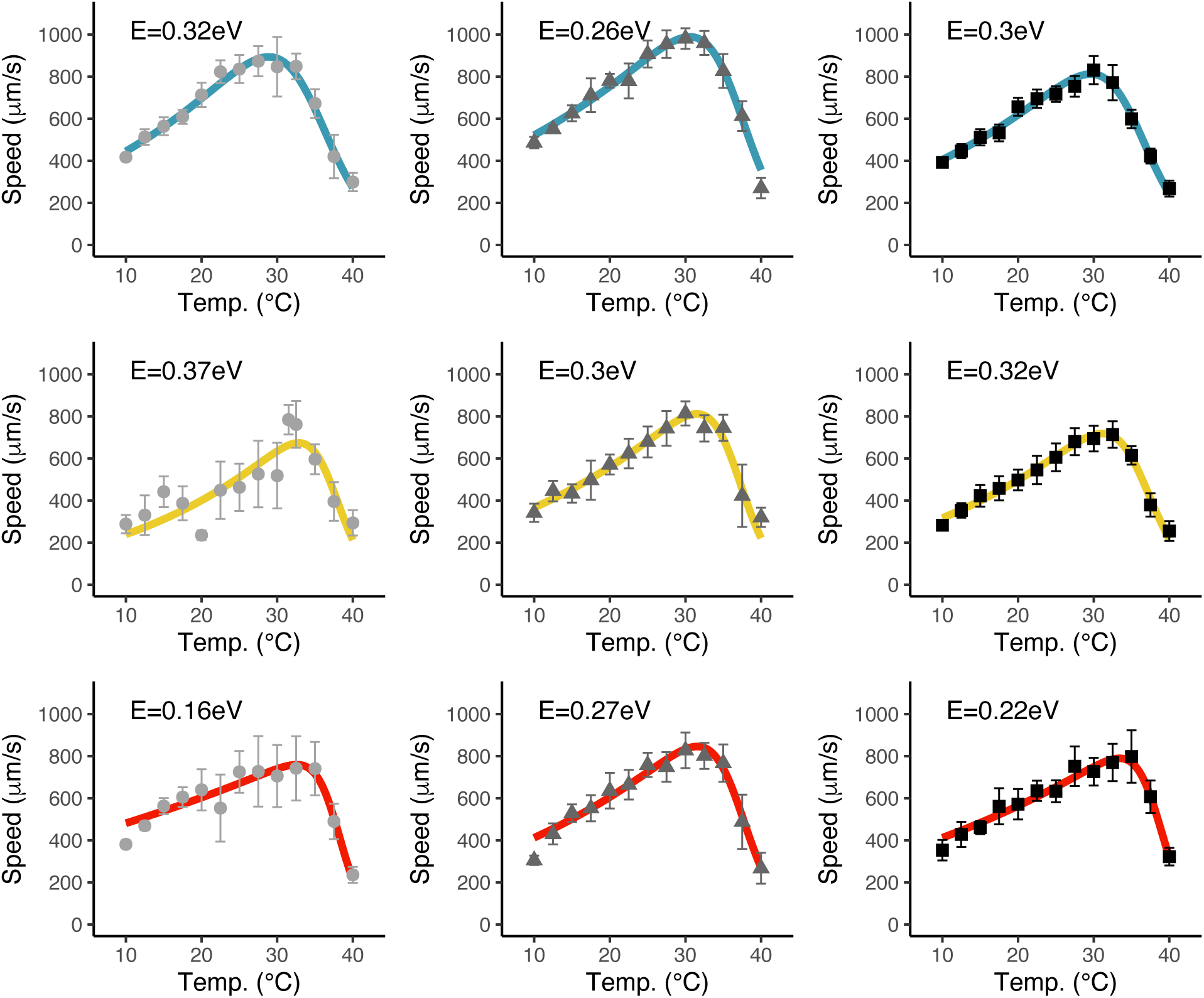
Thermal response curves for movement speed. The arrangement of panels and other details are the same as in figure 12. For each condition of acclimation temperature, medium concentration and tested temperature, we retain the fastest 20 % moving particles, and each marker in the figure represents the mean speed across all retained speed measurements for that condition (typically from two different replicates of the acclimation experiment). Error bars are ±1 standard deviation.

The fitted parameters of the thermal response curves are also reported in table 4.

Given that in these experiments we tested cells through a wide range of temperatures going up to 40 °C, we can also reliably estimate the temperature of maximum movement speed (the ‘optimal temperature’) in each culture condition. This is shown in figure 15.

**Figure 15:**
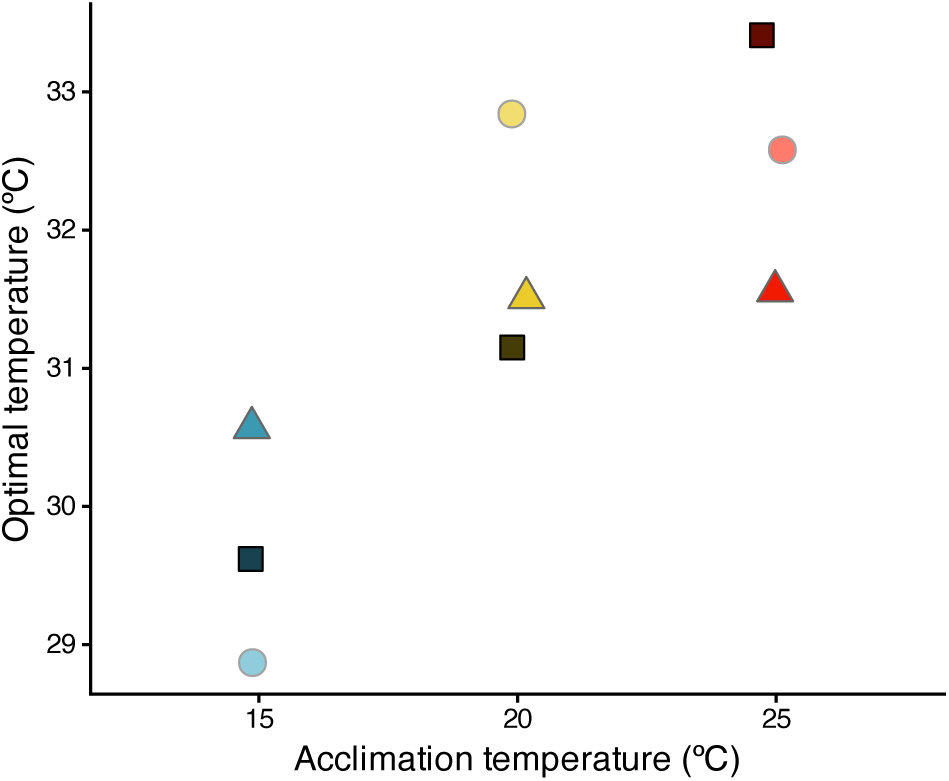
Optimal temperature for speed (temperature at which the movement speed reaches a maximum), as a function of the acclimation temperature. Circles: 50 % medium concentration, triangles: 100 % medium concentration, squares: 200 % medium concentration.

#### 2.4 Thermal response curves for movement speed (medium-term exposure)

Figure 16 reports the thermal response curves for movement speed under a medium-term exposure (days) to the new temperature. In these conditions, an increased temperature only leads to a modest increase of movement speed.

**Figure 16:**
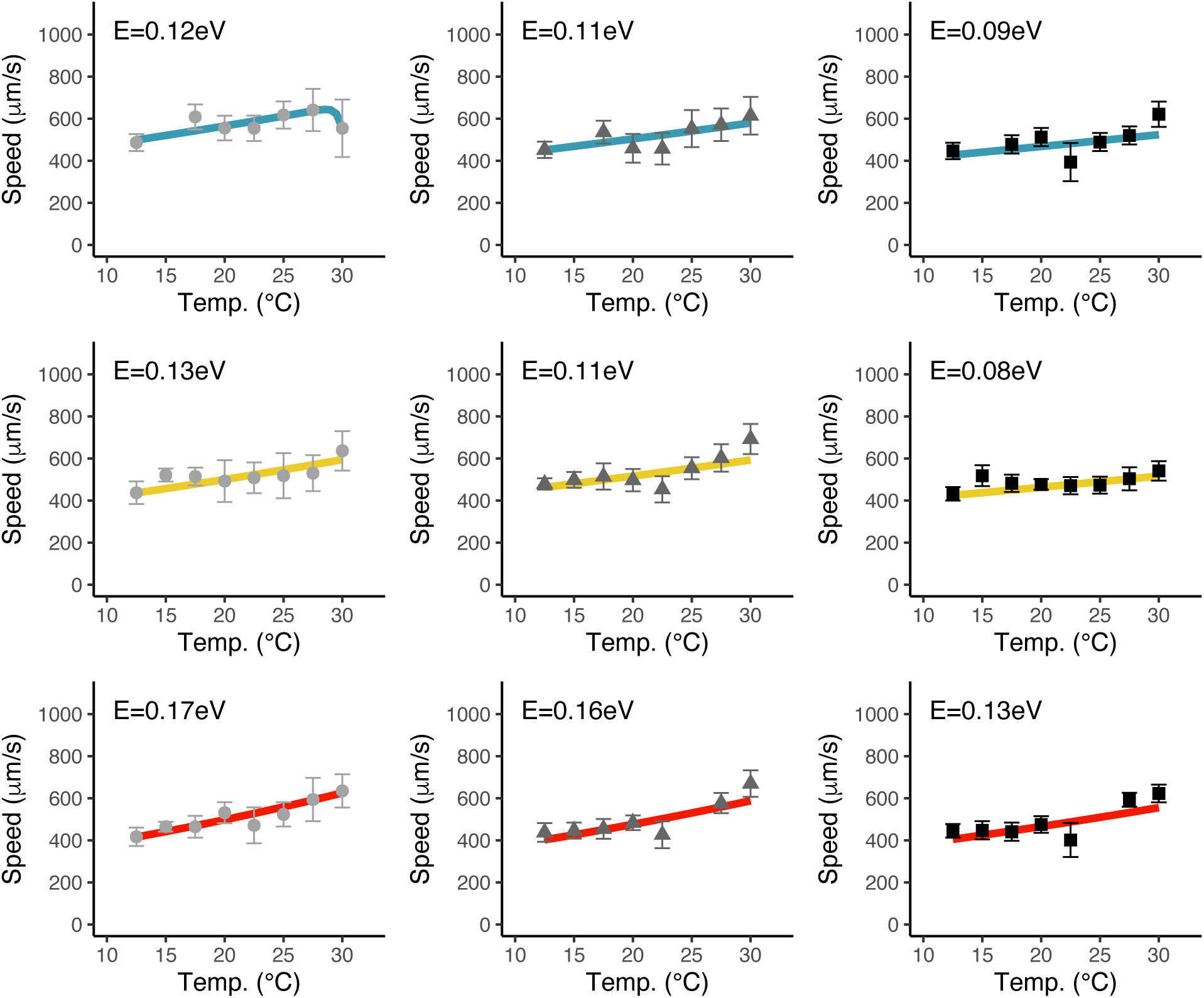
Thermal response curves for movement speed measured after a few generations of acclimation to the new temperature. Each panel corresponds to one acclimation condition (same order as in fig. 12). For each condition of acclimation temperature, medium concentration, and tested temperature, we retained the 20 % faster moving particles. Markers represent the mean across all retained measurements of speed from all four replicates, and error bars are ±1 standard deviation. Continuous lines represent fitted Sharpe-Schoolfield curves (*rTPC* package version, eq. 7).

The fitted parameters of the thermal response curves are also reported in table 5, pointing to low values of activation energy for temperature dependence (0.08 - 0.13 eV) and to a slight decrease of movement speed with increasing medium concentration, at least at 15 and 25 °C. Although confidence intervals are narrow, the effect size is also small (e.g. from 498 to 466 *µ*m/s at 25 °C). For this reason we avoid over-interpreting differences between experimental conditions.

We also note here that a previous study on other ciliate species Beveridge et al. (2010) did not find a difference in activation energy for movement speed when measured in a ‘acute response’ or ‘long-term response’ for *Colpidium striatum*, and found an opposite effect (higher activation energy upon long-term exposure) on *Didinium nasutum*. Interestingly, their studied organisms did not change size with acclimation, which could point to possible mechanistic explanations for these contrasting observations (we will consider the relationship between cell size and movement speed below, in Appendix section 2.11).

#### 2.5 Thermal response curves for population growth

At the end of the acclimation process, each *Tetrahymena* culture was split into different flasks and transferred to temperatures ranging from 12.5 °C to 30 °C for the medium-term transient exposure to different temperature conditions. Cells were then grown at the new temperature and *per capita* growth rates were measured by counting cells at 2 or 3 points in time and fitting an exponential growth function to the counted densities as a function of time.

Figure 17 reports the thermal response curves for population growth measured in this way. The estimated values of activation energy for population growth ranged from 0.37 eV to 0.96 eV, and they did not show a clear pattern depending on the initial acclimation condition, except possibly for a slightly higher activation energy in cells that were originally cultured at high temperature. In practice, there was a relatively large variability in estimates of population growth. A key complication is that thermal response curves for population growth must be measured over multiple generations. Over this timescale, cells begin acclimating to the new temperature conditions. For example, they adjust their movement speed (illustrated before, in fig. 16) and their cell volume (see below, fig. 18). This is a known problem when studying thermal responses, particularly in small organisms with fast metabolic rate (Rohr et al., 2018)).

**Figure 17:**
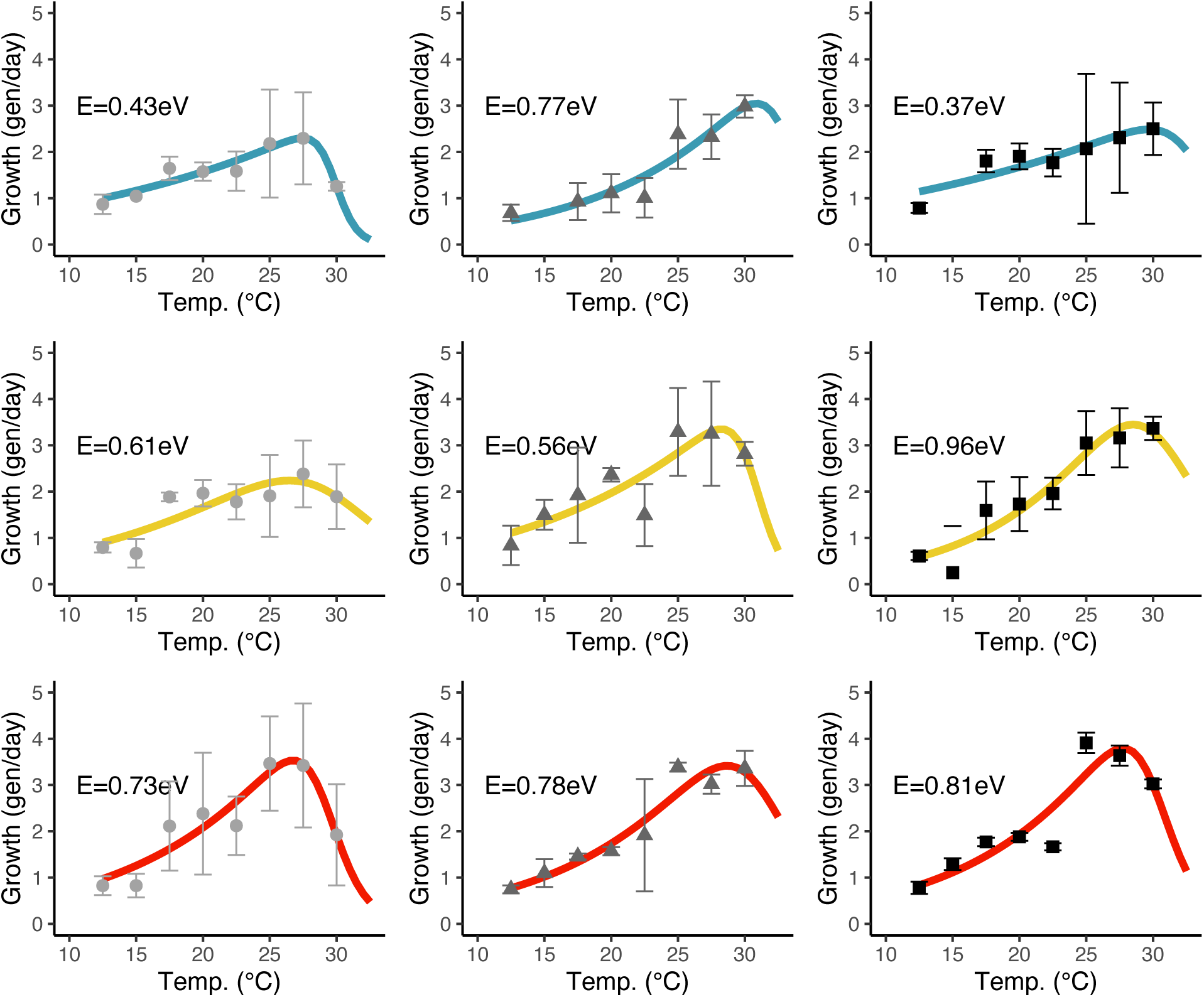
Thermal response curves for population growth measured after a few generations of acclimation to the new temperature. Each panel corresponds to one acclimation condition (same order as in fig. 12). Markers represent the mean across four replicates and error bars are 1 standard deviation. Continuous lines represent fitted Sharpe-Schoolfield curves (*rTPC* package version). Order of the plots and colour codes as in the previous figures.

**Figure 18:**
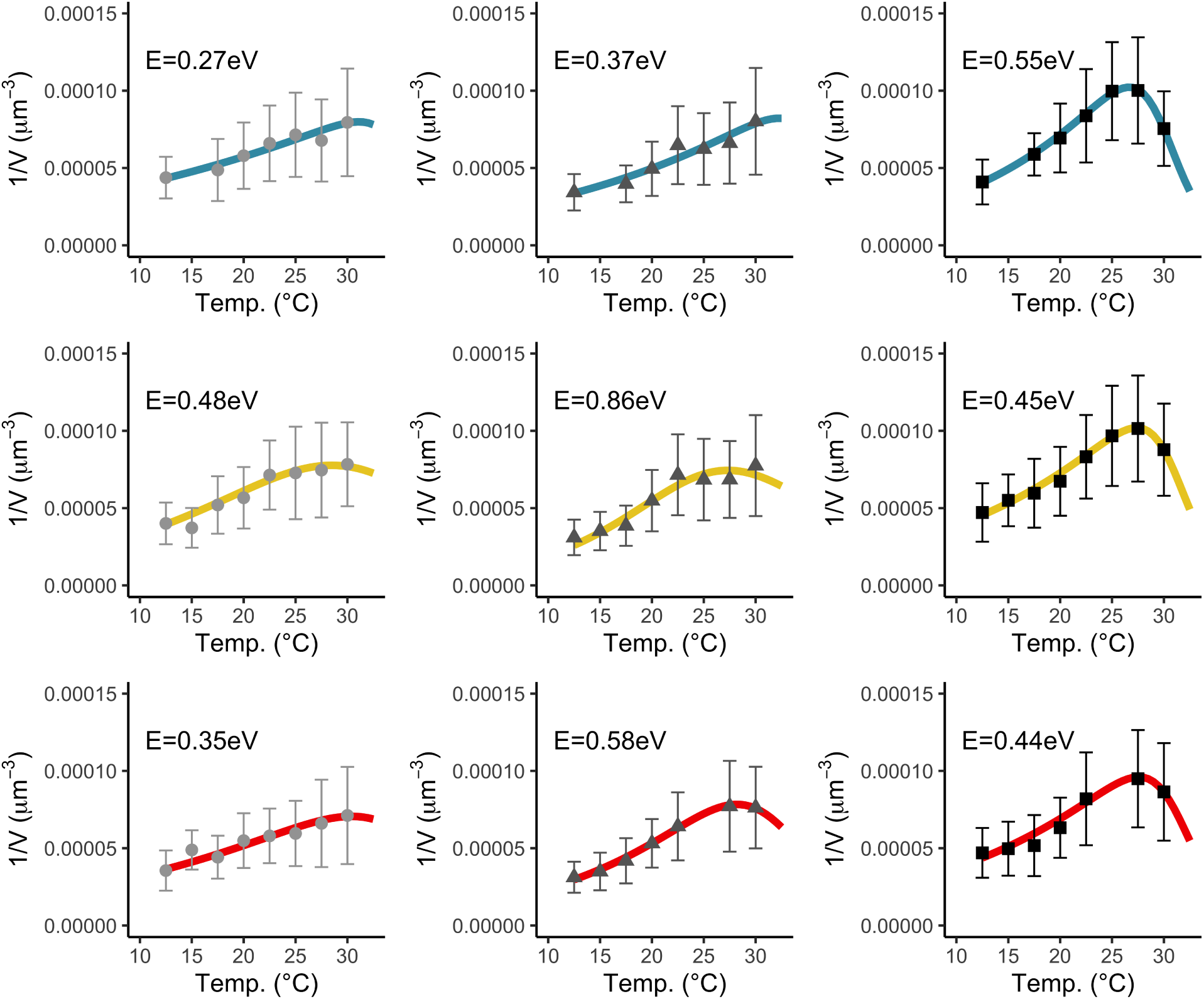
Thermal response curves for the reciprocal of cell volume. Each panel corresponds to one acclimation condition (same order as in fig. 12). Each data point in the figure represents the mean of 1/*V* (estimated by fitting ellipses on the 2D images of *Tetrahymena* and calculating the volume of the corresponding 3D spheroid) across all measured cells (where each cell is a ‘particle’ in the video-tracking). Error bars are 1 standard deviation. Continuous lines represent fitted Sharpe-Schoolfield curves (*rTPC* package version). The text on each graph reports the value of activation energy *E* for the Arrhenius part of the curve.

#### 2.6 Thermal response curves for cell volume (medium-term exposure)

Cells moved to a different temperature started changing their volume relatively rapidly after they were transferred to the new environment. These changes in volume are already well visible over the timescale used for the medium-term exposure experiments that were also used for measuring changes of population growth (sec. 2.5; see also below, sec. 2.10 for more details on the timescale of change).

Because cell volume decreases with temperature (opposite to most rate variables), in figure 18 we analyse the reciprocal of volume 1/*V*, so that thermal performance curves can be compared following the same analysis framework.

Already over the short timescale of this medium-term exposure to different temperature conditions, differences in cell volume are well detectable, with cells exposed to low temperature generally reaching larger volumes (lower values of 1/*V*). The curves app-pear to be non-monotonic, i.e. there is some evidence that the reduction of cell volume stops, and could also be reversed, above 28-30 °C.

#### 2.7 Speed vs. medium concentration

While our experiments only focus on three possible values of growth medium concentration, we would like to report here also the results of two preliminary experiments aimed at testing how medium concentration modulated the movement speed, and the population growth rate of *Tetrahymena*. These experiments were performed on different populations of the same strain of *Tetrahymena pyriformis* 1630/1W, obtained from the same supplier (CCAP).

In the first experiment we measured the movement speed of *Tetrahymena* acclimated to a wide range of growth medium concentrations, including more diluted culture media than those used in our main experiment. The data are presented in fig. 19, which plots the movement speed of cells acclimated to different environments. Environments differed in temperature (15 °C, 20 °C, 25 °C) and in the concentration of culture medium, with a large number of concentrations used, ranging from 0 % to 200 %. The incubation period was different for each temperature to reach a similar number of cell generations: 1.9 days at 25 °C, 3 days at 20 °C and 7 days at 15 °C. In all temperature conditions, speed was minimal when resources were scarce, increased to a maximum for intermediate concentrations, and then plateaued or slightly decreased at higher concentrations.

**Figure 19:**
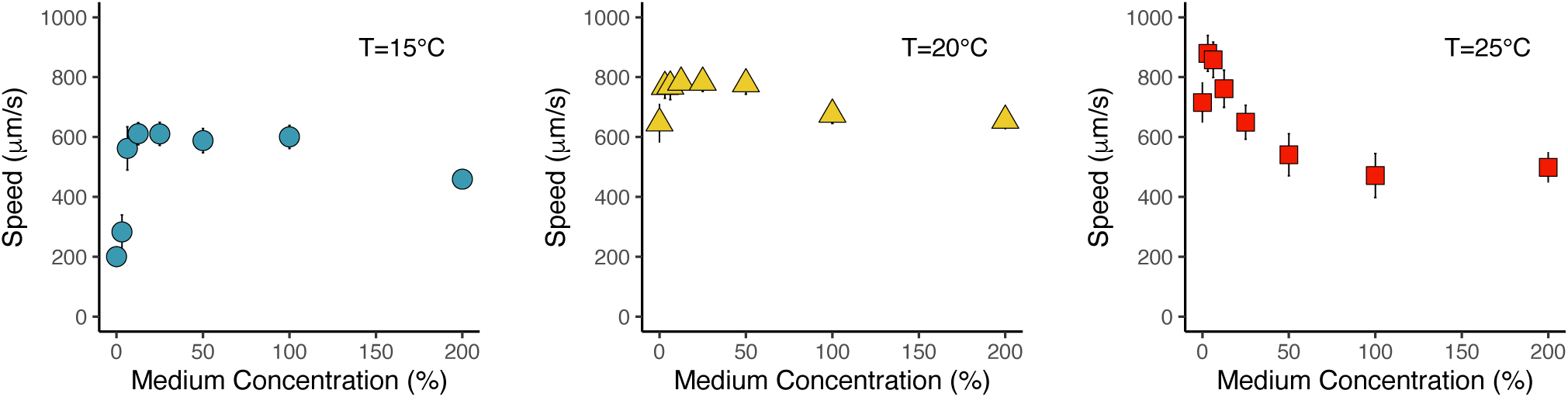
Results of a preliminary experiment looking at movement speed vs. the concentration of the culture medium. Each panel corresponds to one temperature condition. For each condition of medium concentration and tested temperature, we retain the 20% faster moving particles. Markers represent the mean across all measured speed values from the retained particles (pooling together particles from the four replicates) and error bars are ±1 standard deviation.

#### 2.8 Effects of medium concentration on population growth

In a separate preliminary experiment, we tested how the concentration of the growth medium affected the population growth of *Tetrahymena*. To this end, an initial population of *Tetrahymena pyriformis*, strain 1630/1W, initially cultured at 20 °C and 100 % culture medium (20 g/L Proteose peptone + 2.5 g/L yeast extract) was split across a range of different culture conditions, with temperature ranging from 10 °C to 30 °C and culture medium concentrations ranging from 0% (pure water) to 400% (4 times the density of standard growth medium). Cells were kept growing in the new condition, in three parallel replicate lines, for a variable number of days in order to approximately achieve a similar number of generations in each condition (ranging from 1.8 days at 30 °C, to 7 days at 15 °C and 8 days at 10 °C); after this culture period, population densities were counted, and population growth rates were estimated.

Figure 20 reports the results of this experiment, showing that both temperature and growth medium concentration affect the growth rate of *Tetrahymena*. One possible way to model the effects of culture medium concentration on population growth is by fitting a Michaelis-Menten equation to the population growth data (inspired by ref. Rubalcaba et al. (2020)). Specifically, we modelled the *per capita* growth rate of *Tetrahymena* (*G* in the equation 8 below) as a function of medium concentration [*C*] through the following equation:

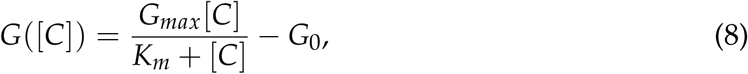

**Figure 20:**
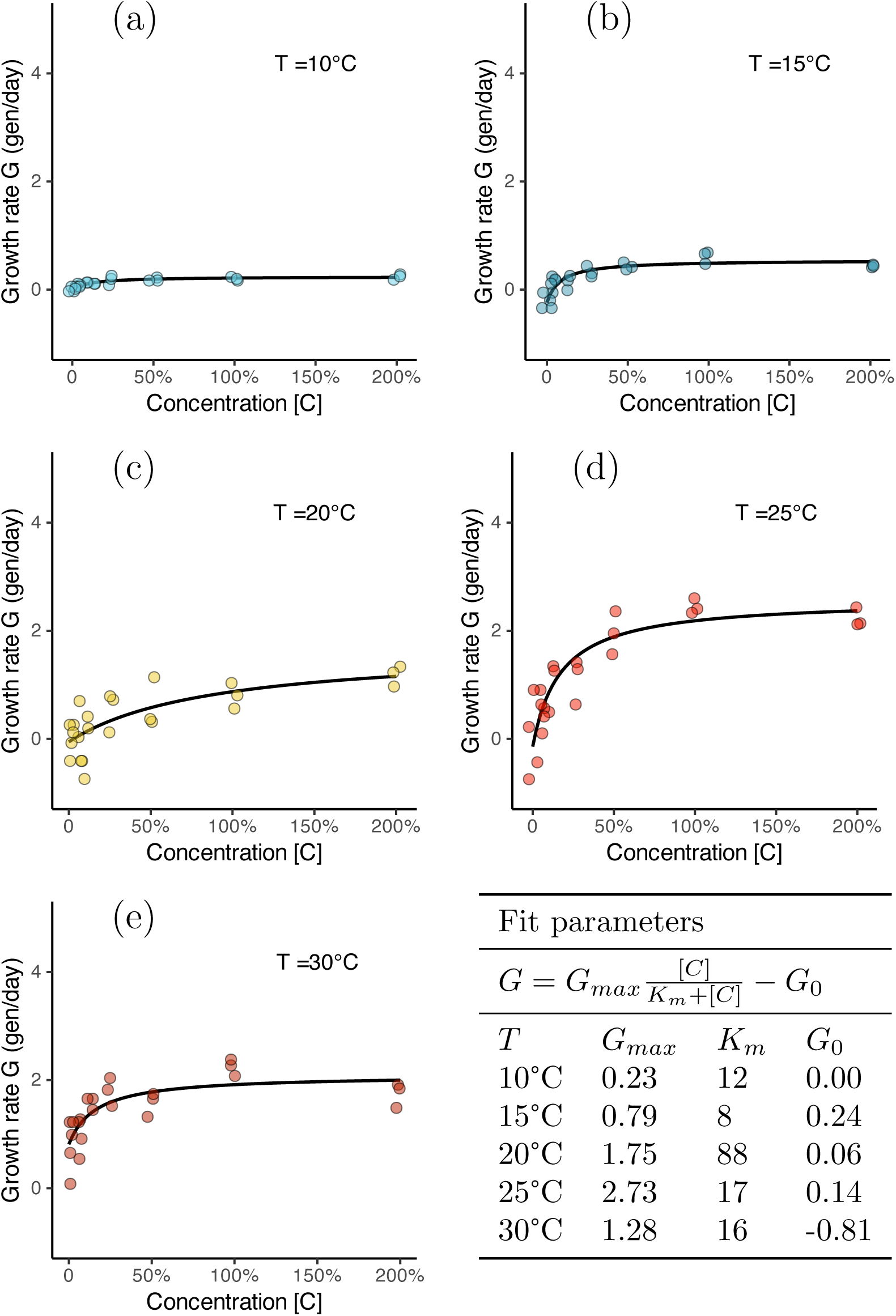
Effect of medium concentration on population growth. Each panel represent a different condition of growth temperature.

where *G_max_*is the maximum growth rate (also in generations per day), *K_m_* is the concentration of growth medium at which *per capita* growth is 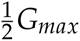, and *G*_0_ is a mortality rate, introduced here to account for the expectation that, in the absence of food, population growth should be negative.

This type of analysis confirms that temperature affects the maximum growth rate of *Tetrahymena* (with *G_max_*being maximum at 25 °C), and that the growth rate of *Tetrahymena* is not strongly limited by the concentration of the growth medium in the range of concentrations used in our main experiment (values of *K_m_* are small). The negative value of *G*_0_ estimated at 30 °Cmay seem counterintuitive, as it implies positive growth even in the absence of resources. However, this likely reflects cell division without net biomass production. As observed elsewhere in our study, cells transferred from lower to higher temperature reduce their cell volume, most likely by undergoing cell divisions. In this case, apparent population growth does not correspond to an increase in total biomass.

#### 2.9 Effects of oxygen concentration on cell size and shape

In our main experiment, we modulated the concentration of resources available in the growth medium in order to study the phenotypic changes of *Tetrahymena* in response to their environment. It is well known, however, that oxygen concentrations – possibly even more than nutrient resources – can play an important role in mediating the temperature-size rule of organisms (i.e. the variation in the typical body size of organisms which depends on the temperature of the environment in which they develop), and could also be responsible for other temperature-induced phenotypic changes (see e.g. Frazier et al., 2001; Hoefnagel and Verberk, 2015; Verberk et al., 2011, 2021; Rubalcaba et al., 2020).

Here, we report on a preliminary experiment in which we cultured axenic populations of the same strain of *Tetrahymena pyriformis* in an hypoxic and in an hyperoxic environment for 15 days. Cells were cultured in standard 100 % Proteose peptone – yeast extract medium in culture flasks, similar to other experiments that we describe in this manuscript. In this experiment, however, the flasks were stored in sealed boxes with an altered ratio of nitrogen and oxygen, mixed in different proportions to reach approximately 50 % of the standard atmospheric oxygen concentration (in the anoxic treatment) or 200 % (in the hyperoxic treatment). The boxes had to be opened every 2-3 days for subculturing *Tetrahymena* into fresh culture medium, and oxygen concentrations were rapidly restored after each subculture.

At the end of the acclimation treatment, we measured the size and shape of *Tetrahymena* cells from microscope images, measures that are reported in fig. 21. The plots in the figure confirm that cells cultured at low temperature were typically larger in size than cells cultured at high temperature. In contrast, exposure to low or high oxygen concentration within the range 50 % – 200 % did not show a detectable effect on cell size and morphology within this range.

**Figure 21:**
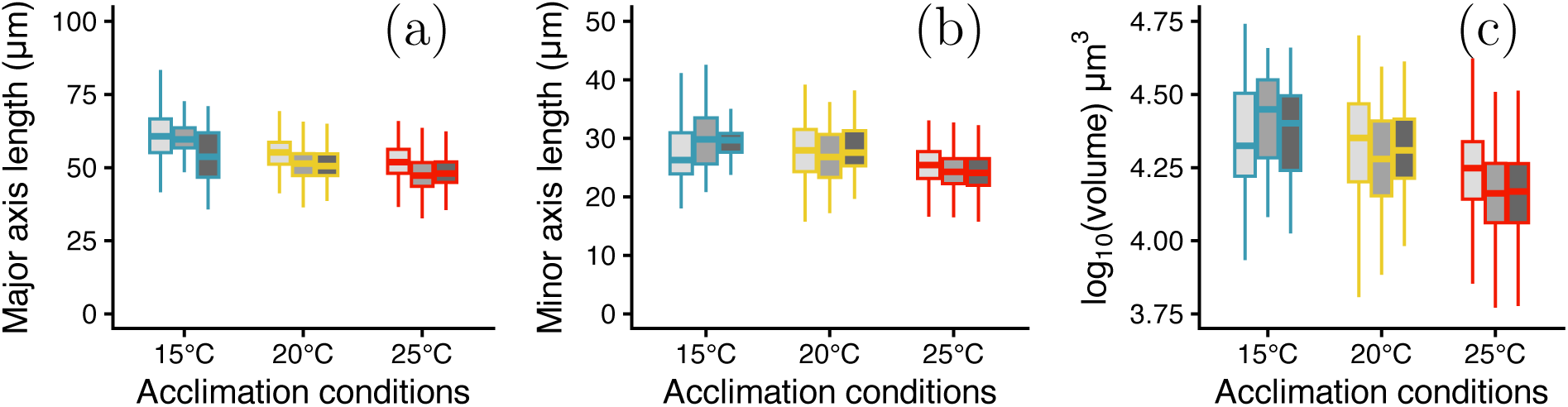
Effect of oxygen availability on cell size and shape. (a) Major axis length (approximately equivalent to cell length). (b) Minor axis length (cell width). (c) Cell volume. Light-shaded boxes (on the left for each temperature condition) indicate the anoxic treatment, middle boxes indicate normoxia, dark-shaded boxes indicate hyperoxia.

#### 2.10 Timescale of changes of cell volume and speed

Our measurements of *Tetrahymena* cell size and movement speed upon a ‘medium-term’ exposure to various conditions of temperature provide an opportunity to test the timescale over which phenotypic changes are produced. This section presents a breack-down of these data as a function of the amount of time (of the order of hours or days) spent at the new temperature conditions.

##### Cell volume changes over time during the ‘medium-term exposure’ to different temperature conditions

Figures 22, 23, and 24 present a re-analysis of the data that describe the thermal response curve for cell volume (the curves presented in figure 18). To collect these data, cells from the nine acclimation conditions (3 × temperature and 3 × medium concentration) were moved to a different temperature condition (in the range 12.5 - 30°C). Cell volume was measured repeatedly on different days after the transfer to the new temperature, and the re-analysis presented here reports the mean and standard deviation of volumes measured on each day. The baselines indicated by flat horizontal lines mark the starting volume in each condition (the volume of cells in the original culture before the transfer to the new temperature conditions).

**Figure 22:**
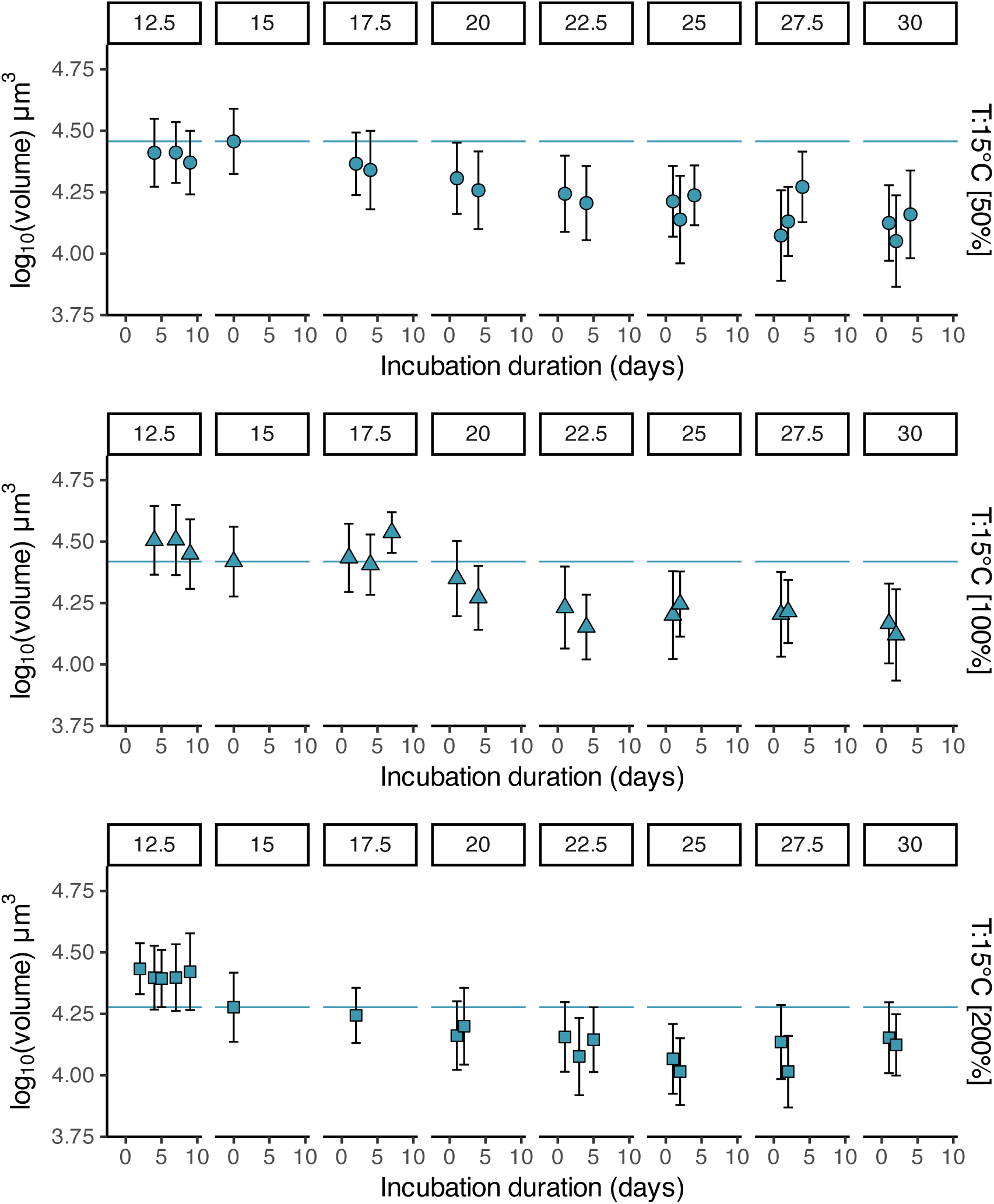
Body size change over time – acclimation temperature 15 °C. Rows correspond to different values of medium concentration; columns correspond to different test temperatures. Markers and errorbars indicate the mean standard deviation across all measurements on a given day of incubation; independent replicate lines were merged together. The blue lines indicate the mean volume of the original cell lines.

**Figure 23:**
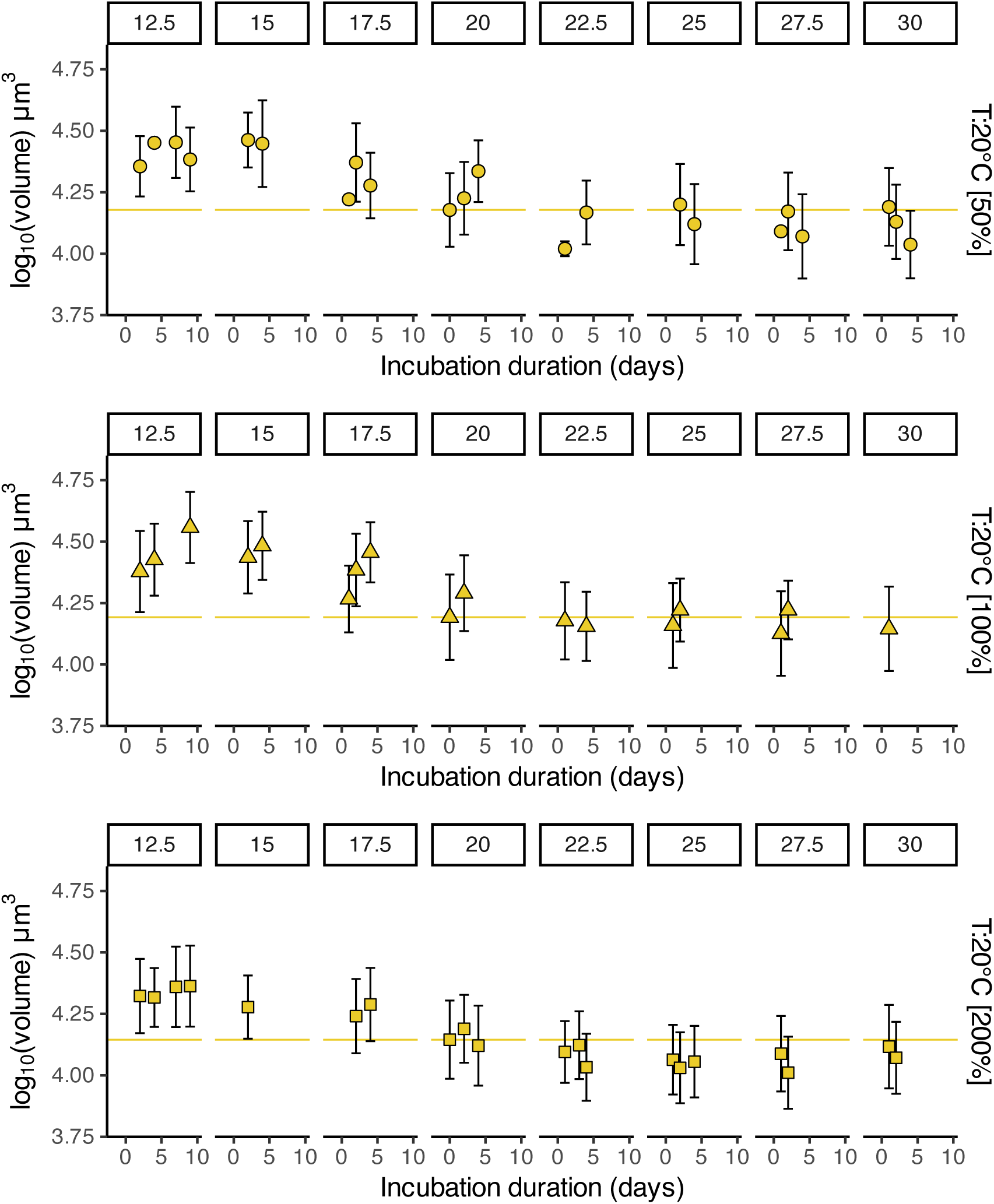
Body size change over time – acclimation temperature 20 °C. Rows correspond to different values of medium concentration; columns correspond to different test temperatures. Markers and errorbars indicate the mean standard deviation across all measurements on a given day of incubation; independent replicate lines were merged together. The yellow lines indicate the mean volume of the original cell lines.

**Figure 24:**
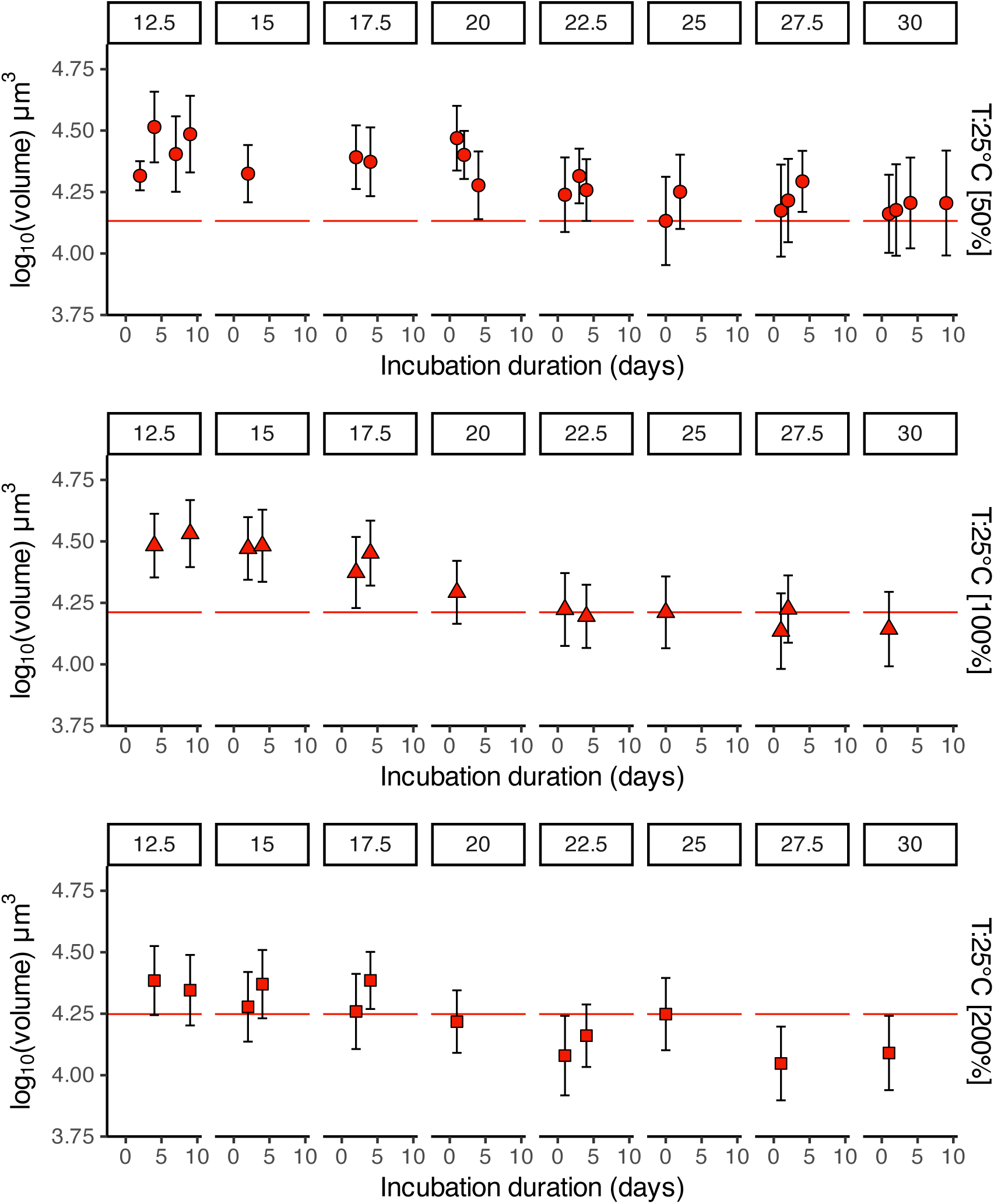
Body size change over time – acclimation temperature 25 °C. Rows correspond to different values of medium concentration; columns correspond to different test temperatures. Markers and errorbars indicate the mean ± standard deviation across all measurements on a given day of incubation; independent replicate lines were merged together. The red lines indicate the mean volume of the original cell lines.

In the figures we can see that the volume measured in each condition deviated from the baseline (with typically larger cells at low temperature and smaller cells at high temperature). However, measurements over different days were similar to each other. These results indicate that the change in volume occurred rapidly, before the first post-transfer measurement at the new temperature was taken, and possibly already at the first cell division after exposure to the new temperature. These observations suggest that cell size regulation might be triggered very soon after the change of temperature, and operate within the first few hours. Unfortunately, we did not characterise this phenomenon in detail, e.g. by measuring cell volume and the proportion of cells undergoing mitosis at short regular intervals after the change of temperature: this is something that could be explored in more detail in a future study.

Incidentally, we did not observe a ‘reverse temperature-size rule’ (i.e. cells temporarily exhibiting larger size at high temperature) as reported by DeLong et al. (2017) for *T. thermophila*. We can speculate that this discrepancy is due either to differences between the two *Tetrahymena* species, or to some synchronization of cell cycles in the cited reference when cells were moved from the stock culture to the experimental growth conditions leading to non-monotonic cell size changes over time in their experiment.

##### Speed changes over time during the ‘medium-term exposure’ to different temperature conditions

Figures 25, 26 and 27 present a re-analysis of the data that describe the thermal response curve for movement speed under a medium-term exposure (the curves presented in figure 4, right column, and Appendix figure 16). To collect these data, cells from the nine acclimation conditions (3 × temperature and 3 × medium concentration) were moved to a different temperature condition (in the range 12.5 - 30°C) and then movement speed was measured repeatedly on different days after the transfer to the new temperature. The re-analysis presented here reports the mean and standard deviation of speed measured on each day (note that, similar to all other measurements of speed, the speed values that we report are those from the 20% fastest moving cells, to avoid counting cells that might have been stuck against the microscope slide). The baselines indicated by flat horizontal lines mark the speed value in the acute speed response (analogous to the speed that would have been measured at the very beginning after the transfer to the new temperature). Measurements of speed deviated from those recorded during the acute responses, but measurements taken on different days remained similar to each other, suggesting that speed changed rapidly, already before the first ‘medium-term’ speed measurement at the new temperature was taken.

**Figure 25:**
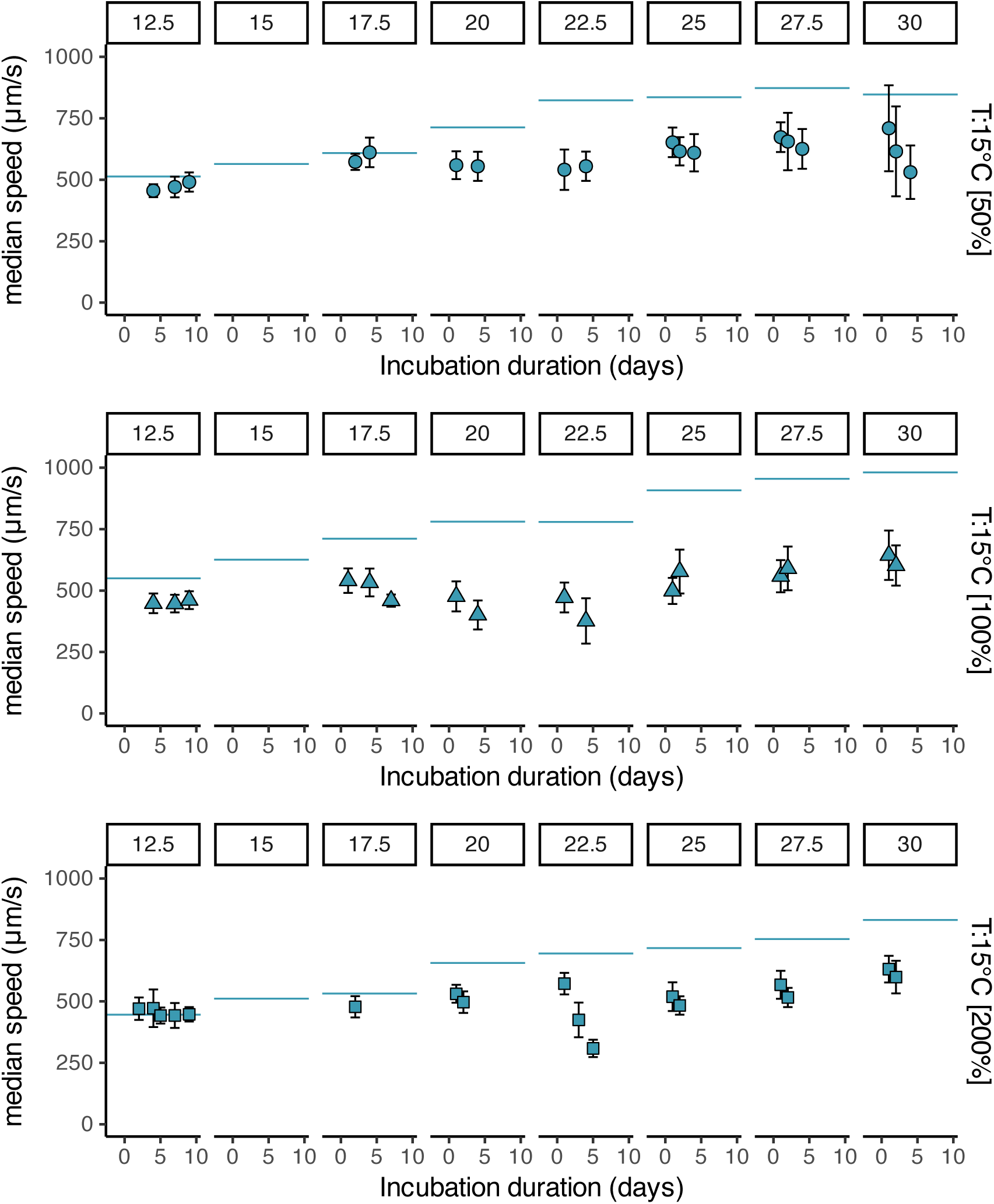
Speed change over time – acclimation temperature 15 °C. Rows correspond to different values of medium concentration; columns correspond to different test temperatures. Markers and errorbars indicate the mean ± standard deviation across all measurements on a given day of incubation; independent replicate lines were merged together. The blue lines indicate the mean speed value in the acute speed response (analogous to the speed that would have been measured at the beginning of this medium-term speed response).

**Figure 26:**
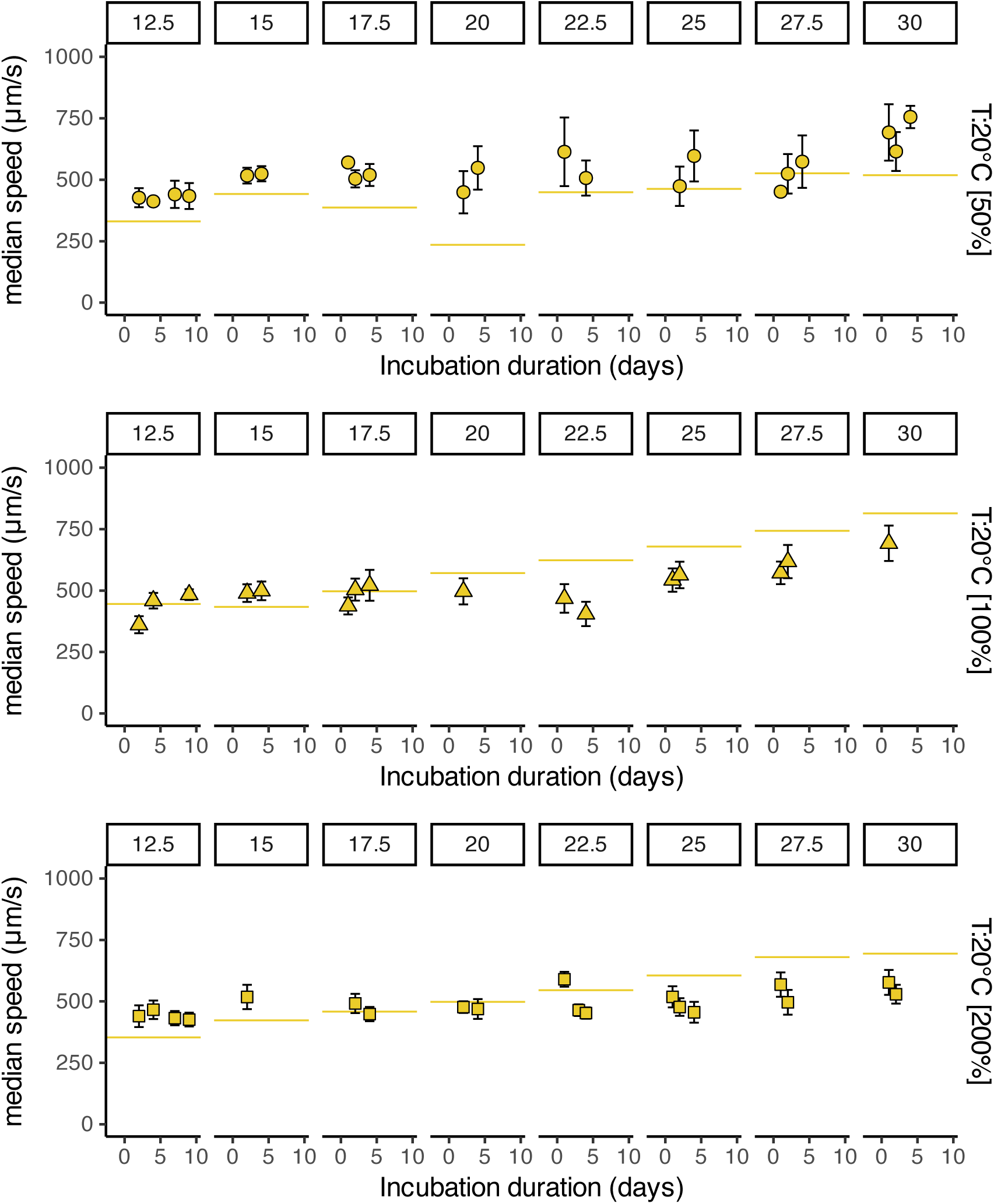
Speed change over time – acclimation temperature 20 °C. Rows correspond to different values of medium concentration; columns correspond to different test temperatures. Markers and errorbars indicate the mean ± standard deviation across all measurements on a given day of incubation; independent replicate lines were merged together. The yellow lines indicate the mean speed value in the acute speed response (analogous to the speed that would have been measured at the beginning of this medium-term speed response).

**Figure 27:**
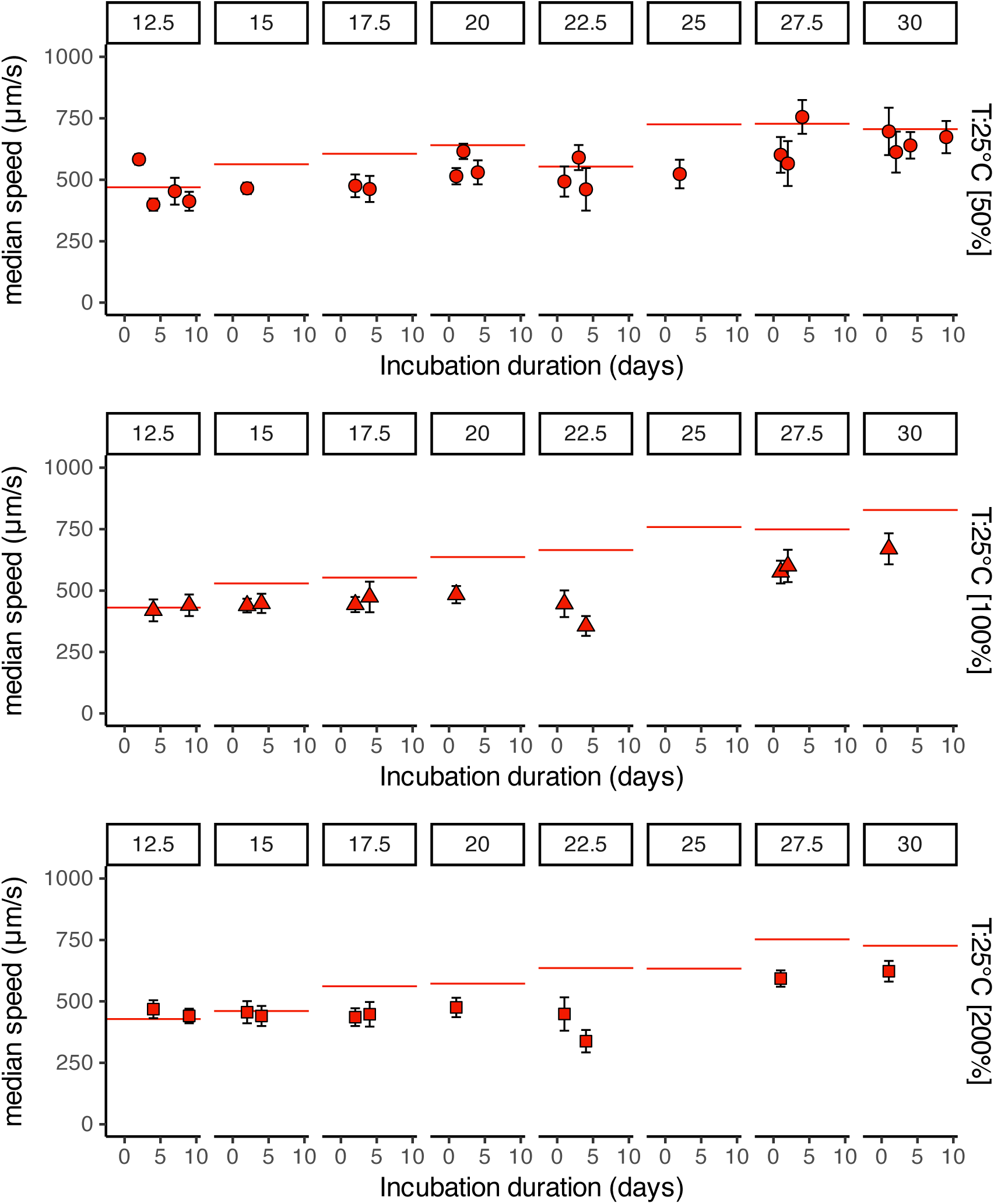
Speed change over time – acclimation temperature 25 °C. Rows correspond to different values of medium concentration; columns correspond to different test temperatures. Markers and errorbars indicate the mean ± standard deviation across all measurements on a given day of incubation; independent replicate lines were merged together. The red lines indicate the mean speed value in the acute speed response (analogous to the speed that would have been measured at the beginning of this medium-term speed response).

#### 2.11 Impact of cell size on movement speed

Larger organisms move typically faster than smaller ones. They also have a faster *per capita* metabolic rate, simply because of their size. The observation that, in our experiments, *Tetrahymena* cells changed their volume rapidly and substantially, depending on the experimental conditions, begs for the question: to what extent do the observed changes in cell volume alone account for the other observed changes in metabolic rate and movement speed?

To better understand the possible relation between cell volume and speed, we can consider the example of an isogeometric unicellular organism (an organism that could have different sizes but which preserves the same shape): its metabolic rate – and total available power – would scale as the volume of the cell, or as the third power of the linear dimension (assuming a linear scaling between metabolic rate and cell volume in unicellular eukaryotes; DeLong et al., 2010), and this metabolic power would have a positive impact on speed; the viscous drag, having a negative impact on speed, would scale as the linear dimension (e.g. eq. 3, expanded in Appendix eq. 13). If we further assume that a constant fraction of metabolic energy is allocated to movement, then speed *U* depends on the invested power *P* as *P* = *sU*^2^ (eq. 3). Here, *P* is proportional to *V* and *s* is proportional to the linear cell dimension (*s* ∝ *V*^1/3^). Combining these relationships leads to the prediction: *U* ∝ *V^α^* with *α* = 1/3. (If, instead of assuming a linear scaling of metabolic rate with cell volume we assumed a 3/4 power scaling (as in Kleiber’s law), the same reasoning would lead to a predicted relationship between movement speed and cell volume as *U* ∝ *V^α^*, with *α* = 5/24).

Based on these scaling arguments, we could predict that a change by a factor 2 in the cell volume would translate into a factor 2^1/3^ ≈ 1.26 in the change of speed (or 2^5/24^ ≈ 1.16 if we assumed a 3/4 power scaling of metabolic rate with cell volume).

Inevitably, this isogeometric unicellular organism is only a very crude approximation of our *Tetrahymena*. Among the limitations of this approximation we could mention the fact that the beating cilia themselves responsible for cell propulsion cover the surface of the cell, and their number would scale as the square of the linear cell dimension, possibly affecting the fraction of metabolic energy allocated to movement. The assumption that a constant proportion of metabolic energy is spent for fuelling movement is probably also simplistic. Nevertheless, the scaling arguments presented above provide a starting point against which the empirical data can be compared, until more refined models can eventually be formulated. Let’s now have a closer look at the scaling relations between cell size and movement speed that can be observed empirically from our own data.

One way to explore the relationship between movement speed and cell volume, excluding the confounding effects of other variables such as temperature and available resources, is by exploiting the intrinsic variability of cell sizes in our experimental populations: in each culture condition there were smaller and larger cells – partly related to their stage of growth in-between mitotic divisions. Irrespective of their different sizes, these cells were otherwise all exposed to identical culture conditions. We re-analysed the data that are summarised in each data point of figure 14 by plotting the logarithm of movement speed as a function of the logarithm of cell volume. We excluded from this re-analysis data in which cells were tested in the deactivation part of their thermal performance curve (i.e. at 32.5 °C or higher temperature) because we were concerned about possible confounding interactions between cell size, shape, metabolic rate and speed (for example, we anecdotally observed that cells exposed to temperatures higher than 30 °C changed their shape as an acute response to temperature), and for even higher temperatures of 37.5 or 40 °C speed was not even constant over time (figure 10).

Figure 28 presents an example of the plots resulting from this re-analysis, by showing the speed-volume relationship for cells acclimated at 20 °C and 100 % medium concentration, for each value of temperature at which cells were tested (i.e. for the data in fig. 14, middle panel). The curves depict a generally positive trend of increasing speed with increasing cell volume. The slopes of log *U* = *α* log *V* + *K* fitted in a least squares sense across all the 81 conditions that correspond to the activation part of the *Tetrahymena* thermal response curve gave a value of *α* = 0.36 ± 0.12 (mean ± s.e.), not dissimilar from the theoretically predicted value *α* = 1/3.

**Figure 28:**
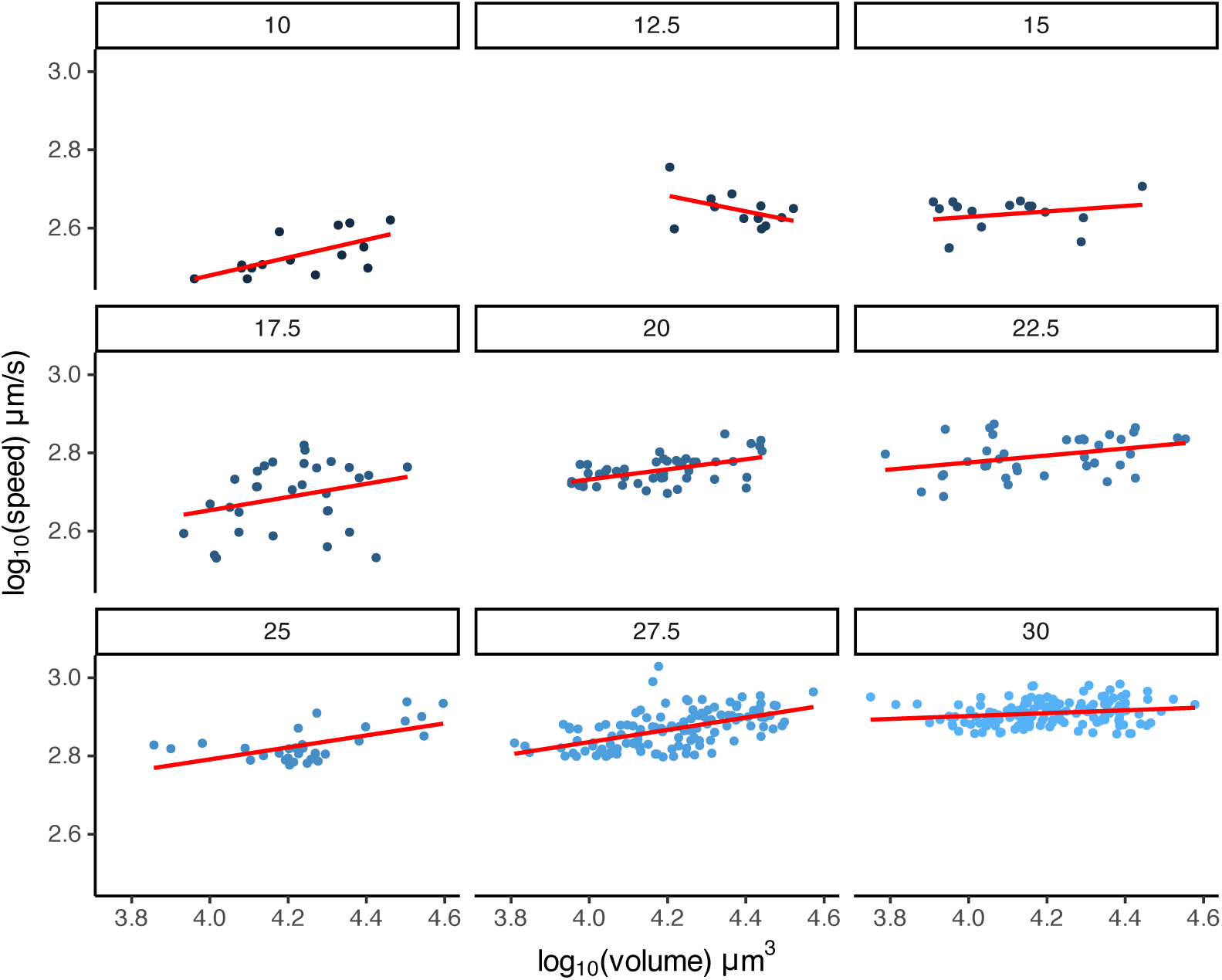
Cell speed as a function of cell volume for cells cultured at 20°C and 100 % medium concentration and tested at the same medium concentration and at different temperatures.

### 3 Additional details on theoretical arguments

The energy that individual organisms (here, individual *Tetrahymena* cells) can invest into fuelling reproduction and population growth is the result of a balance between energy intake from feeding *F*, and energy spent for basal metabolism (maintenance) *B*, and for movement (propulsion) *P*:

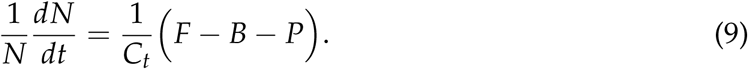

In the equation, the left-hand side represents the *per capita* growth rate of the population (*N* is the population size – the number of cells – and 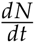 is the growth rate of the entire population), *F*, *B*, and *P* have units of power (e.g. nanoWatts) and *C_t_* can be thought of as the energy amount required to form a new cell. (In natural populations, energy can also be lost to predation and to other causes of mortality, which are assumed absent in our axenic populations of *Tetrahymena* maintained in exponential growth).

The dependence of the individual variables *F*, *B*, *P*, and *C_t_* on the characteristics of the organism or of the environment can further be made explicit to estimate how these variables are passively affected by environmental conditions over short timescales, and to predict how the organism might actively modulate them through acclimation or adaptation over longer timescales.

#### Basal metabolic rate

The basal metabolic rate *B* is known to depend both on temperature *T* and on cell volume *V* (relative to a reference volume *V*_0_):

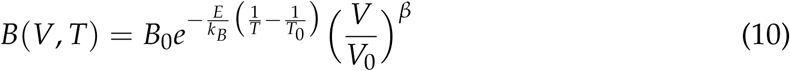

where *B*_0_ is the value of metabolic rate measured at the reference temperature *T*_0_. The dependence on temperature follows the Arrhenius equation introduced above (eq. 6) and described by the activation energy *E* (with *k_B_* the Boltzmann constant, *T* the environmental temperature expressed in Kelvin, and *T*_0_ the reference temperature. The scaling of metabolism with volume *V* (here expressed as relative to a reference volume *V*_0_) follows an allometric relation with exponent *β* Gillooly et al. (2001); Brown et al. (2004).

#### Movement

Swimming entails an energetic cost, associated with overcoming the drag of water (neglecting other costs associated with movement, such as the increased risk of predation, see Visser (2007)). For small organisms swimming at low Reynolds numbers, the power *P* required for swimming at speed *U* is

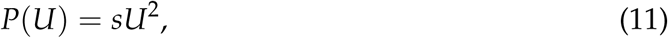

where the factor *s* accounts for the energy dissipation due to drag, and to the conversion of metabolic energy to movement, and it can be expanded as

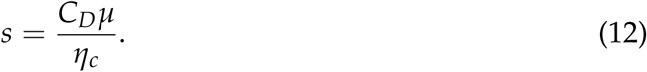

Here, *C_D_*is the drag coefficient, *µ* is the dynamic viscosity of the culture medium, and *η_c_* is the efficiency of conversion of metabolic energy into movement.

Approximating *Tetrahymena* as a prolate spheroid that moves along its long axis, the drag coefficient itself *C_D_* is given by the approximate formula:

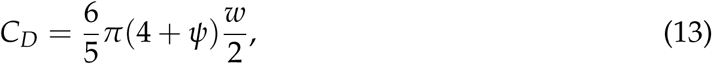

where *w* is the cell width and 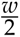 the small radius (or semi-minor axis), while *ψ* is the elongation (ratio of length over width, 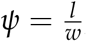; see Clift et al., 2005).

The parameter *µ*, which expresses the dynamic viscosity of water is in turn also dependent on salinity and on temperature: it decreases at high temperature, leading to a small reduction of drag with increasing temperature. In our analyses we calculate viscosity with the following version of Andreade equation,

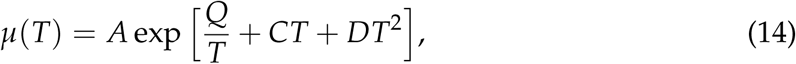

where the parameters take values: *A* =1.856 × 10^−14^ g/(m s), *Q* = 4209 K, *C* = 0.04527 K^−1^, *D* = −3.376 × 10^−5^ K^−2^.

Finally, the parameter *η_c_* in equation 12 quantifies the efficiency of energy conversion in the cilia, and accounts for the fact that only a fraction of the metabolic energy allocated to movement results in propulsive power. Ciliary propulsion is not efficient and values of *η_c_* calculated for *Paramecium* are for instance of the order of 0.078 % Katsu-Kimura et al. (2009). We do not have a measure of *η_c_* for *Tetrahymena* but, based on the typical values of relevant quantities that we measured in our experiments, a value of *η_c_*as small as 0.02 % would not be unrealistic (considerations presented in the Appendix, section 3).

#### Feeding

All energy acquisition is associated with feeding. We assume a Holling type II functional response in which the encounter rate with food is proportional not just to food density, but also to the encountered volume of medium or, equivalently, to the movement speed of *Tetrahymena*. This is described by the following equation for the feeding rate *f* (*U*):

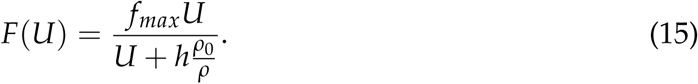

Here, *U* is the movement speed of *Tetrahymena*, *h* is the ‘half saturation speed’ or the speed at which the feeding rate reaches half the maximum value, *f_max_*is the maximum feeding rate (which in our system might be limited by digestion time more than by prey handling). Finally, *ρ* is the concentration of resources in the environment – in our case, the concentration of the growth medium – always measured in relation to a standard medium with concentration *ρ*_0_ (in our experiments, *ρ*_0_ is the concentration of the 100 % growth medium).

It should be noted that the functional forms of all the equations presented here, from 2 to 4, represent choices among multiple possible alternatives. For example, equation 4 assumes a direct encounter model for food – food encounters increase linearly with speed – while alternative models, e.g. based on diffusional deposition (see e.g. Karp-Boss et al., 1996) might also be relevant in our system. Similarly, equation 12 assumes that the efficiency of conversion of metabolic energy to propulsion *η_c_*is not itself dependent on temperature. Because of these and many other assumptions in the formulation of the model, and because of the uncertainty in the empirical values of some parameters (some that we did not measure directly in our experiments, such as feeding rates, and some that can change over time, such as movement speed and the volume of *Tetrahymena* cells), we intend the modelling presented here as a tool to gain qualitative insight rather than a precise quantitative fitting of empirical data.

An additional complexity is represented by the fact that feeding in *Tetrahymena* is not only limited by prey encounter rate (or equivalently, volume clearance rate for a system like our experimental system in which ‘prey’ is represented by dissolved proteins), but also by capture rate. Encounter rate can be assumed to be determined by movement speed, driven by the activity of somatic cilia, while capture rate also depends on the activity of oral cilia surrounding the oral apparatus. While this distinction between prey encounter and capture introduces an extra complexity, we note that both oral and somatic cilia are likely to experience similar hydrodynamic constraints (e.g. power costs scale proportionally to *U*^2^).

#### Complete model

Table 6 presents a full version of the model that describes *Tetrahymena* population growth as a function of the different items of the energy budget of cells.

**Table 6:**
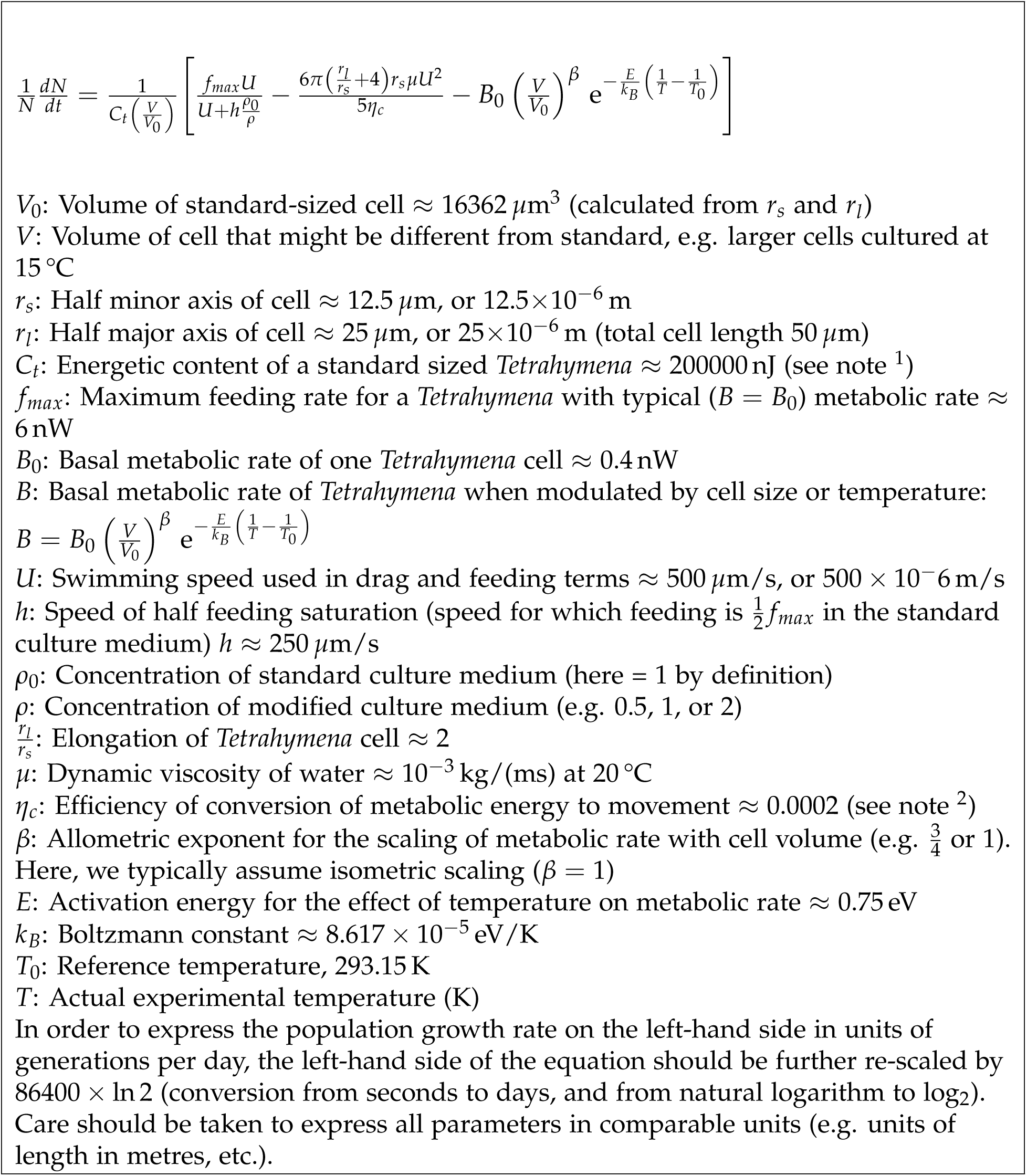
A possible full expansion of eq. 1 presented in the main text, alongside with tentative values of parameters.

#### 3.1 Derivation of optimal swimming speed

The optimal foraging speed of *Tetrahymena* is the foraging speed that maximizes *per capita* population growth. Here we calculate it for the ‘simplified’ model (that aggregates some of the variables):

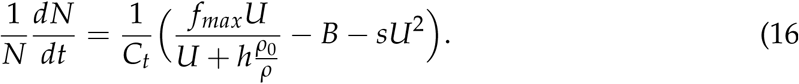

*Per capita* growth is maximizes when the derivative of the energy balance with respect to speed is zero:

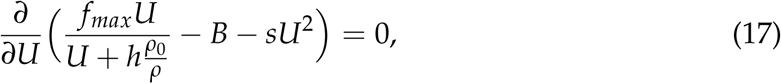

which gives

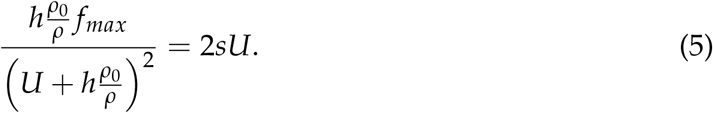

When solved for *U*, equation 5 gives the following expression for the optimal swimming speed *U*^∗^:

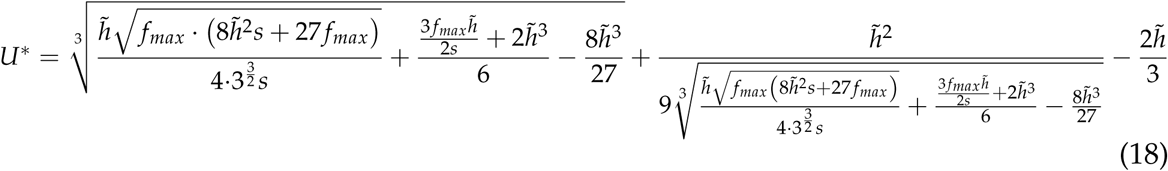

where we replaced 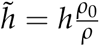 to simplify the notation.

As we can see in the derivation, the basal metabolic rate *B* does not depend on speed, meaning that optimal speed is only affected by metabolic rate through the effects of metabolism on the parameters of the functional response.

The optimal speed calculated with this formula is shown in figure 1 (for a cell of length *l* = 50 *µ*m, width *w* = 25 *µ*m, maximum feeding rate *f_max_*= 6 nW, ‘half saturation speed’ *h* = 250 *µ*m/s, and efficiency of conversion of metabolic energy to movement *η_c_* = 0.02 %). It can be interesting to note the qualitative similarity with the corresponding experimental data in figure 19 and fig. 4(i).

##### Predicted relationship between activation energies for respiration and movement speed

Here we derive a prediction for the activation energy for movement of a microorganism swimming at low Reynolds numbers under the assumption that the organism does not change its shape or size when it is exposed to a different temperature, that it allocates a constant fraction of its metabolic energy to movement, and that the efficiency of energy conversion within the ciliary layer is independent of temperature. This kind of reasoning has the double purpose:

1. Formulating an initial and general prediction for the scaling of movement speed with temperature in microscopic organisms. In this case, the prediction will be that the activation energy for speed should be substantially lower than the activation energy for respiration.
2. Providing a null hypothesis against which to compare the experimental data: deviations of data from the predicted values will be informative of additional factors that play a role in determining how *Tetrahymena* movement speed changes with temperature. For example, deviations from the expected values could indicate that one or more of the assumptions that we made above are violated (e.g. the fraction of metabolic energy allocated to movement might not be constant, or the efficiency of energy conversion might be dependent on temperature).

Our starting point are equations 3 and 12, which combined describe the relationship between power of swimming *P* and speed *U*, as a function of the drag coefficient *C_D_*, the efficiency of conversion of metabolic energy into movement *η_c_*, and the viscosity *µ* of the medium in which the microorganism moves:

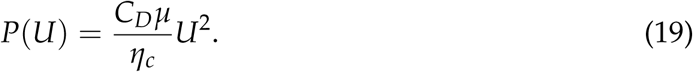

The drag coefficient *C_D_* only depends on cell size and shape, as in equation 13 (or in the corresponding, non-approximated equation 40). The dependence of viscosity *µ* on temperature is expressed by equation 14.

Looking at equation 19 we can see that if *µ*, *η_c_* and *C_D_* were independent of temperature, and *P* represented a fixed proportion of the total metabolic power, independent of temperature, then we would simply have *P* ∝ *U*^2^, and we would predict that the activation energy for speed *U* should be 1/2 the activation energy for metabolic rate. In practice, as water viscosity *µ* also changes with temperature (eq. 14), we can calculate the predicted dependence of speed on temperature also incorporating into equation 19 the calculated values for *µ*. For example, figure 29(a) shows a simulated thermal response curve for metabolic rate, and the corresponding modelled thermal response curve for movement speed (fig. 29(b)). A simulation across a range of values of activation energy gives the relationship in figure 29(c), which is well summarised by the equation *y* = 0.0868 + 0.487*x*, where *y* is the activation energy for movement speed and *x* is the activation energy for metabolic rate (respiration).

**Figure 29:**
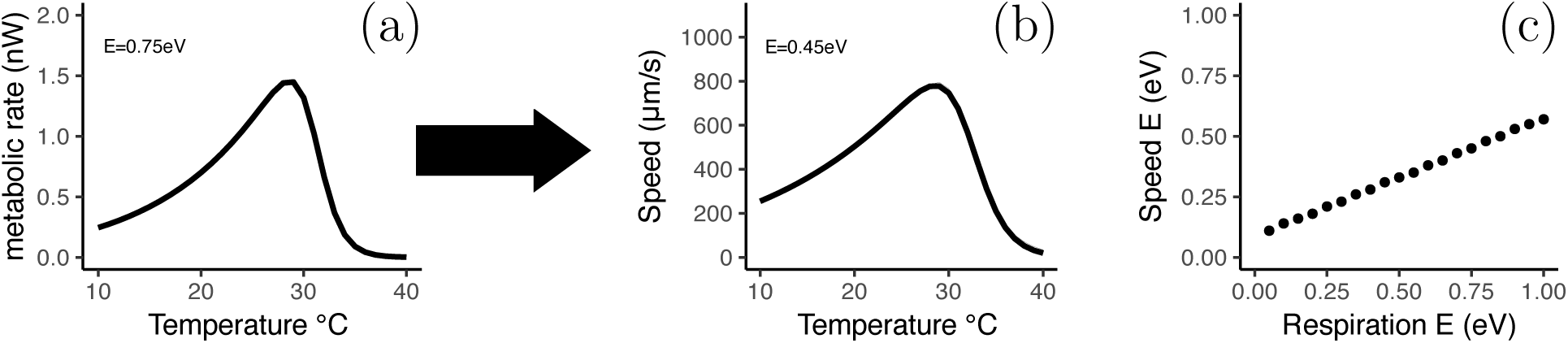
Modelled relationship between metabolic rate and speed in thermal response curves for short-term exposure. **(a)** Thermal response curve for metabolic rate simulated with a Sharpe-Schoolfield equation (eq. 7) with rate at reference temperature *B_V_*_0_ = 0.7 nW, activation energy *E* =0.75 eV, reference temperature *T*_0_=293.15 K, deactivation energy *E_d_* =7 eV, and high temperature *T_h_* =304.15 K. The thermal response curve for metabolic rate in (a) translates into the thermal response curve in panel (b) for movement speed once accounting for the viscous drag of moving at low Reynolds numbers and of temperature-dependent changes in water viscosity. Curve in **(b)** based on the assumption that cells move at 500 *µ*m/s when tested at the reference temperature of 293.15 K, and that the efficiency of conversion of metabolic energy into movement is *η_c_*= 1.0276 10^−4^, independent of temperature. The best-fitting value of activation energy is here *E* =0.45 eV, slightly larger than 0.75/2=0.375 eV because of the additional effect of temperature on water viscosity. **(c)** Predicted relation between activation energy for movement speed as a function of the activation energy for respiration based on the same parameters and assumptions as in the other figure panels.

#### 3.2 Oxygen availability to *Tetrahymena* cells

The reduced solubility of oxygen in water at high temperature has often been indicated as a possible factor underlying the temperature-size rule of ectotherms (the observation that many organisms tend to reach a smaller adult body size when they grow in a warmer environment; Verberk et al., 2021; Forster et al., 2012).

While we did not aim at studying specifically the role of oxygen in our experiments, it can still be interesting to consider briefly to what extent *Tetrahymena* growth could be limited by oxygen availability directly, rather than by the metabolic ability of cells to process oxygen and nutrients.

Oxygen solubility in water decreases with temperature, and is hence likely to become a limiting factor at high temperature. At 30 °C, O_2_ concentration in a saturated water solution would be about 252 *µ*M (see below, section 3.4, for details). Considering that, in our cultures, oxygen concentration was always very close to saturated values, could oxygen availability be directly limiting the growth of an organism the size of *Tetrahymena* in these conditions?

Without giving a complete answer to this question, we could consider a limiting case scenario, of a spherical cell of radius *r_o_* that does not move – meaning that all the oxygen consumption required to fuel its metabolism has to be provided through diffusion alone – and yet, whose metabolic rate is the same as that of an active *Tetrahymena* cell. The current *I* of oxygen to a spherical, stationary cell of radius *r_o_* can be calculated as described in section 3.3 below, and it is given by equation 31. We report the equation also here:

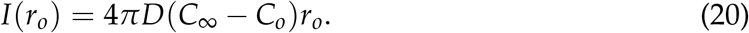

Here, *C*_∞_ is the concentration of oxygen in the culture medium away from the cell, which at 30 °C would be approximately *C*_∞_ = 252*µ*M = 0.252mol/m^3^. *C_o_* is the concentration of oxygen required at the position of the cell membrane in order for the cell to be able to function normally. Mitochondria are not limited by oxygen availability if the local oxygen concentration is above 10 *µ*M (based on Wilson et al., 1988). The diffusion coefficient (at 30 °C) for oxygen in water (see sec. 3.5) is *D* = 2.71 × 10^−9^m^2^ / s: If the cell diameter is 50 *µ*m and, consequently, *r_o_* = 25 *µ*m, we can calculate the current into the cell as follows (expressing all units of distance in metres):

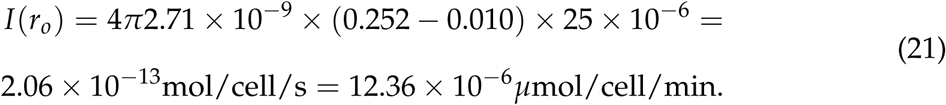

The fact that these values are one or two magnitude orders larger than the measured respiration rate of individual cells in our experiments, indicate it as unlikely that oxygen limitations played a major direct role in shaping *Tetrahymena* acclimation (also increasing the concentration of oxygen *C_o_* required on the cell surface by one order of magnitude, to 100 *µ*M does not change substantially this result).

The calculation above is intended to illustrate how we would estimate the supply of oxygen to a spherical cell for a specific value of temperature and of cell radius. The same calculations can be applied across the full range of temperature values used in our experiments and for a wide range of cell volumes in order to model oxygen supply to cells of different sizes and at different temperatures. In parallel, we can also model oxygen demand as a function of cell size and temperature based on eq. 7. We can then calculate the ratio, which we indicate with the Greek letter Φ, between oxygen supply and demand predicted for each combination of parameters of cell size and temperature.

The isolines of the supply/demand ratio Φ are presented in figure 30, in an analysis and visual representation inspired by Deutsch et al. (2022).

**Figure 30:**
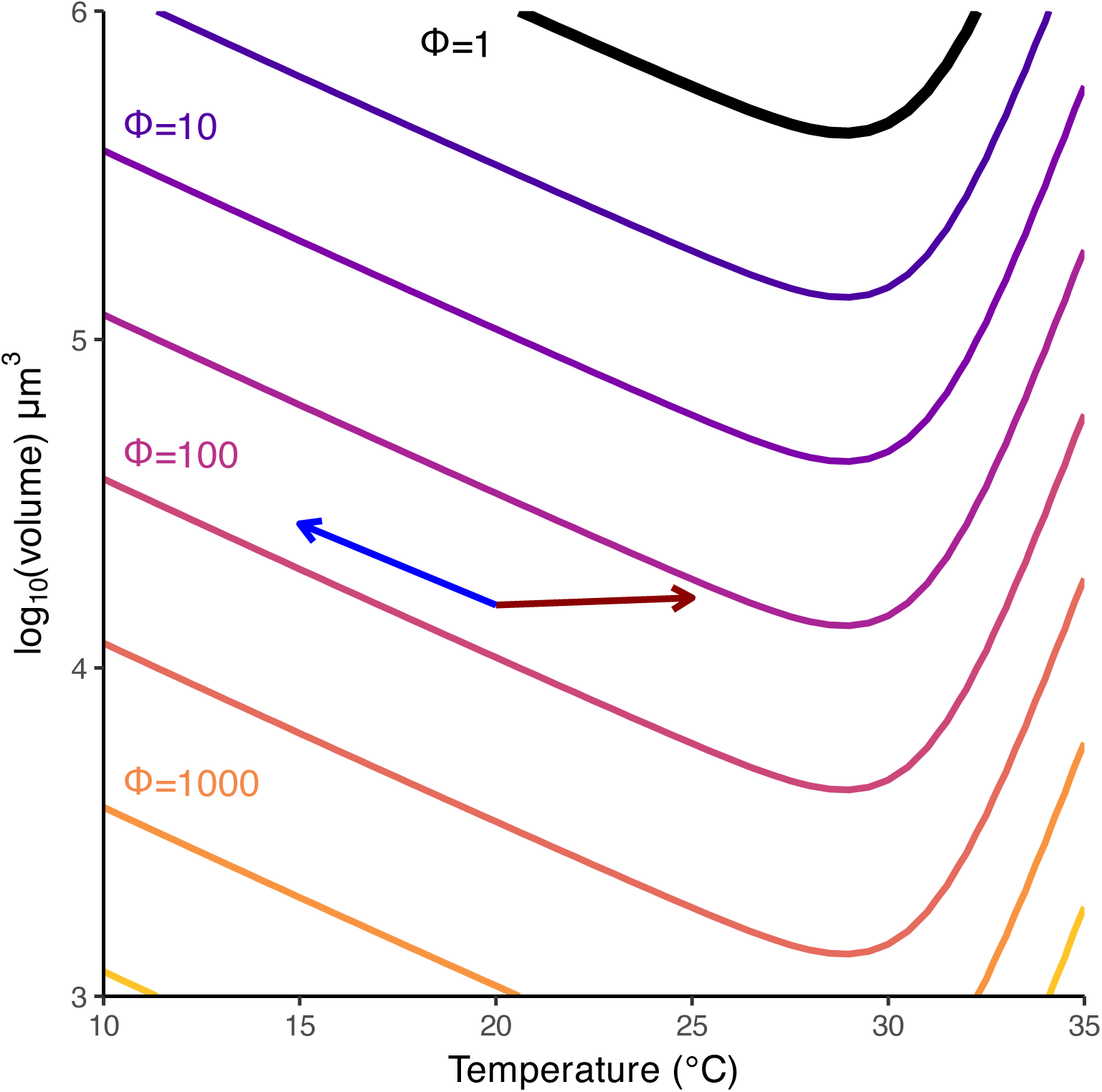
Supply and demand of oxygen. Demand of oxygen is calculated based on an activation energy of 0.89 eV and an oxygen/energy demand of 0.69 nW at 20°C. Oxygen supply is calculated based on the solubility and diffusion coefficients of oxygen in water, scaled with temperature, and assuming that the concentration required at the position of the cell membrane is 100 *µ*M. The arrows correspond to the changes of volume of *Tetrahymena* cells between 20°C (base of the arrow) and the other two main temperature conditions of 15 and 25°C (tips of the arrows). The curves of Φ indicate different ratios between oxygen supply and oxygen demand as a function of temperature and cell volume (large values of Φ indicate that supply is in excess with respect to demand). The black curve indicates the critical value of Φ = 1

In the figure, the value of Φ = 1 is marked by the thick black line near the top of the graph. For large cells, falling in the region above the thick black line, the supply of oxygen would be insufficient to meet the metabolic demand of the cell. Conversely, small cells, especially if they are cultured at low temperature (bottom-left part of the graph) are predicted to have an oxygen supply far in excess of metabolic demand. The volume of *Tetrahymena* cells in our experiments is marked by the two arrows in the figure, with the base of the arrows marking the volume of cells acclimated at 20 °C (and growth medium concentration 100 %) and the tips of the arrows marking the volume of cells acclimated at 15 °C and 25 °C, respectively for the blue and for the red arrow (medium concentration 100 %).

From the plot in figure 30 we can see that the change in cell volume observed for *Tetrahymena*, indicated by the arrows, is roughly aligned with the isolines of Φ. Other data on the volume change of *Tetrahymena* from our experiment, such as those at different medium concentrations (figure 2(c) in the main text), or those from the thermal response curves for volume change (e.g. fig. 18) would also present a similar alignment. The alignment of the cell-size changes of *Tetrahymena* with the isolines of Φ could suggest that cell volume changes are a mechanism that guarantees a constant oxygen supply-demand ratio to the cell. However, we should also note that the region of the graph that corresponds to the cell volume of *Tetrahymena* is very far from the thick black curve that marks the critical value of Φ = 1: no matter how cells in this region of the graph change their volume or their metabolic rate, oxygen supply will always be far in excess of demand!

##### An alternative approach to the study of oxygen supply and demand

In the type of modelling that we just described above we made a number of assumptions, which we would like to review here before we introduce an alternative modelling approach that has been followed in the scientific literature:

1. We assumed that cells were not moving. In reality, the fast movement of *Tetrahymena* means that the concentration of oxygen at the cell membrane is likely to be much higher (*C_o_*close to *C*_∞_), as the cell constantly reaches into new fresh culture medium.
2. For simplicity, we assumed spherical cells. This is unlikely to affect the calculations significantly given the aspect ratio of *Tetrahymena*.
3. The assumption which certainly might have the biggest consequences is that oxygen is ‘consumed’ at the cell surface: we set *C_o_*to take a fixed value at the position of the cell membrane. We could speculate that the diffusion of oxygen across the cell membrane and to the mitochondria is particularly inefficient and for this reason a very high concentration *C_o_* is required (though in terms of diffusion across cellular membranes at least, this does not seem likely, see Subczynski et al. (1989)).

This situation – in which oxygen is utilised uniformly everywhere inside the cell – can be modelled by considering that, inside the cell, both diffusion and oxygen consumption affect the availability of oxygen to the most internal parts of the cell. As we don’t focus on oxygen in our study, we leave a detailed modelling of oxygen supply and consumption inside *Tetrahymena* cells to a future study, but we just mention an analogous system, which was modelled by Grimes et al. (2014). These authors considered a model, in which the oxygen concentration at the cell membrane (though their model was not originally developed for cells, but for tumour spheroids) is essentially *C*_∞_, and oxygen reaches the inner core of the cell (or tumour spheroid) if

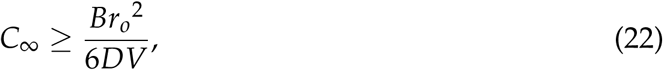

where *C*_∞_ is the concentration of oxygen at the cell membrane, *r_o_* is the cell radius, *D* is the diffusion coefficient, *V* is the cell volume, and *B* is the metabolic rate (expressed in units of molar oxygen consumption in order to match the units of oxygen concentration *C*_∞_). Rearranging equation 22 we get that anoxic regions inside the cell are avoided if the cell radius 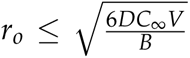. We take 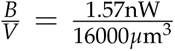, or 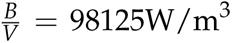, as the ratio between metabolic rate at 30 °C and cell volume, roughly matching the experimental data). The term 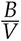 represents a measure of metabolic rate per unit of biomass inside the cell, and can be converted to units of molar oxygen depletion by assuming the equivalence 1 mol O_2_ = 478576 J, giving 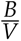 = 0.20 mol O_2_/m^3^/s. Hence, by this model, the maximum cell radius before cell size impacts diffusion-driven oxygen supply for a cell at 30 °C would be 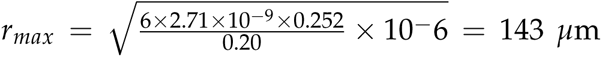. In a sense, then, also this alternative set of arguments seems to suggest that oxygen availability only becomes a limiting factor for cells or organisms substantially larger than *Tetrahymena* when these are tested in our growth conditions.

All these calculations on whether oxygen becomes a limiting factor for *Tetrahymena* growth in our experiments are based on crude approximations, and do not provide sufficiently strong evidence in favour of conclusion or another. However, the fact that oxygen was not a direct limiting factor is also supported by empirical evidence. This includes the observation that *Tetrahymena* population growth remained in the exponential growth phase also for densities much higher than those used in our experiments (presented above in section 1.2). At higher population densities, oxygen is more likely to become partially depleted from the culture medium, and yet growth was not affected. Oxygen consumption in the Sensor Dish Reader also remained stable throughout our measurements, with oxygen decreasing linearly in time also after it had reached low values below 100*µM*, showing no evidence of altered metabolism within a wide range of oxygen concentrations. Finally, in our preliminary experiments in which we altered oxygen concentration directly (SI section 2.9) we did not observe changes of cell size or shape, also suggesting that the maximum size attained by *Tetrahymena* remained stable also when oxygen concentration in the culture medium was halved.

All these different pieces of evidence seem to exclude the possibility of direct effects of oxygen availability on *Tetrahymena* metabolism and population growth in our experimental conditions.

But, if in our experimental conditions of axenic cultures kept at low population densities in well-aerated culture flasks, *Tetrahymena* probably never encountered growth-limiting concentrations of oxygen, we should note that our culture conditions are far from being representative of the conditions that *Tetrahymena* is likely to experience in its natural environment, such as in a pond, and even of conditions experienced in laboratory cultures, as soon as cultures are left to grow and reach high cell densities beyond the exponential growth phase. Our *Tetrahymena* cells might have evolved reaction norms that decrease body size when temperature increases, due to previous selection when growing conditions with respect to oxygen were less favourable (reviewed in Verberk et al. (2021)). In this case, some of the phenotypic changes – such as the changes in cell volume – that we observe when we expose *Tetrahymena* to a different temperature would represent adaptive norms of reaction that provide a selective advantage for coping with limiting oxygen supply/demand ratios, except that these phenotypic changes would not be directly beneficial in our culture conditions, and they would be triggered not by oxygen directly, but by other environmental cues (for instance, by temperature itself). This would be the reason why we observe them also in our experimental conditions in which oxygen, most likely, was not limiting.

To conclude this section, we should also mention one final observation: within the range of parameters that we used in our calculations the curves that describe the ratio between supply and demand Φ (fig. 30) are almost entirely determined by changes in oxygen demand (through the temperature- and body-size-dependence of metabolic rate). The modelled oxygen supply to the cells only showed little variation across the entire temperature range (figure 31(c), calculated on the basis of eq. 20). Oxygen supply is little affected by temperature changes because while an increasing temperature reduces the solubility of oxygen (decreasing *C*_∞_, 31(a); see below, section 3.4 for details of the calculations), it also increases the diffusion coefficient *D* that also contributes to oxygen supply in eq. 20 (fig. 31(b)). The estimated rates of oxygen supply also depend on the parameter *C_o_* in eq. 20, but it remains generally true that oxygen demand, rather than supply, is likely to be the main determinant of the supply/demand ratio (see also Verberk et al. (2011)).

**Figure 31:**
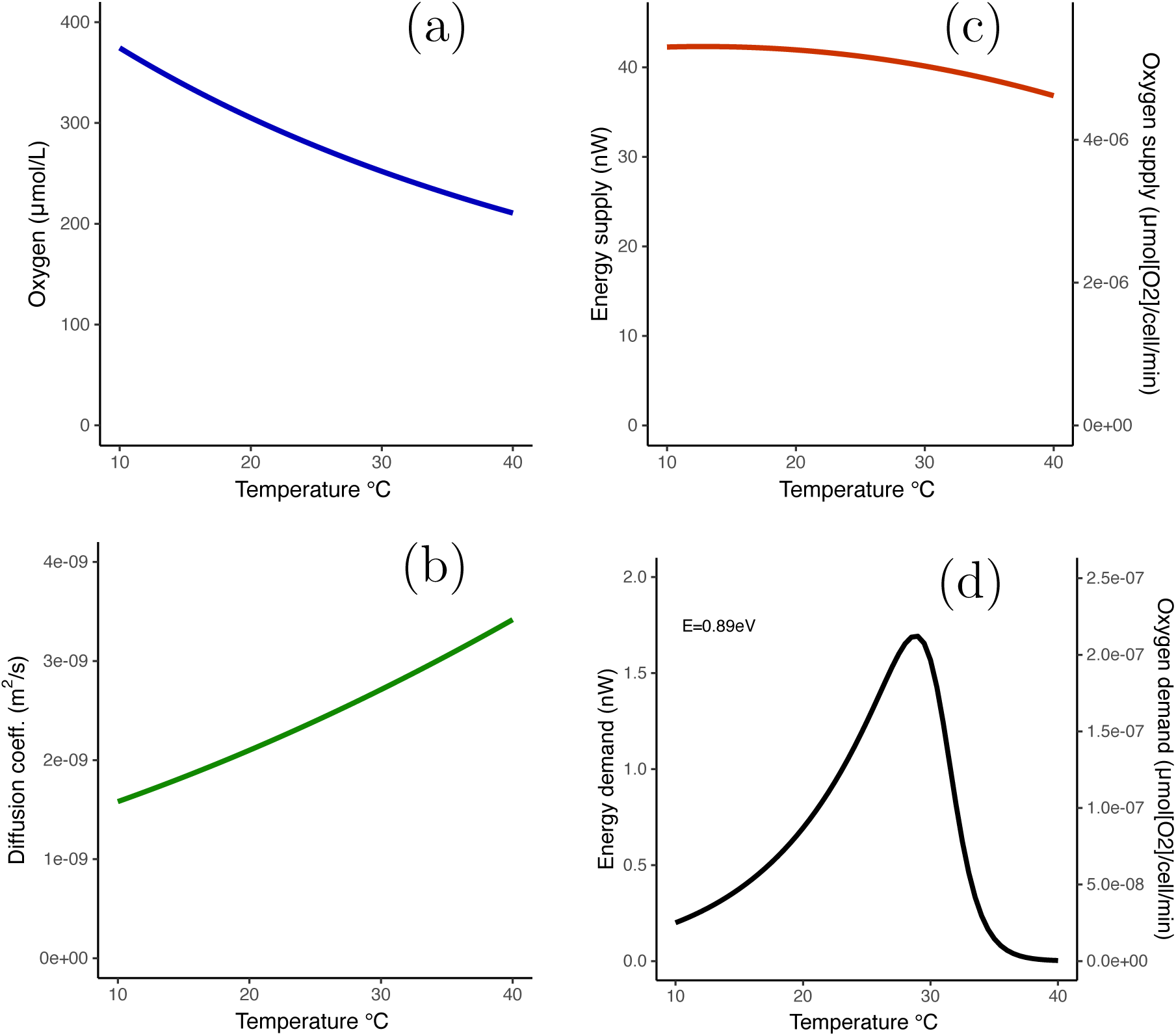
Effects of temperature on oxygen concentration and diffusion. **(a)** Effects of temperature on oxygen concentration at fully saturated equilibrium. **(b)** Effects of temperature on the diffusion coefficient of oxygen in water. **(c)** Estimated oxygen supply to a spherical cell of volume 10^4.5^*µ*m^3^ assuming that the cell absorbs oxygen up to a concentration 100 *µ*M on the cell surface. **(d)** For comparison, estimated energy and oxygen demand for a cell with the same volume (10^4.5^*µ*m^3^) based on plausible parameter values obtained experimentally from our experiments.

The importance of metabolic demand, relative to supply, limits the explanatory power of the type of supply/demand curves presented in figure 30, because supply-demand curves for other nutrients and resources different from oxygen, or for the removal of catabolic products would likely be similar, as long as their shape is mostly determined by the cellular metabolism rather than by the specific nature of the molecules considered. Experiments in which oxygen concentrations are directly manipulated would be required in order to fully disentangle the role of oxygen availability from other factors.

#### 3.3 Diffusion around a sphere

For the convenience of our readers, here we provide a derivation of equation 20, that we used above to model the diffusive supply of oxygen to an individual cell. The problem considered is that of a small spherical unicellular organism of radius *r* that lives in a non-turbulent medium and obtains oxygen through diffusion. The concentration of oxygen at an infinite distance from the cell is *C*_∞_, while the concentration on the surface of the cell is *C_o_*. The extreme when *C*_0_ = 0 corresponds to the assumption of a perfectly absorbing cell surface (any molecule that touches the surface of the cell is immediately taken). Based on these boundary conditions for the diffusion equation, we can estimate the gradient of oxygen concentration around the cell, and the total flow or current of oxygen into the cell per unit time. This topic is covered for instance in the book by Berg Berg (1993).

We start by writing the diffusion equation in spherical coordinates, assuming radial symmetry (i.e. neglecting cell elongation for the purpose of our approximate calculations):

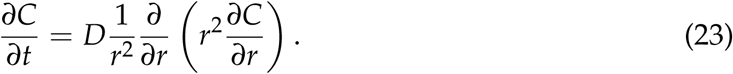

At the steady state 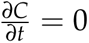. This means that we must have:

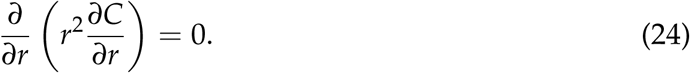

If the left side is equal to zero, then its integral with respect to *r* is equal to a constant that we indicate here as *K*_1_:

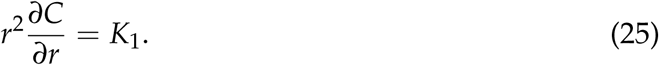

We can further rearrange and integrate to get

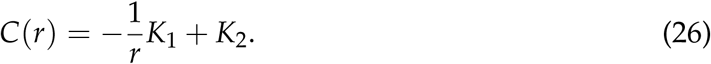

Now we can get the values of *K*_1_ and *K*_2_ from the boundary conditions. At *r* = ∞ we have *C*(*r*) = *K*_2_ = *C*_∞_. At *r* = *r_o_* we have *C*(*r_o_*) = *C_o_*.

This leads to the following expression when *C_o_* = 0:

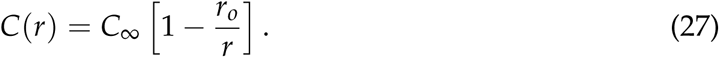

For C_o_ ≠ 0 we have instead

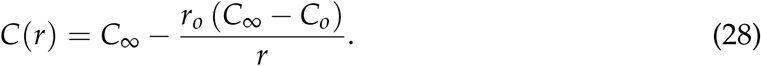

One important observation from eq. 28 is that the concentration at any given point does not depend on the diffusion coefficient, but only on the radius of the cell and on the two concentrations *C_o_*and *C*_∞_.

Now that we know the gradient, we can calculate the flux if we remember that

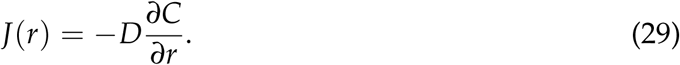

In fact, we obtain

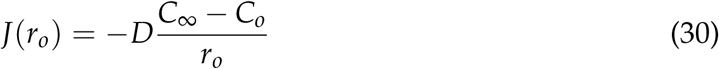

for the flux at the position of the cell membrane. In this case the flux does depend on the diffusion coefficient. In general the relevant quantity is not the flux at the position of the cell membrane, but the total rate of nutrients arriving to the cell, which is flux times area. Let’s call this the current or the flow of oxygen into the cell, and indicate it with the letter *I*. This is easily calculated if we remember that the surface area of a sphere is 4*πr*^2^_o_:

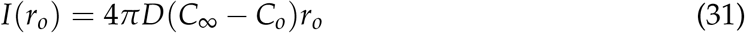

The main result of these calculations is that the current into the cell scales with the radius of the cell. This means that for large cells it might becomes soon insufficient to supply enough molecules (oxygen, nutrients) to the entire cell volume (which scales instead as *r*^3^_o_).

#### 3.4 Temperature dependence of oxygen concentration in fully saturated water

At equilibrium, the amount of oxygen (or another gas) dissolved in water is proportional to the partial pressure of the gas above the water. The proportionality constant is called Henry solubility constant. For oxygen at 25 °C we have 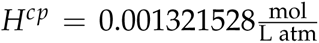. This means for instance that at one atmosphere, with a partial pressure of oxygen of 0.20948 atmospheres we have a concentration in water of 277*µ*mol/L or 8.86mg/L. In practice, the presence of solutes in water also affects the solubility of oxygen so that we should have a different constant for each level of salinity.

When the temperature increases or decreases, the Henry solubility constant also changes and this change is well described by Van’t Hoff equation:

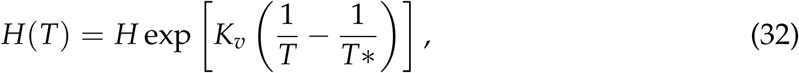

where

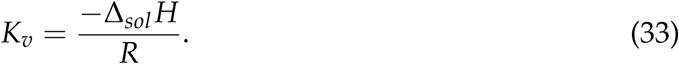

As a worked example, let’s calculate the concentration of oxygen at equilibrium at a temperature of 30 degrees Celsius.

The partial pressure of oxygen at one atmosphere is *p_O_*_2_ = 0.21*atm*. In order to know how much of this oxygen is dissolved in water we need to know the Henry solubility constant. The Henry solubility constant at 25 °C (298.15 K) is *H^cp^* = 0.00132mol/L/atm. The Van’t Hoff parameter *K_v_*for *O*_2_ is 1700. We can then calculate the Henry solubility constant at 30 °C (303.15 K) as

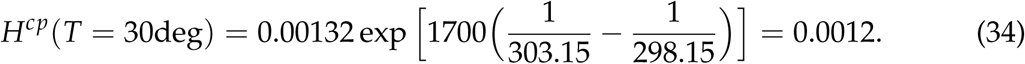

Oxygen concentration is then calculated by multiplying [*O*_2_] = 0.21 × 0.0012 mol/L, that is, [*O*_2_] = 252 *µ*M. The conversion to mg/L is easily obtained by considering the molecular weight of MW(*O*_2_) = 32 g/mol: 252*µ*mol/L × 10^−3^ mmol/*µ*mol × 32 mg/mmol = 8 mg/L.

#### 3.5 Diffusion of molecules in water

The diffusion coefficient in liquids is described by the Stokes-Einstein equation. In the special case of spherical particles moving through a liquid with low Reynolds number we have

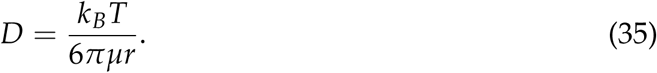

Where *D* is the diffusion coefficient, *k_B_* is the Boltzmann constant, *T* is the absolute temperature, *µ* is the dynamic viscosity of water and *r* is the radius of the particle. Note that the denominator is similar to the drag coefficient for a sphere.

Given this dependence on temperature we have that

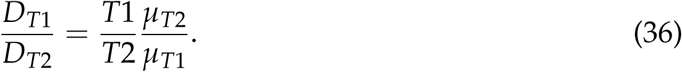

At 20 degrees Celsius, the diffusion coefficient of oxygen in water is about 2.10 × 10^−9^m^2^/s; the diffusion coefficient of CO_2_ in water is 1.77 × 10^−9^m^2^/s. The diffusion coefficient of many other small molecules is in the same order of magnitude.

We can then estimate the diffusion coefficient for oxygen in water at 30 °C as 2.71 × 10^−9^m^2^/s

#### 3.6 Implications of cell shape for swimming

In our experiments, cells increased size mainly through becoming more elongated, which mainly happened at low temperature and high food concentrations (see fig. 11). One might ask about the possible implications of different cell shapes for swimming. For example, what shape a cell should have in order to minimise the drag when swimming at low Reynolds numbers. This problem was addressed, for example, by Cooper and Denny Cooper and Denny (1997), who calculated the passive drag of a prolate spheroid moving in a viscous medium. The spheroid had constant volume and variable elongation, so they could analytically find the value of elongation that minimised the drag. As the approximated equation used in the original article is only valid for extremely elongated spheroids and would lead to incorrect predictions, we re-derive it here.

Consider a prolate spheroid with a volume *V*:

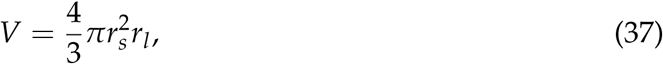

where *V* is fixed, and the only thing that can vary are the small and large radius *r_s_* and *r_l_*, under the constraint of maintaining the same cell volume. We can define the elongation *ψ* of the spheroid as 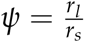. Once we have done this, we can rewrite the formula for the volume as

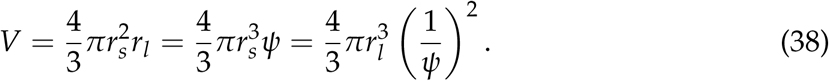

From this, we get an equation for *r_s_* as a function of volume and elongation

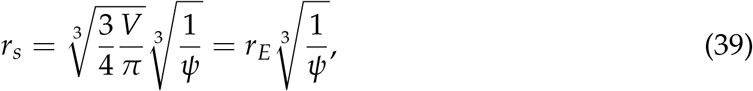

where *r_E_*is an equivalent radius (the radius of a sphere with the same volume).

If we approximate *Tetrahymena* as a prolate spheroid moving along its long axis, there are multiple equivalent formulas that allow calculating the drag coefficient – e.g. Katsu-Kimura et al. (2009); Clift et al. (2005). one such exact formula for the drag coefficient of a spheroid is:

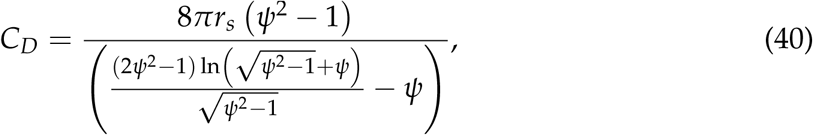

where *ψ* is the elongation of the spheroid (ratio of the long radius *r_l_* over the short radius *r_s_*). Equation 40 is well approximated by the following approximate formula:

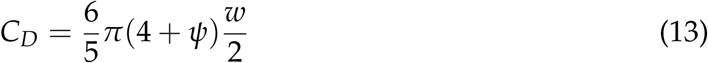

(Clift et al., 2005).

Remembering this approximate formula, it is easy to replace the small radius 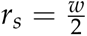 with the expression that we had above in terms of equivalent radius *r_E_* and elongation *ψ*, to obtain:

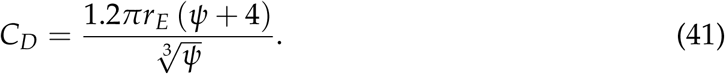

We then derive with respect to the elongation *ψ*, and we get

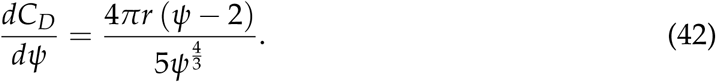

The derivative indicates that the minimum drag is for an optimal value of elongation *ψ*^∗^ = 2. Using the exact equation for the drag coefficient (40) rather than the approximate one does not make a huge difference, giving an optimal elongation *ψ*^∗^ ≈ 1.95. One important take-home message is that the shape that minimizes drag based on these calculations always has an aspect ratio of approximately 2:1 (length:width), which should not depend on characteristics of the growth medium, temperature, or the typical movement speed of the cells. We should however remember that these calculations are based on formulas for the passive drag, while *Tetrahymena* are active swimmers. It is possible that the more elongated shape of *Tetrahymena* cells cultured at low temperature might allow for a more powerful, albeit less efficient, swimming, due to an increase in the number of cilia oriented perpendicularly relative to the direction of movement, or to other similar factors (see sec. 6.1 – 6.2 in Velho Rodrigues et al., 2021, for some possible models). Alternatively, the change of elongation which we observe depending on temperature should be attributed to other internal constraints on cell size and shape, rather than to adaptive phenotypic responses related to the efficiency and power of swimming.

## Footnotes

1 According to Brown et al. (2018), the energy content of many organisms is 22.4 kJ/g of dry body weight. In addition, Gates et al. 1982 Gates et al. (1982) state that the ratio of dry to wet weight for *Tetrahymena* (of a different species) is 59%. Using these two numbers and assuming a density of *Tetrahymena* similar to that of water (1 *µ*m^3^ of *Tetrahymena* = 10^−12^ g) we get 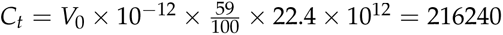 nJ.

2 Katsu-Kimura et al. 2009 Katsu-Kimura et al. (2009) report an efficiency of 0.00078 in Paramecium. We do not have a measure of *η_c_* for *Tetrahymena*, but we can estimate its approximate value relative to respiration energy consumption (which likely encompass both energy expended for movement and basal metabolic rate) based on the typical values of relevant quantities that we measured in our experiments, e.g. by considering a respiration rate of ∼ 0.7 nW, of which about 0.3 nW are allocated to movement, a movement speed of ∼ 500 *µ*m/s, a small radius *r_s_* = 12.5 *µ*m and an elongation *r_l_*/*r_s_* = 2, and a viscosity *µ* = 0.001 kg/(m s), leading to a value of *η_c_* as small as ∼ 0.02%. An efficiency of 0.00025 corresponds to a power expenditure for movement of ≈ 0.3 nW.

